# Impacts of human-introduced species on the geography of life on Earth

**DOI:** 10.1101/2025.07.05.663167

**Authors:** César Capinha, Rebecca Pabst, Boris Leroy

**Affiliations:** Centre of Geographical Studies, Institute of Geography and Spatial Planning, University of Lisbon, Lisbon, R. Branca Edmée Marques, 1600-276 Lisboa, Portugal; Associate Laboratory Terra, Portugal; Global Health and Tropical Medicine, GHTM, LA-REAL, Institute of Hygiene and Tropical Medicine, IHMT, NOVA University Lisbon, Lisbon, Portugal; Laboratoire de Biologie des Organismes et des Écosystèmes Aquatiques-BOREA, Muséum national d’Histoire naturelle (MNHN), SU, CNRS, IRD, UA, 43 rue Cuvier, F-75005 Paris, France

**Keywords:** Anthropocene, Biodiversity redistribution, Biogeography, Biological invasions, Global change, Human-mediated dispersal, Macroecology, Novel ecosystems, Species ranges

## Abstract

Human activities are increasingly transporting species beyond their native ranges, where they often establish and become permanent additions to recipient biotas. These introductions, documented for tens of thousands of taxa, are a departure from the natural constraints of dispersal that have long shaped species distributions. However, current knowledge on their impacts on global biogeography remains poorly understood and fragmented. Here, we review empirical evidence across 15 classical biogeographical rules to assess how non-native species are altering the spatial structure of global biodiversity. Our synthesis reveals that while some patterns, such as species-area relationships, are often reinforced under invasion, others, including the delineation of global biogeographic regions, relationships between island isolation-diversity and between body sizes and latitudinal gradient (i.e., Bergman’s rule), already show profound reshaping in contemporary assemblages. Additionally, other patterns (e.g., latitudinal gradient of species diversity; body size and insularity) show context-dependent changes, shaped by factors such as spatial scale, taxonomic group, and introduction history. These transformations are often more pronounced at broad spatial scales and in highly invaded systems such as islands and temperate regions. Our findings demonstrate that biological invasions are selectively but profoundly reshaping the geography of life on Earth, with major implications for conservation, macroecology, and the future of biodiversity patterns in the Anthropocene.

## I. INTRODUCTION

Human activity is fundamentally changing global biodiversity, with biological invasions (defined as the human-mediated introduction and establishment of species outside their native ranges) emerging as one of the most pervasive and irreversible drivers of ecological change. As of recent estimates, nearly 40,000 non-native species have become established globally (Roy *et al*., 2023), forming permanent components of recipient communities. These invasions frequently result in substantial ecological impacts, including biodiversity loss, disruption of ecosystem functions, and reductions in human well-being (Doherty *et al*., 2016; Wainright *et al*., 2021; Roy *et al*., 2023; Carneiro *et al*., 2025).

Biological invasions are, at their core, a biogeographical process, representing a departure from the limitations of natural dispersal and evolutionary history that have structured species distributions over deep time. Unlike long-term geological forces or contemporary drivers such as anthropogenic climate change (Lenoir *et al*., 2020), invasions enable the rapid movement of taxa across vast areas and over formerly insurmountable biogeographic barriers, including oceans, deserts, and polar regions (Tingley *et al*., 2015; Ficetola, Mazel & Thuiller, 2017; Williams, Zipkin & Brodie, 2025). The prevalence of non-native species is now high across many regions (Dawson *et al*., 2017), in some cases exceeding native diversity (Leprieur *et al*., 2008; Clements, Catania & Searcy, 2019; Essl *et al*., 2019), and the pace of introductions continues to accelerate due to globalization of trade and transport networks (Seebens *et al*., 2017; Seebens *et al*., 2021; Capinha *et al*., 2023). These human-mediated biotic flows are also globally asymmetrical, with some regions functioning primarily as sources and others as sinks (Capinha *et al*., 2023; Leroy *et al*., 2023; Bertelsmeier *et al*., 2025), and they encompass a broad taxonomic and functional spectrum, from microorganisms to some of the largest plants and animals on Earth (Seebens *et al*., 2017; Briski *et al*., 2024). Collectively, these dynamics suggest that the reshaping of Earth’s biota via biological invasions is ongoing and intensifying while also increasingly decoupled from the patterns shaped by natural dispersal and evolutionary processes.

While substantial research has focused on understanding the ecological effects of non-native species (e.g., Simberloff *et al*., 2013; Bacher *et al*., 2025), the broad consequences of invasions for global biogeography and macroecology are poorly known because they were never synthesized. There are many broad-scale patterns of biodiversity that are generalized as biogeographical and macroecological patterns, often termed ‘rules’ (Brown, 1995; Gaston & Blackburn, 2000; Diniz-Filho, 2023). These key patterns are likely altered and reshaped by invasions, as reported in the growing body of observational evidence ranging from species-area relationships (e.g., Guo *et al*., 2021a) and diversity gradients (e.g., Blanchet *et al*., 2010; Dawson *et al*., 2017; Moser *et al*., 2018) to the delineation of biogeographic regions (e.g., Leroy *et al*., 2023; Aulus-Giacosa *et al*., 2024). Yet, the existing literature is fragmented, concentrating on specific patterns (e.g., Guo *et al*., 2021b), taxa (e.g., Van der Geer *et al*., 2018; Dyer *et al*., 2020), or geographic regions (e.g., Helmus, Mahler & Losos, 2014; Fristoe *et al*., 2021), which limits our ability to assess their cumulative and cross-scale effects.

Here, we synthesize available evidence on how biological invasions are reshaping the geography of life on Earth by evaluating the impacts of non-native species across 15 foundational biogeographical and macroecological patterns and rules (hereafter collectively referred to as “rules” for simplicity) (Gaston & Blackburn, 2000; Lomolino, Riddle & Whittaker, 2017; Diniz-Filho, 2023). These include classic spatial gradients (e.g., latitude-diversity, bioregions delineation, isolation-diversity), structural relationships (e.g., species-area and distance decay of similarity), and trait-based rules (e.g., Bergmann’s and Gloger’s). Our synthesis focuses on findings drawn from a systematic review of observational studies across taxa and regions (see Methods). We examine both the spatial patterns identified for non-native assemblages or individual species, and their additive or disruptive effects on native and contemporary assemblages (see Glossary in Table 1). The section that follows synthesizes this evidence by relationship and effect types (Fig. 1), spatial scale and taxonomic unit, offering a global perspective on how biological invasions are modifying long-standing patterns of biodiversity distribution and identifying key priorities for future research.

**Fig. 1.**
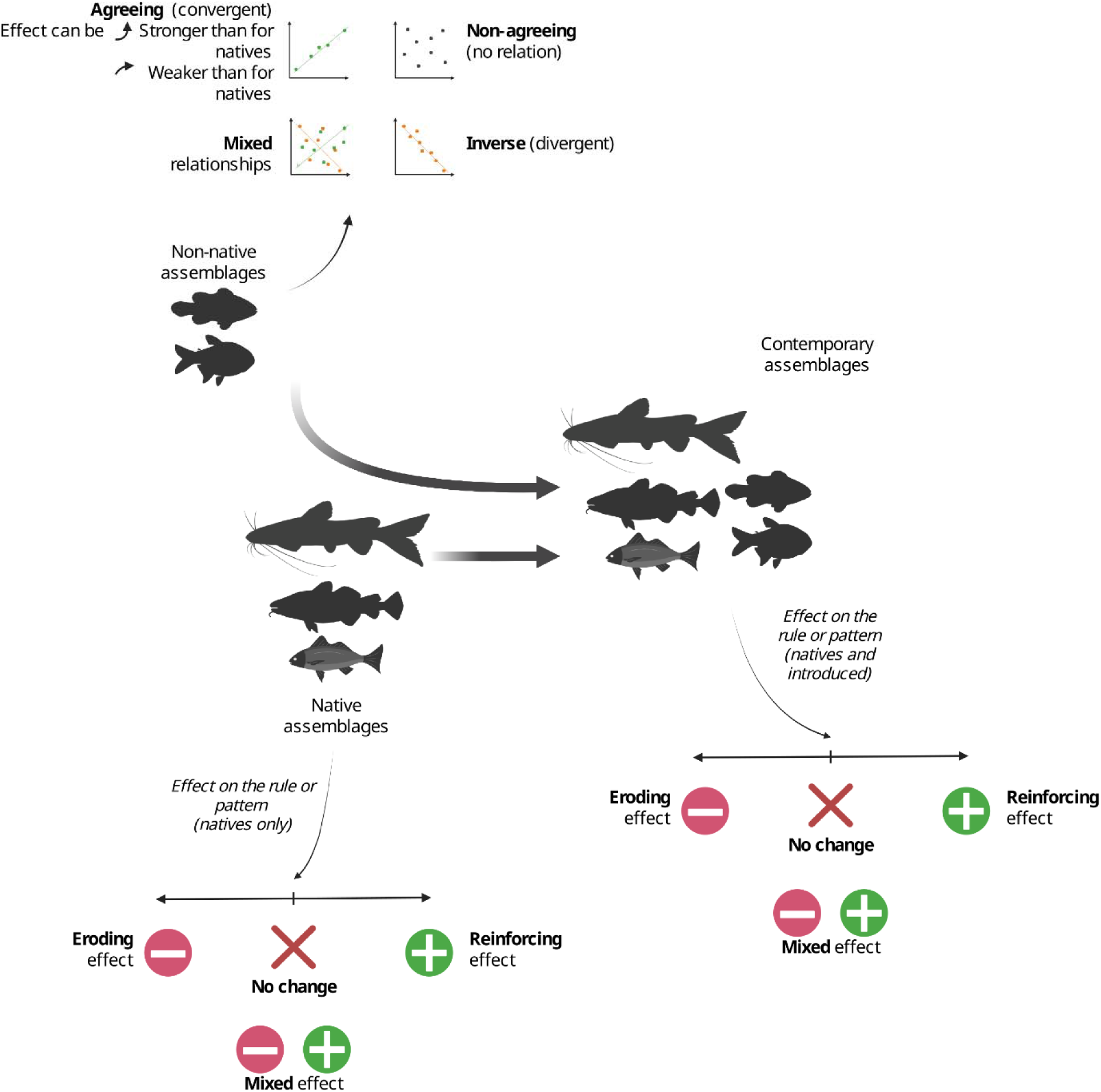
Main relationship types, effect categories, and biological units of analysis identified in this study. We distinguish among three main units of analysis: 1) non-native assemblages or individual non-native taxa; 2) native assemblages or individual native taxa, and 3) contemporary assemblages (i.e., current communities including both native and non-native species). For non-native taxa, we recorded whether reported patterns: (i) aligned with the theoretical expectation of the rule, (ii) showed an inverse relationship, or (iii) exhibited other non-agreeing patterns. For native and contemporary assemblages, we classified non-native species’ effects into three categories: reshaping or eroding effects, indicating a weakening or deviation from expected patterns; reinforcing effects, indicating a strengthening of expected patterns; and no change, when no clear effect was observed. A fourth category, ‘mixed’ effects, was used when studies reported variable outcomes across different contexts (e.g., distinct effects in different regions). For a few rules, relationships or effect categories additional descriptors were included to better describe the nature of reported patterns (e.g., using relative comparisons such as “weaker” or “stronger” in distance decay patterns).

**Table 1.**
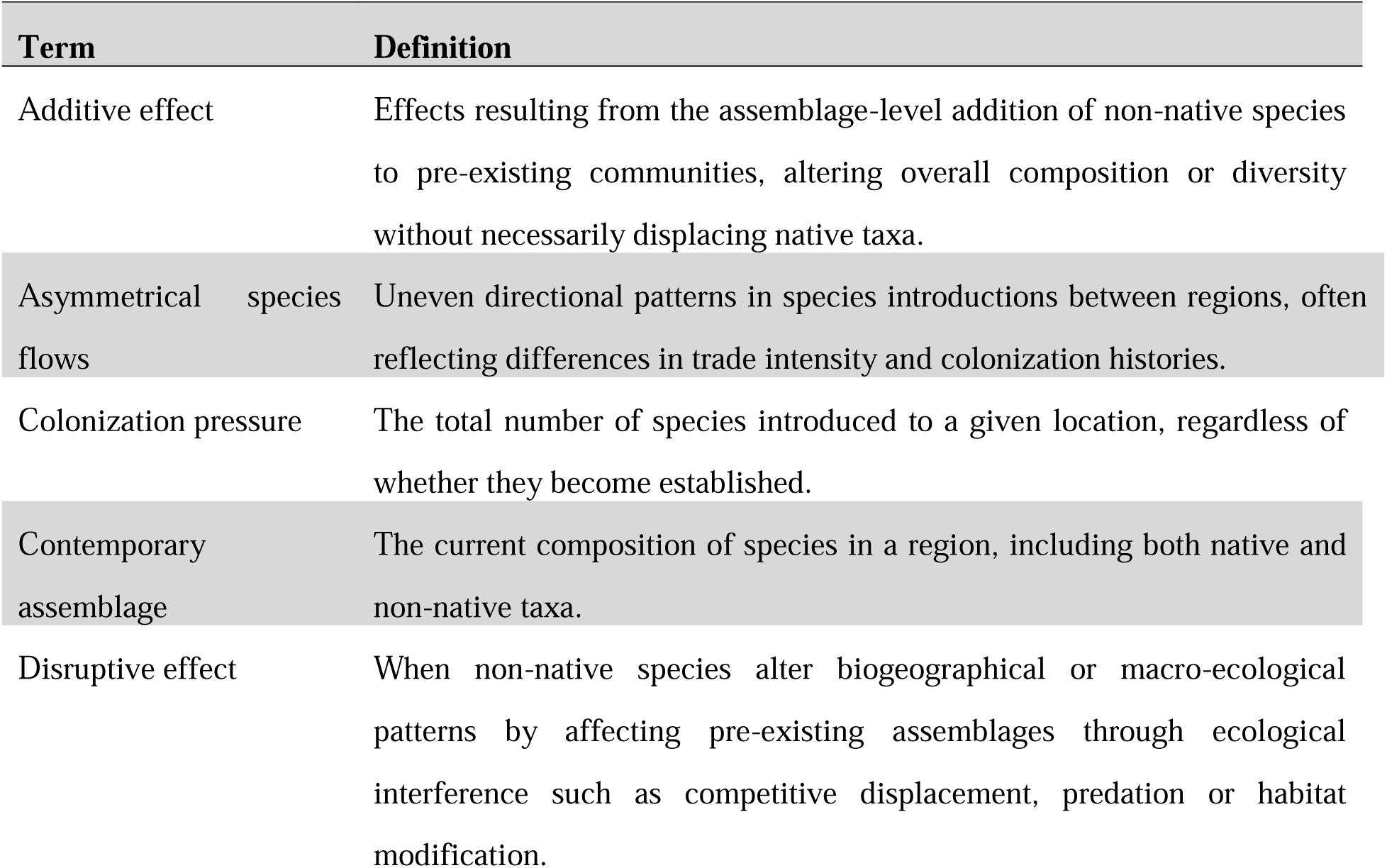
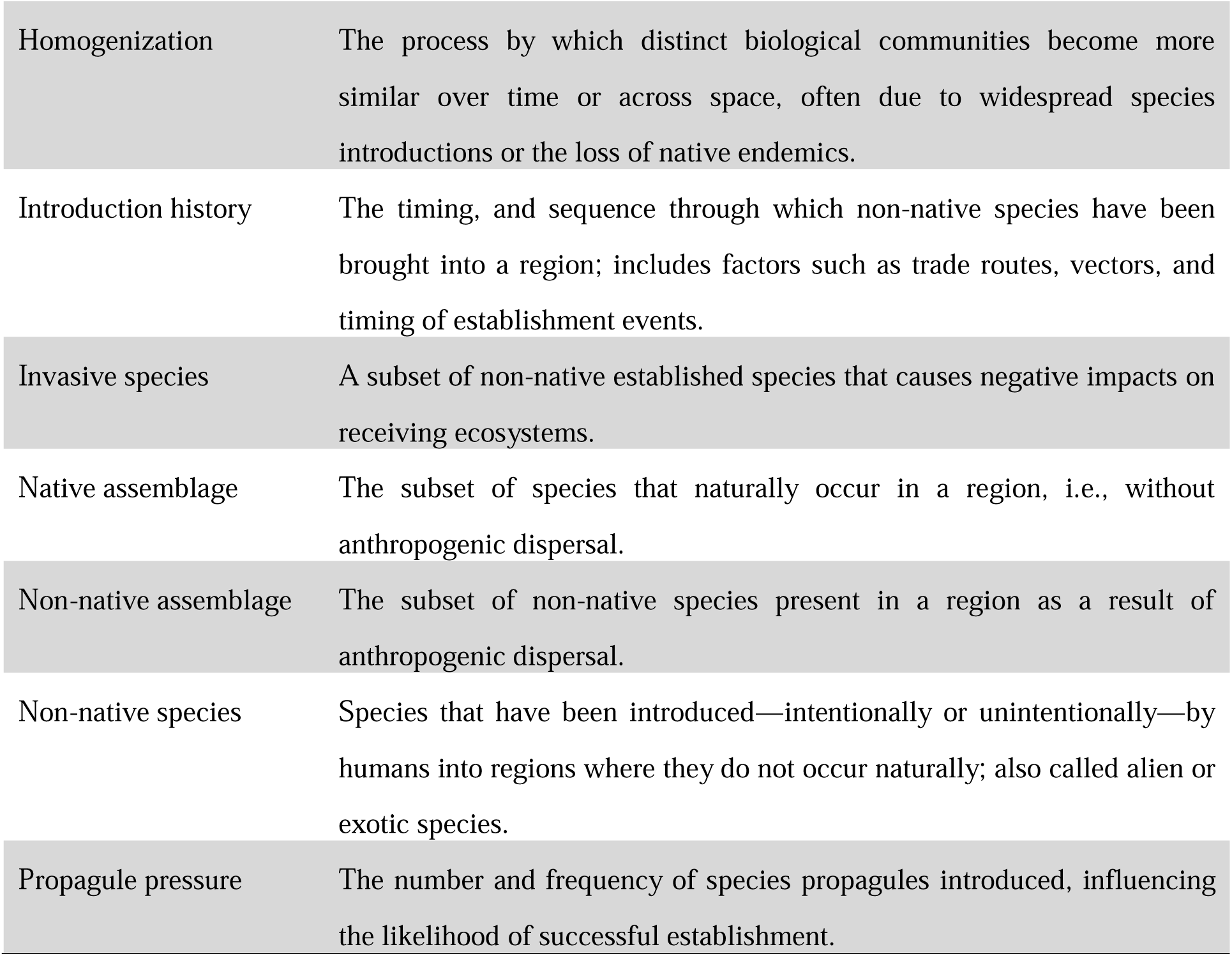
Glossary of main terms.

## II. IMPACTS OF BIOLOGICAL INVASIONS ON BIOGEOGRAPHICAL AND MACOECOLOGICAL PATTERNS AND RULES

### (1) Latitudinal gradient in species diversity

The latitudinal gradient in species diversity (LGSD) is one of the most well-established patterns in biogeography, with species richness peaking in the tropics and a decline toward the poles. This gradient is consistently observed in native taxa across a wide range of ecosystems (Gaston, 2000). Our review shows that, although the LGSD has been one of the most frequently studied biogeographical patterns in relation to non-native taxa (Fig. 2), few studies have directly assessed how these species influence patterns in native or contemporary assemblages (Fig. 3a). Overall, at the largest scales (i.e., continental to global), non-native assemblages rarely align with the expected LGSD, while at narrower scales (i.e., regional to local), the results are more variable (Fig. 3a).

**Fig. 2.**
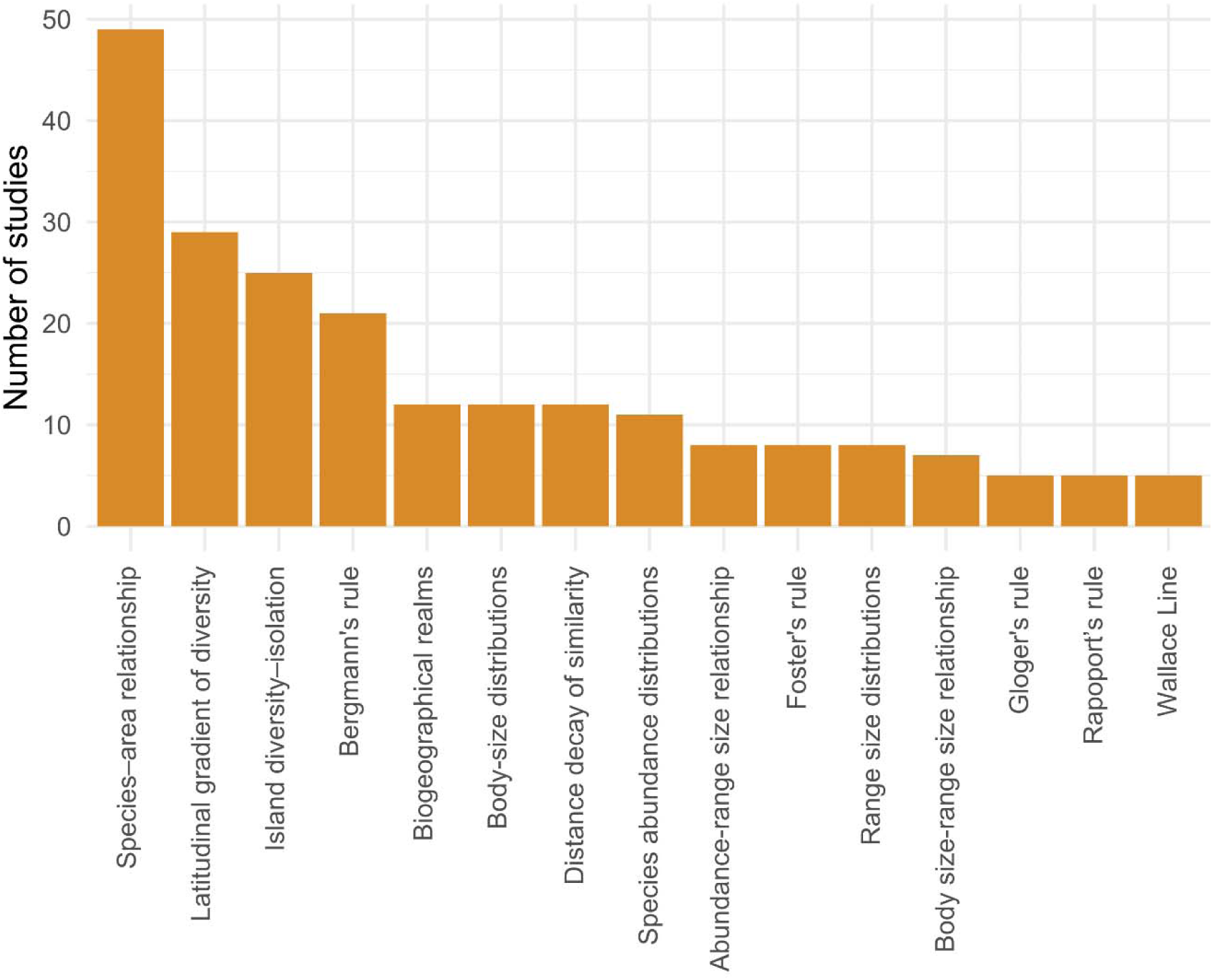
Number of identified studies examining the spatial patterns or effects of non-native taxa for each of the 16 biogeographical patterns or ecogeographical rules assessed.

**Fig. 3.**
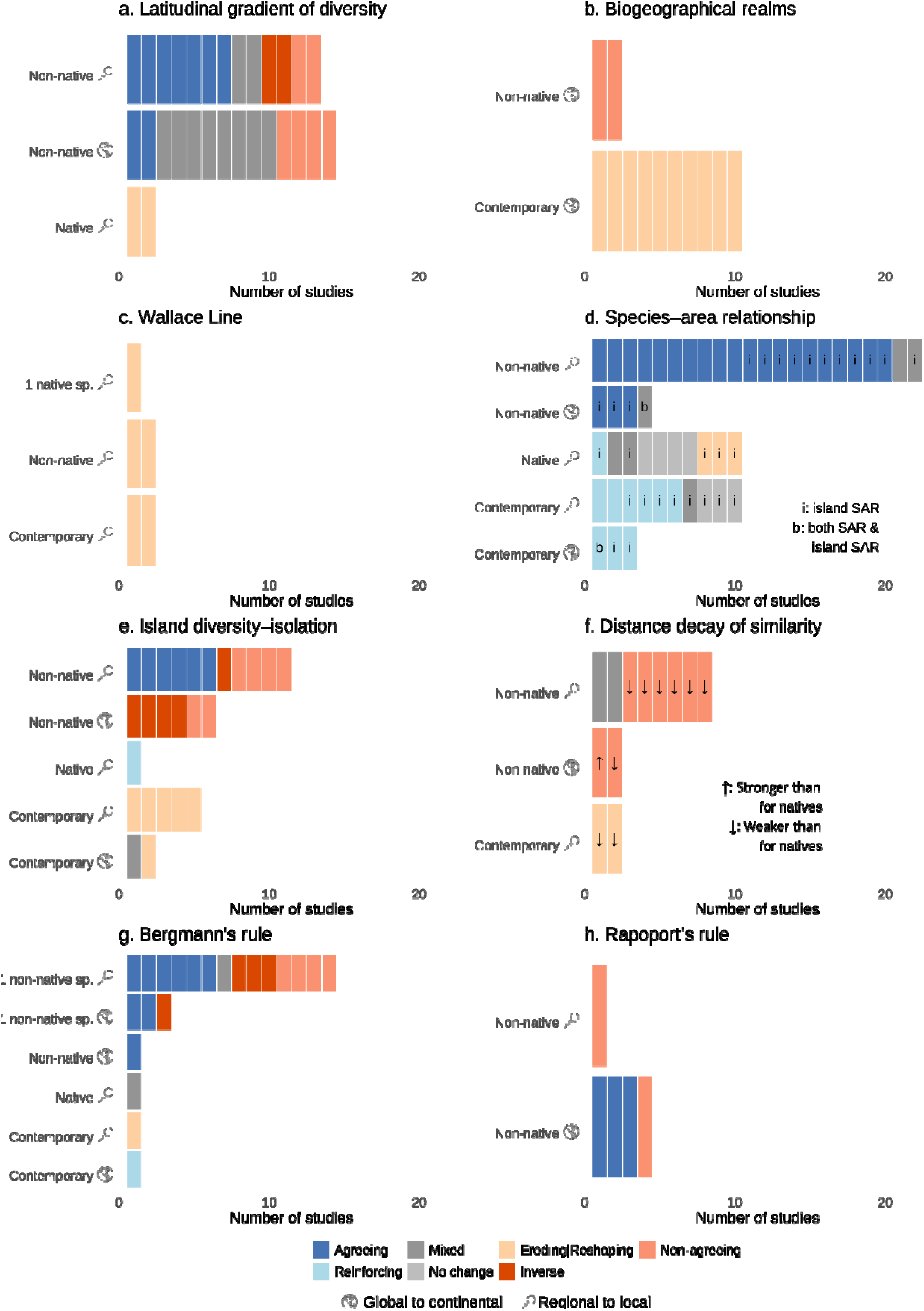
Synthesis of studies assessing the patterns or effects of non-native species in relation to (a) the latitudinal gradient in biodiversity, (b) the delineation of global biogeographical regions, (c) Wallace Line, (d) (island) species-area relationship, (e) the relationship between island isolation and species diversity, (f) distance decay of compositional similarity, (g) Bergmann’s rule and (h) Rapoport’s rule. Plots show the number of studies (horizontal axis) per rule. Colours indicate the nature of agreement with theoretical expectations. Blue tones in the right indicate results that align with the expected biogeographical pattern or rule, orange and red tones indicate deviations. Grey tones denote mixed results or no observed effect.

At global and continental scales, 86% of studies (12 out of 14) report that non-native species richness deviates from the expected latitudinal gradient. Non-native species richness often showed no consistent relationship with latitude. A recurring finding is a peak in species richness at temperate latitudes, rather than the tropics, resulting in a hump-shaped distribution. This pattern has been documented for non-native plants (e.g., McKinney, 2006; Pyšek & Richardson, 2006; Sax 2008; Guo *et al*., 2021a), birds (Sax, 2008; Dyer *et al*., 2017, 2020), mammals and fishes (Sax, 2008). Nonetheless, while a peak in species richness at temperate latitudes has been consistently reported, multiple groups (herpetofauna, ants, plants, birds, mammals and fishes) also exhibited a decrease in richness with increasing latitude within temperature regions (e.g., McKinney, 2006; Pysek & Richardson, 2006; Schnitzler, Hale &Alsum, 2007, Sax, 2008; Dyer *et al*., 2020; Guo *et al*., 2021a).

At regional to local scales, patterns are extremely variable. Roughly half of the studies reviewed (n=7) report a decline in richness with increasing latitude, consistent with the LGSD (e.g., Habit, 2012; Mologni *et al*., 2021; Ragkousis *et al*., 2023). Several studies (n=6) also reveal non-agreeing patterns, including inverse relationships (i.e., where richness increases with latitude) (e.g., Pyšek, 1998; Celesti-Grapow *et al*., 2010). Notably, a few (n=2) also identify cases where dominant invasive species disrupt latitudinal patterns in native assemblages. For example, the plants *Spartina alterniflora* in Chinese salt marshes (Zhang *et al*., 2019) and *Alternanthera philoxeroides* in subtropical Asia (Gao *et al*., 2021) have been shown to suppress native richness and erode latitudinal signals.

Overall, at all spatial scales, the introduction of non-native species has led to marked deviations from the natural LGSD patterns. Natural LGSD patterns were driven by a combination of multiple non-anthropogenic drivers (i.e., abiotic, biotic and dispersal limitations), diversification rates and time for species accumulation (Pontarp *et al*., 2019). The observed deviations from the natural LGSD suggest that now, anthropogenic drivers also play a major role in shaping spatial patterns of diversity. Indeed, anthropogenic drivers as uneven colonization pressure (i.e., variation in the number of species introduced; Lockwood, Cassey & Blackburn, 2009), often outweigh climatic constraints in explaining invasion dynamics (Dawson *et al*., 2017). Temperate regions, in particular, tend to harbour greater non-native richness due to longer histories of introduction and climatic similarity with source regions (Pyšek & Richardson, 2006; Dawson *et al*., 2017). However, despite substantial deviations, the multiple reports of a decrease in non-native species richness with increasing latitude suggest that natural drivers continue to exert influence on spatial patterns of diversity, even within non-native assemblages. These spatially variable anthropogenic drivers, which interact across latitudinal and spatial scales, will likely result in continued deviations from the natural LGSD in the near future.

### (2) Biogeographical regions of the world

Biogeographical regions are foundational units of biogeography (de Candolle, 1820; Wallace, 1876). These regions historically hosted distinct species assemblages, shaped by shared evolutionary history and limited species exchange due to major natural barriers (Ficetola *et al*., 2017). Our synthesis (Fig. 2; n=12), reveals strong evidence that human-mediated species introductions are profoundly reshaping these long-standing boundaries.

Most studies (n=10), spanning a range of taxonomic groups, reported that non-native species are redrawing global bioregion boundaries through additive effects (Fig. 3b), confirming the hypothesis of a “New Pangaea” proposed by Rozensweig (2001). For freshwater fishes, Leroy *et al*. (2023) documented the emergence of a novel biogeographical region, PAGNEA (Pan-Anthropocenian Global North and East Asia), resulting from extensive introductions across North America, Europe, and East Asia. Similarly, Villéger *et al*. (2015) found strong evidence that species introductions, especially of generalist and widespread taxa, are diminishing the distinctiveness of regional fish faunas, particularly in the Nearctic and Palearctic regions.

Bernardo-Madrid *et al*. (2019) provided a comprehensive assessment of birds, mammals, and amphibians, showing that introductions are fragmenting and redrawing global zoogeographical regions. Their findings highlight both homogenization—where regions merge—and fragmentation—where novel patterns emerge due to uneven invasion pressure and climate alignment. Parallel conclusions were reached by Liu *et al*. (2021) assessing the regionalisation patterns of non-native herpetofauna in their native and current ranges.

Additional studies confirmed that introductions are also reshaping global floristic patterns. Yang *et al*. (2021) and Daru *et al*. (2021) both demonstrated that non-native plants substantially reduce taxonomic and phylogenetic turnover among regions, leading to floristic convergence. These changes align with the “New Pangea” hypothesis, whereby ongoing globalization blurs the distinctions among historically unique floras. Further, Brown *et al*. (2023) showed that phytogeographical regions are being redefined or merged due to species introductions and extinctions, particularly where invasion pressure is uneven.

The breakdown of regional boundaries has also been documented for invertebrates. Recently, Aulus-Giacosa *et al*. (2024) found that introductions of non-native ants have merged formerly distinct tropical region into a single pantropical cluster, while Capinha *et al*. (2015) reported that the current ranges of gastropods with non-native ranges follows global climate more than geographical distance, forming novel regions aligned with temperature gradients rather than traditional dispersal barriers.

On islands, the erosion of biogeographical distinctiveness is also apparently pronounced. Capinha, Marcolin & Reino, (2020) showed that the composition of island herpetofaunas has become increasingly homogenized, with rising taxonomic convergence across Caribbean, Atlantic, and Indo-Pacific islands. Comparable trends were observed in island birds, with studies reporting widespread homogenization across archipelagos (Cassey *et al*., 2007; Soares *et al*., 2022).

These highly convergent results suggest that while natural geologic, geographic and climatic drivers historically shaped biogeographical regions, anthropogenic species introductions are now a dominant force, interacting with these natural drivers to reshape regions. The reshaped biogeographic regionalisation appears to be primarily driven by the combination of climate filtering and asymmetric species exchanges. Climate matching facilitates the establishment of non-native species in regions with similar environmental conditions, promoting convergence across distant but climatically similar areas (Capinha *et al*., 2015; Aulus-Giacosa *et al*., 2024). Asymmetric species exchange, where some regions act predominantly as donors (e.g., Asia, North America, Europe) and others as recipients (e.g., islands, Australasia), further drives global patterns of homogenization (Daru *et al*., 2021, Leroy *et al*., 2023). In parallel, the extinction of regionally distinct native species can further accelerate the breakdown of historical regions (Bernardo-Madrid *et al*., 2019; Leroy *et al*., 2023). Collectively, these processes are reshaping long-standing biogeographical boundaries, with novel communities increasingly structured by global climatic convergence and the accelerating influence of human-mediated species exchange.

### (3) Wallace’s Line

Wallace’s Line is arguably the most iconic biogeographical boundary globally, marking a sharp faunal break between Asian and Australasian regions (Wallace, 1860). This boundary reflects a deep evolutionary divergence and long-term isolation of Wallacea, a transitional zone within the Australasian bioregion located east of the Line. Our synthesis, though based on limited studies (n=5; Fig. 2; Fig. 3c), shows that human mediated introductions already lead to substantial faunal intermixing across Wallace’s Line.

In a comprehensive review of human-mediated animal introductions in Wallacea and the surrounding archipelagos, Heinsohn (2001) documented the confirmed or probable introduction of over 50 terrestrial vertebrate species into the region, including more than two dozen mammal species, highlighting substantial human-driven faunal mixing. These introductions, predominantly of Palearctic or Asian origin, have been particularly concentrated on islands near Wallace’s Line, such as Sulawesi and Lombok. Similarly, Yanuarita *et al*. (2020) reported the presence of approximately 20 non-native freshwater fish across the region. Further illustrating the erosion of biogeographical distinctiveness in the area, Herder *et al*. (2021) found that in ancient Lake Poso, Central Sulawesi, one of Asia’s deepest lakes and a centre of freshwater endemism, the number of non-native fish species already surpasses that of native species.

Wallace’s Line is also being blurred within the distribution of individual species and emblematic groups. For example, most placental mammal genera are confined to the west of the Line, with marsupials dominating to the east. However, macaques (*Macaca* spp.) represent a notable exception, occurring on both sides. Evans *et al*. (2020) used mitochondrial genome data to trace multiple eastward dispersals of *Macaca fascicularis*, some likely the result of ancient natural movements, while others, particularly into the Lesser Sundas, are attributed to human-mediated translocations within the past 10,000 years. A comparable pattern is observed for *Rasbora lateristriata*, a freshwater fish distributed across Java, Bali, Lombok, and Sumbawa. Molecular analyses by Kusuma *et al*. (2016) suggest that its recent expansion across Wallace’s Line was likely facilitated by human activity.

Together, these patterns underscore the increasing faunistic permeability of Wallace’s Line, driven by human-mediated species movements. While some cross-boundary introductions have historical origins, contemporary pathways are diverse, including trade, cargo transport, and pet releases (Heinsohn 2001; Yanuarita *et al*. 2020). These anthropogenic transportation vectors are accelerating species exchange across Wallace’s Line, fundamentally eroding this iconic boundary. With continuing globalization, this trend is likely to intensify.

### (4) Species-area relationships and island species-area relationships

Species-area relationships (SARs), and their island-specific form (iSARs) capture how species richness increases with area (Preston, 1960). These relationships have long been used to understand patterns of biodiversity and the effects of habitat size and isolation. Our synthesis identified a substantial body of research assessing how biological invasions relate to or affect (i)SARs worldwide (n=49; Fig 2). We show that non-native species frequently produce additive or interactive effects that reinforce or align with expected patterns.

Most identified studies examined (i)SARs for non-native assemblages (n=26; 53%), with the majority reporting positive relationships (n=23; Fig. 3d), consistent with theoretical expectations. Among those assessing the additive effects of non-native species on contemporary assemblages, the majority found a reinforcing role in shaping (i)SARs (9 out of 13; Fig. 3d). Three meta-analyses, Baiser & Li (2018), Guo (2014), and Guo *et al*. (2021b), have also compared native and non-native (i)SARs across various spatial and taxonomic contexts. These works show that (i)SARs for non-native species tend to be more variable than those for natives, particularly among vertebrates. In contrast, non-native vascular plants frequently display (i)SARs more similar to those of native species. When native and non-native assemblages are considered together, overall (i)SARs are reinforced and become steeper in highly invaded systems (i.e. exceeding natural expectations). This effect is typically more pronounced on islands, where high non-native colonization pressures and low native species diversity lead to a higher representation of non-native taxa. Examples of these influences are provided by van Der Geer, Lomolino & Lyras, (2017) and Matthews *et al*. (2023), who found that human-introduced mammals and birds in islands worldwide have compensated for extinction-driven losses and inflated species richness beyond historical baselines.

Some studies (n=10) also describe cases of non-native taxa affecting (i)SARs by altering native assemblages through ecological mechanisms such as predation, competition, or habitat modification. Their effects sometimes weaken (i)SARs of non-native assemblages (Fig. 3d). For example, invasive Norway rats in the Falkland Islands suppressed native passerine bird richness, flattening iSARs; following rat eradication, richness rebounded and iSARs began to resemble those of uninvaded islands (Tabak *et al*., 2015). Similarly, on Bahamian islands, introduced *Anolis* lizards reduced insect richness and iSAR slopes through predation, particularly affecting Diptera and Hymenoptera (Murakami and Hirao, 2010). However, some studies also reported neutral or even reinforcing effects. For example, in mainland Norway, introduced *Pinus mugo* increased vascular plant richness and steepened SARs by creating open, heterogeneous microhabitats, in contrast to native pine stands with dense canopies and low understory diversity (Vetaas *et al*., 2014).

Together, these studies demonstrate that non-native species consistently influence (i)SARs worldwide through both assemblage-level additive effects and ecological interactions. Reinforcing effects are most frequently reported (particularly in island and human-modified systems), while weakening effects, though less commonly reported, are typically associated with ecological disruptions caused by invasive species. These results contrast with patterns for previous rules where anthropogenic drivers led to either disruption or non-alignment with natural patterns. Here, the observations that non-native have patterns consistent with natural (i)SAR suggest that in this case, non-native species are generally driven by similar ecological processes as native species.

### (5) Island diversity-isolation relationship

A core prediction of island biogeography theory is that species richness tends to decline with increasing island isolation due to reduced immigration from source regions (MacArthur & Wilson, 1967). This inverse isolation-richness relationship has been well documented for native taxa across most island systems (Whittaker & Fernández-Palacios, 2007). Our review (n=25; Fig. 2) reveals that biological invasions markedly erode this pattern (Fig. 3e).

Among six broad-scale studies on non-native assemblages at global to continental scales, all reported non-aligned patterns from the expected negative isolation-richness gradient, with four indicating inverse (i.e., positive) relationships (Fig. 3e). This trend is broadly consistent across taxa. For example, Sánchez-Ortiz *et al*. (2020) analysing 7,718 sites across 81 islands worldwide observed higher non-native species richness on more isolated islands among plants, vertebrates, and invertebrates. Similarly, in a global analysis of over 250 tropical and subtropical islands, Moser *et al*. (2018) found that while native richness declined with isolation, non-native richness increased for mammals, ants, reptiles, and vascular plants, with no consistent pattern for birds. Crucially, when contemporary assemblages were investigated, the negative isolation-richness relationship became exceptionally weakened across all groups except birds.

Studies at regional scales report either a negative relationship or no significant associations between non-native species richness and island isolation (Fig. 3e). This variability was also highlighted by Guo (2014), whose meta-analysis of bird and plant invasions across island groups revealed inconsistent isolation relationships, shaped by taxonomic identity and geographic context. While positive relationships are seemingly uncommon at regional scales, all five studies assessing additive effects on contemporary assemblages have nonetheless documented a weakening of the isolation-richness gradient (Fig. 3e). This likely reflects weaker increases in non-native richness with isolation, reducing the overall effect. These outcomes are geographically and taxonomically widespread. For example, Helmus *et al*., (2014) found that native anole richness declined with isolation across 37 Caribbean islands, whereas non-native richness increased, substantially diminishing the isolation effect in combined assemblages. Similar patterns have been reported for the broader Caribbean herpetofauna (Gleditsch *et al*., 2023), vascular plants in the Lesser Antilles (Rojas-Sandoval, Ackerman & Tremblay, 2020), freshwater fishes in the Caribbean (Furness, Reznick & Avise, 2016), and Mediterranean island floras (Ficetola & Padoa-Schioppa, 2009).

The high accumulation of non-native species on isolated islands has often been linked to the intentional introduction of species to compensate for perceived deficits in native biodiversity, e.g., for agricultural, ornamental or recreational purposes (Sánchez-Ortiz *et al*., 2020). Remote islands also tend to support ecologically naïve native species that lack evolutionary defences against novel biota, thereby likely facilitating non-native invasions (Moser *et al*., 2018). Overall, these results highlight that contemporary drivers of the diversity of island biotas have shifted due to the breakage of biogeographic barriers, and, as a result, isolation no longer constrains species richness as strongly in today’s island systems. This disruptive trend is likely to continue in the future, because while some island systems currently have strong biosecurity measures, such as New Zealand and some oversea territories of Global North countries, most island systems lack legal instruments to prevent biological invasions (Lenzner *et al*., 2020).

### (6) Distance decay of similarity

Distance decay of similarity (DDS), the decline in community similarity with increasing geographic distance, is a well-known pattern in ecology and biogeography (Nekola & White, 1999). Among native species, DDS typically arises from a combination of dispersal limitations, environmental filtering, and shared evolutionary histories. However, our synthesis reveals that non-native taxa are disrupting this pattern, by mostly weakening, and in rare cases strengthening, the expected relationship, generally leading to increased biotic homogenization (Fig. 3f).

Most reviewed studies were performed at infra-continental scales (10 out of 12), with only one work assessing DDS at the global scale, and another at the continental scale (Fig. 3f). At the global scale, Capinha *et al*. (2015) reported a substantial weakening of distance decay of similarity (DDS) for non-native terrestrial gastropods in their current ranges versus their native ranges. However, at a continental scale, Leprieur *et al*. (2009) found that non-native freshwater fish in Europe have steeper DDS slopes than native species, a pattern attributed to the spatial clustering of introduction pathways at the continental scale. At regional scales, most studies identified weaker DDS patterns among non-native taxa relative to their native counterparts (Fig. 3f). For instance, Okimura, Koide & Mori (2016) found flatter DDS slopes for alien plant assemblages along roadside corridors in Japan, indicating reduced spatial turnover and diminished regional differentiation compared to native assemblages. Mologni (2022) reported that while native plant communities across New Zealand’s islands exhibited clear spatial turnover, non-native species showed no detectable DDS pattern. Likewise, La Sorte & Pyšek (2009) found that European archaeophytes introduced to North America exhibited virtually no DDS, in contrast to the pronounced distance decay observed among native plant assemblages. In addition, two studies found mixed results: Jehlík, Dostálek & Frantík (2019) observed stronger DDS among recently introduced neophytes in Central European river ports compared to archaeophytes and native species. La Sorte *et al*. (2008) also found group-specific differences: while natives and archaeophytes were associated with weaker DDS across urban floras, neophytes exhibited steeper slopes.

Two studies assessed the effects of non-native species DDS on contemporary assemblages. Reeve *et al*. (2022) found that current plant assemblages in Australian eucalypt understories exhibit flatter DDS slopes compared to native-only assemblages, a shift attributed to the broad distributions of common non-native species. Similarly, Ding *et al*. (2017) reported that fish communities in Chinese plateau lakes have undergone a substantial decline in DDS, reflecting a transition from historically high native turnover to increased homogenization, driven by widespread non-native introductions and endemic species loss.

Observed variation in DDS patterns for non-native assemblages largely reflect differences in introduction histories, the spatial distribution of propagule sources, and country-specific colonization pathways—all of which shape spatial gradients in species turnover (La Sorte & Pyšek 2009; Leprieur *et al*., 2009). These differing contexts are also resulting in variable and context-dependent impacts on contemporary assemblages. While impacts on contemporary assemblages vary across contexts, compositional similarity is increasingly decoupled from geographic distance and progressively more influenced by anthropogenic drivers.

### (7) Bergmann’s Rule

Bergmann’s Rule posits that animal body size increases with latitude or decreasing temperature, originally formulated for endotherms as a thermoregulatory adaptation (Bergmann, 1847). More recently, the rule has been extended or analogized to ectotherms and applied to a broader range of taxa and contexts (Vinarski, 2014). Our synthesis reveals that non-native species relate to this biogeographical rule in diverse ways, and importantly, have already reshaped its expression at broad scales.

Of the 21 studies identified (Fig. 2), only four have examined how non-native species relate to, or influence, Bergmann’s Rule at the assemblage level (Fig. 3g). Blanchet *et al*. (2010), analysing body size patterns of native and non-native freshwater fish across 1,058 river basins worldwide, found that species introductions significantly reshaped latitudinal trends. Prior to introductions, Bergmann’s Rule was evident only in the Northern Hemisphere. However, in contemporary assemblages, a latitudinal gradient consistent with Bergmann’s Rule also emerged in the Southern Hemisphere, driven largely by the introduction of larger-bodied species at higher latitudes. Relatedly, Blackburn, Redding & Dyer (2019) found that larger-bodied non-native birds were more frequently found at higher latitudes globally, generating a Bergmann-consistent pattern. Less consistent outcomes have also been found. Alò, Pizarro & Habit (2023), examining freshwater fish assemblages across a broad latitudinal and elevational gradient in Chile, found that native species more frequently conformed to Bergmann’s Rule than did non-native species, which displayed weaker or absent latitudinal size trends.

A much larger number of identified studies have assessed Bergmann’s Rule at the individual species level (Fig. 3g), with heterogeneous results. Several species clearly conform to the rule (n=8). For example, the European starling (*Sturnus vulgaris*) in Australia exhibited larger body sizes at cooler, higher-latitude sites (Cardilini *et al*., 2016). Similarly, the non-native fruit fly *Drosophila subobscura* developed a latitudinal wing size cline in North America, aligning with Bergmann’s Rule (Huey *et al*., 2000). Similar increases in body size with latitude have been observed in the invasive rice water weevil (*Lissorhoptrus oryzophilus*) in China (Huang *et al*., 2018) and in the Polynesian rat (*Rattus exulans*) across 105 island populations in Oceania and Wallacea (van der Geer, 2018).

Conversely, several species exhibit patterns that deviate from or contradict Bergmann’s Rule (n=9). For instance, the introduced common myna (*Acridotheres tristis*) in New Zealand showed larger body sizes in warmer northern regions, an inverse of the expected pattern (Baker & Moeed, 1979). The invasive Cuban treefrog (*Osteopilus septentrionalis*) also showed a decline in body size with increasing latitude across its introduced range (McGarrity & Johnson, 2008). Similarly, no consistent latitudinal body size clines were found in introduced populations of the emerald ash borer (*Agrilus planipennis*) or the Mediterranean house gecko (*Hemidactylus turcicus*) in North America (Marshall *et al*., 2013; Granatosky & Krysko, 2014). Interestingly, erosional effects over native species were also identified. On Marion Island, for example, house mice preyed selectively on larger native weevils, flattening or inverting body size-temperature relationships otherwise consistent with Bergmann’s Rule (Treasure & Chown, 2014).

A major driver of Bergmann-consistent patterns in non-native species is biased colonization pressure, with larger-bodied species disproportionately introduced at higher latitudes, an effect observed by Blanchet *et al.,* (2010) and Blackburn *et al.,* (2019). These patterns often reflect introduction history artifacts rather than post-establishment adaptation. Beyond this, mechanisms such as phenotypic plasticity, dispersal strategy and life-history traits have been shown to mediate or override climatic-related latitudinal effects (Jardeleza *et al*., 2022; McGarrity & Johnson, 2008; Alò *et al*., 2023). Together, these findings reveal that non-native species have already produced substantial shifts in spatial body size patterns, including the large-scale emergence of Bergmann-consistent gradients where none previously existed. While individual non-native species may adapt to climatic gradients, the emergence of Bergmann-like patterns at the assemblage level appears driven largely by anthropogenic introductions, highlighting how this mechanism is actively reshaping classical biogeographical rules.

### (8) Rapoport’s Rule

Rapoport’s Rule describes a biogeographical pattern in which species occurring at higher latitudes tend to occupy broader geographic, particularly latitudinal, ranges than those in tropical regions (Rapoport, 1982). This gradient is traditionally attributed to greater climatic variability and broader ecological tolerances in temperate and polar environments compared to tropical regions. A small number of identified studies (n=5; Fig. 2) have tested this rule in the context of non-native species, particularly at large spatial scales. While several studies report patterns consistent with Rapoport’s Rule, the underlying mechanisms often differ from those in native assemblages.

Three studies (Fig. 3h) found that non-native assemblages tend to exhibit broader latitudinal ranges at higher latitudes, aligned with Rapoport’s Rule. Dyer *et al*., (2020), in a global analysis of 355 alien bird species, found that latitudinal range extent increased with latitude. Similarly, Sax (2001) identified comparable trends across birds, mammals, plants, and freshwater fish on multiple continents. Kirk, Hays & Petranek (2021) further demonstrated that species originating from higher latitudes tend to establish broader latitudinal ranges when introduced, reinforcing the idea that native-range latitude can predict non-native spread.

However, not all studies support this pattern. Procheş *et al*. (2012), examining *Pinus* species globally, found that naturalized range sizes were primarily driven by introduction effort, not latitude of origin. Similarly, Mountier *et al*. (2018) observed that introduced ferns and lycophytes in New Zealand tended to have narrower latitudinal ranges than natives, likely due to shorter residence time and geographic constraints within the island system.

While abiotic stress filtering, such as cold tolerance, may explain some patterns consistent with Rapoport’s Rule, both conforming and deviating findings among non-native species appear to be primarily linked to spatial patterns of human-mediated introductions. Broad latitudinal ranges frequently reflect widespread or high-frequency introductions rather than adaptive tolerance to climatic variability (Dyer *et al*., 2020; Procheş *et al*., 2012; Mountier *et al*., 2018). Overall, while non-native assemblages frequently display range size gradients consistent with Rapoport’s Rule, these patterns are strongly influenced by human introduction dynamics, underscoring their increasingly important role over natural drivers as time progresses.

### (9) Gloger’s Rule

Gloger’s Rule proposes that animals in warmer, more humid climates tend to exhibit darker pigmentation, while those in cooler, drier regions are typically lighter (Delhey, 2019). Identified studies assessing this rule in non-native species are few (n=5; Fig. 2), but also highlight a range of outcomes, from clear conformity (n=3) to pronounced divergence (n=1; Fig. 4a).

**Fig. 4.**
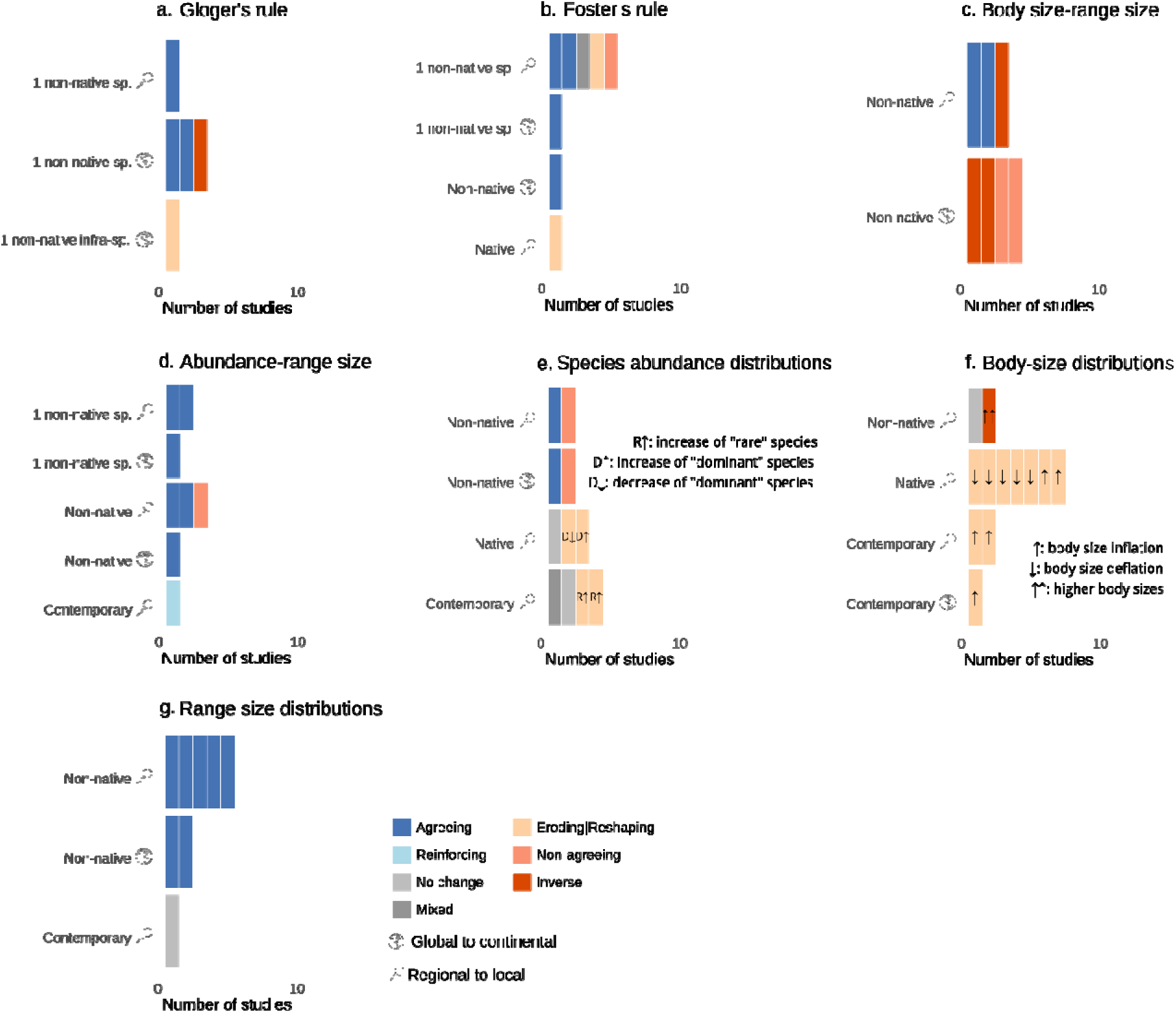
Synthesis of studies assessing the patterns or effects of non-native species in relation to (a) Gloger’s rule, (b) Foster’s rule, (c) body size-range size relationship, (d) abundance-range size relationship, (e) species abundance distributions, (f) body size distributions and (g) range size distributions. Plots show the number of studies (horizontal axis) per rule. Colours indicate the nature of agreement with theoretical expectations. Blue tones in the right indicate results that align with the expected biogeographical pattern or rule, orange and red tones indicate deviations. Grey tones denote mixed results or no observed effect.

A remarkable example of conformity comes from house sparrows (*Passer domesticus*) introduced to North America in the 1850s, where pigmentation patterns have diverged markedly from the ancestral stock within just over a century. These introduced populations now show significantly darker plumage in wetter northern and coastal regions and paler coloration in arid southwestern areas (Johnston & Selander, 1964). In regions of North America, feral pigs (*Sus scrofa*) also exhibited darker eumelanin-based pigmentation in areas characterized by higher temperatures, humidity, and UV radiation, patterns consistent with Gloger’s Rule (Newell, Walker & Caro, 2021). Similarly, Lahti (2008) found that eggshell coloration in village weavers (*Ploceus cucullatus*) shifted toward a uniform medium blue-green in response to solar radiation after their introduction to the Caribbean and Indian Ocean islands, suggesting a climate-driven pigmentation response.

However, deviations from Gloger’s Rule have also been documented. In the ladybird *Coccinella septempunctata*, native Eurasian populations displayed darker pigmentation in more humid environments, as predicted by the rule, but in the introduced North American range, darker individuals were found in colder areas (O’Neill *et al*., 2017). In Israel, native house sparrows (*Passer domesticus*) exhibited an overall clinal pattern in line with Gloger’s Rule; however, unusually lighter plumage in the southern population was attributed to the likely introduction of a non-native subspecies, *P. d. indicus* (Ben Cohen & Dor, 2018).

Taken together, these findings suggest that while Gloger’s Rule can manifest in non-native species through adaptation to local climatic conditions, such patterns may also be shaped, or disrupted, by evolutionary shifts or pre-existing trait variation. However, the current evidence base is limited, preventing broad conclusions about the generality of Gloger’s Rule in the context of biological invasions and their effects on broader biogeographic patterns.

### (10) Foster’s Rule (Island Rule)

Foster’s Rule, or Island Rule, describes a biogeographical pattern whereby small-bodied species tend to evolve larger body sizes (insular gigantism), and large-bodied species tend to become smaller (insular dwarfism) when isolated on islands (Foster, 1964). Identified studies (n=8; Fig. 2) indicate that non-native species frequently undergo body size shifts consistent with this rule and affect these trends in native taxa, despite some indications of context-dependency (Fig. 4b).

At the non-native assemblage level, van der Geer, Lomolino & Lyras (2018) analysed a comprehensive dataset of 385 populations of 56 non-volant mammal species introduced to 285 islands worldwide. They found systematic body size shifts consistent with Foster’s Rule: smaller species typically became larger and larger species smaller, with more pronounced changes in populations with longer isolation times. Additional species-level studies support these patterns. For instance, the Polynesian rat (*Rattus exulans*) exhibited consistent insular gigantism across more than 100 island populations (van der Geer, 2018), particularly on smaller, species-poor islands. In Bermuda, the great kiskadee (*Pitangus sulphuratus*) evolved a ∼5% increase in body mass and bill width within 50 years of introduction (Mathys & Lockwood, 2009). Similarly, invasive guttural toads (*Sclerophrys gutturalis*) on Mauritius and Réunion displayed insular dwarfism not observed in their native African populations (Baxter-Gilbert *et al*., 2022).

However, these patterns are not universal. Several studies highlight that ecological interactions can modulate, or even reverse, body size shifts. Barun *et al*. (2015) found that introduced mongooses (*Herpestes auropunctatus*) followed Foster’s Rule only on predator-free islands but exhibited reduced body sizes in the presence of native martens, suggesting character displacement. Likewise, black rats (*Rattus rattus*) on islands with high trophic complexity showed suppressed gigantism compared to populations on more pristine islands (Russell *et al*., 2011), and house mice on Antipodes Island showed no significant size change over four decades (Russell, 2012).

The long-term potential of non-native species to disrupt insular body size trajectories has also been demonstrated. Using palaeontological data from four Mediterranean islands, van der Geer *et al*., (2013) found that native small mammals often evolved larger sizes in the absence of predators and competitors. These trends, however, were frequently halted or reversed following the arrival of invasive species such as the rodents *Mus musculus*, *Rattus* spp., and *Apodemus sylvaticus*.

Altogether, these findings demonstrate that non-native species can undergo rapid, directional body size evolution consistent with Foster’s Rule, while also reshaping its expression in native species. Key drivers include isolation time, ecological release, inter- and intraspecific interactions, and island characteristics such as area, latitude, and elevation. While additional broad-scale assessments would be valuable, the consistency of findings across available studies, coupled with the high prevalence of invasions on islands (Dawson *et al*., 2017), suggests that non-native species may already be significantly reshaping body size patterns in islands globally.

### (11) Body size-range size relationships

Another long-established macroecological pattern is the positive relationship between a species’ body size and its geographic range size, with larger-bodied species generally occupying broader ranges (Gaston & Blackburn, 1996). However, reviewed studies (n=7; Fig. 2) strongly suggest that this rule is often disrupted in invasion contexts (Fig. 4c).

Most available evidence (n=5), typically involving broader taxonomic and geographic scopes, reveals deviations from this pattern. At a global scale, Dyer *et al*. (2016) found no consistent relationship between body mass and alien range size in 319 bird species. Instead, factors such as colonization pressure and native range size emerged as stronger predictors. Duncan *et al*. (2001) and Forsyth *et al*. (2004) found that smaller-bodied birds and mammals achieved broader distributions in Australia, contradicting the expected positive scaling. Similar trends were observed by Allen *et al*. (2013) in Florida, where smaller-bodied birds and reptiles exhibited greater range expansion. In marine invertebrates, Byers *et al*. (2015) also reported weak or negative associations between body size and introduced range size.

Still, two smaller-scale studies do report body size-range size relationships in line with classical expectations. Specifically, Su *et al*. (2016) found that larger-bodied alien birds had broader introduced ranges in Taiwan and Roy, Jablonski & Valentine (2002) documented a similar pattern among a set of 13 non-native bivalves on the Pacific coast of North America. These examples highlight some context-dependency and suggest that in some cases, larger size may indeed be associated with wider ranges in non-native taxa.

Taken together, these findings indicate that the expected positive relationship between body size and range size is not consistently upheld in invasion contexts. Deviations appear driven by introduction dynamics, including species use (e.g., game species often being large-bodied), transportability, and propagule pressure (e.g., small-bodied species may be more easily transported and go unnoticed, resulting in higher introduction rates). Other factors, such as ecological generalism and time since establishment, also play a role, as many introduced species are still expanding and have not reached range equilibrium (e.g., Dyer *et al*., 2016). Despite this complexity, the recurring nature of these deviations across taxa and regions (e.g., Duncan *et al*., 2001; Dyer *et al*., 2016) suggests that biological invasions may already be actively reshaping size–distribution relationships in contemporary biotas.

### (12) Abundance-range size relationships

A well-established pattern in macroecology is the positive triangular relationship between a species’ local abundance and its geographic range size, the abundance-range size (ARS) relationship (Holt *et al*., 1997). Species that occupy small geographic areas cannot have a locally high abundance, whereas species locally abundant tend to occupy broad geographic areas. Our synthesis shows that non-native species frequently conform to this rule, sometimes even reinforcing ARS patterns in contemporary assemblages.

Among the eight studies identified, seven reported significant positive ARS correlations (Fig. 4d). Fristoe *et al*. (2021), drawing on over one million vegetation plots across Europe, found strong positive correlations between local abundance and geographic range size for long-established non-native plant species. This pattern closely mirrors that observed for native European flora, where species that are more locally abundant also tend to be more geographically widespread. Similarly, in a study of 70 terrestrial invasive plant species across the continental United States, O’Neill, Bradley & Allen, (2021) identified a significant positive correlation between establishment range size and abundance range size. Osunkoya *et al*., (2020) also reported consistent positive abundance-distribution relationships for 64 invasive plant species across Queensland, Australia. Notably, Russell *et al*., (2005) found that while native vascular plants in New Zealand grasslands exhibited no significant ARS relationship, non-native species showed strong and significant positive correlations. When both groups were analysed together, the effect of non-natives created a significant ARS pattern.

Positive ARS relationships have also been identified at the individual species level. Ewers *et al*. (2023), showed that the global invader Chinese mitten crab (*Eriocheir sinensis*) exhibited temporally linked increases in abundance and geographic range, with regional abundance increases typically preceding range expansion. Similar species-level ARS dynamics were observed for pink salmon (*Oncorhynchus gorbuscha*) in the Norwegian and Barents Seas (Pauli *et al*., 2023), and for upslope-expanding ship rat (*Rattus rattus*) in New Zealand forests (Harris *et al*., 2022).

Despite general support for the ARS relationship, at least one exception has also been documented. Guan *et al*. (2023), examining 32 invasive plant species on the high-altitude Tibetan Plateau, found no significant correlation between abundance and range size, suggesting that ARS patterns for non-native species hold some level of context-dependency.

Several mechanisms may explain this variability. Fristoe *et al*. (2021) identified time since introduction as a key factor, with older introductions showing stronger ARS correlations—likely due to longer periods for demographic growth and spatial spread. Functional traits such as habitat generalism, dispersal ability, and growth strategy also likely contribute, though their individual effects can be hard to isolate. Given that natural ARS relationships already arise from complex, interacting processes (Borregaard & Rahbek, 2010), the addition of non-native species and their indirect drivers adds a further layer of complexity. Overall, ARS appears broadly applicable to non-native taxa, often reinforcing this macroecological pattern. However, its strength and expression remain shaped by anthropogenic factors, including introduction histories and the selection of species traits, highlighting the role of human context in contemporary abundance–range size dynamics.

### (13) Species abundance distributions

Species abundance distributions (SADs), i.e. the patterns that describe how common or rare species are in terms of abundance within an area, are a core component of community structure and biodiversity theory. SADs typically exhibit a right-skewed shape, where most species are rare and only a few are dominant (Gaston & Blackburn, 2000). Identified studies (n=11; Fig. 2) indicate that non-native species affect SADs in variable ways, often causing notable disruptions (Fig. 4e).

Studies examining SADs in non-native assemblages revealed both similarities to (n=2) and departures (n=2) from native community patterns. Hansen *et al*. (2013), studying aquatic ecosystems across Hawaii, North America, and Europe, found that invasive and native species shared similarly right-skewed SADs, indicating comparable ecological structuring. Similarly, Bennett *et al*. (2012), observed that non-native plant species in British Columbia meadows displayed SADs largely consistent with those of native species. In contrast, Labra, Abades & Marquet (2005), reported that non-native birds in North America exhibited more even abundance distributions than native birds. Hulme (2008), found a different pattern, with non-native plants in the UK disproportionately represented among rare species and underrepresented among dominants.

Studies assessing the effects of non-native species on native or contemporary assemblages (n=7) reported varied outcomes, with SAD pattern disruptions being commonly documented (n=5; Fig. 4e). For example, studies of Azorean arthropod communities have shown that non-native species tend to occupy the rare end of the SAD, increasing bimodality and multimodality in community structure (Matthews, Borges & Whittaker, 2014; Boieiro *et al*., 2018; Tsafack *et al*., 2022). Similarly, changes driven by dominant invaders reshaping community structure were also documented. Kortz & Magurran (2021), found that invasion by the pine *Pinus elliottii* in the Brazilian Cerrado led to reduced evenness and increased dominance. Pires-Teixeira *et al*. (2021) reported comparable shifts in marine rocky reef communities invaded by corals and algae, where dominance by invaders fundamentally altered abundance distributions.

Collectively, these findings show that while SADs in non-native assemblages can resemble those of native communities, reshaping effects are common. Such changes may result from the additive effects of non-native species but are also often driven by ecological interactions, such as competition, predation, or habitat alteration, that shift native species’ abundances. Invasions can reduce evenness through dominance by a few taxa or increase rarity by introducing many low-abundance species. Although outcomes vary with context, species traits, and invasion history, there is clear evidence that invasions can significantly restructure community abundance patterns.

### (14) Body size distributions

Body size distributions are fundamental descriptors of ecological communities and typically exhibit right-skewed (L-shaped) patterns, characterized by a predominance of small-bodied species and few large-bodied ones (Gaston & Blackburn, 2000). Our synthesis (n=12; Fig. 2) indicates that biological invasions frequently and substantially impact these patterns, either through direct impacts of introduced species on native assemblages or via the introduction of species with distinctive size traits (Fig. 4f).

A notable demonstration of the effects of biological invasion on body size distributions is given by Blanchet *et al*. (2010), who examined freshwater fish introductions worldwide and found that non-native species were disproportionately large-bodied compared to natives. These introductions resulted in significant changes of body size distributions, including increases in median body size and altered distributional shape especially in highly invaded regions of the Southern Hemisphere. Similarly, Roy, Jablonski & Valentine (2001), found that introduced marine bivalves along the California coast were significantly larger-bodied than native species. A comparable pattern was reported for non-native marine bivalves in the Mediterranean Sea, where Lessepsian invaders often include larger-bodied species that shift native assemblage structure (Nawrot *et al*., 2017). Likewise, Vitule *et al*. (2012), documented that non-native freshwater fish in the Upper Paraná, Brazil were typically larger than native species, contributing to community-wide body size inflation. However, not all systems show consistent reshaping. On Marion Island, a comprehensive study by Gaston, Chown & Mercer (2001), found that although introduced arthropods were slightly larger-bodied on average, their presence did not significantly alter the overall shape of the body size distribution.

Non-native species also frequently alter the body size structure of native assemblages. In New Zealand, rodents introduced by humans disproportionately impacted large-bodied invertebrates, flattening historically right-skewed distributions (Gibbs, 2009). Similar disruptions have been linked to habitat alteration. In Spanish shallow lakes, common carp (*Cyprinus carpio*) reduced large-bodied zooplankton via sediment disturbance and direct predation, favouring smaller, tolerant taxa (Florian *et al*., 2016).

Likewise, habitat alterations driven by non-native muskrats (*Ondatra zibethicus*) in Finnish wetlands led to increased dominance of smaller-bodied species (Nummi, Väänänen & Malinen, 2006). Selective predation by introduced opossum shrimp (*Mysis diluviana*) in Flathead Lake, USA (Ellis *et al*., 2011), and by *Gambusia holbrooki* fish in New South Wales, Australia (Hinchliffe *et al*., 2017), have also altered invertebrate communities by removing large-bodied taxa. In contrast, in Brazilian reservoirs, invasive peacock bass (*Cichla* spp.) selectively suppressed small-bodied native fish, shifting community composition toward mid-sized forms (Franco *et al*., 2022). Similarly, in Harp Lake, Canada, the introduction of a predatory water flea (*Bythotrephes cederstrœmi*) led to declines or local extinctions of small-bodied crustacean zooplankton species, while larger species increased in abundance, fundamentally reshaping the size structure of the native community (Yan & Pawson, 1997).

Overall, available evidence demonstrates that biological invasions are substantially reshaping body size distributions across diverse taxa and ecosystems, both through the introduction of species with non-random size traits (particularly large-bodied species, e.g., Blanchet *et al*. 2010) and through disruptive ecological impacts such as predation and habitat alteration. These effects challenge the stability of classical right-skewed patterns and highlight the growing role of human-introduced taxa in restructuring body size patterns of contemporary biota.

### (15) Range size distributions

Species-range size distributions, describing how many species occupy large versus small geographic areas, are one of the most universal patterns in biogeography and typically exhibit a right-skewed (L-shaped) pattern, with most species having small ranges and few achieving broad distributions (Gaston, 1996). Identified studies (n=8; Fig. 2) indicates that biological invasions frequently replicate this pattern but can also introduce subtle shifts linked to the dynamics of species introductions and establishment (Fig. 4g).

Across both terrestrial and aquatic ecosystems, non-native species consistently exhibit strongly right-skewed range size distributions. In New Zealand, Mountier *et al*. (2018), found that non-native ferns and lycophytes were primarily restricted to small geographic ranges, with only a limited subset expanding more broadly over time, patterns associated with residence time and habitat generalism. Comparable distributions have been reported for non-native plants in Europe (Weber, 1997; Williamson *et al*., 2009) and across the United States (McKinney, 2004). In aquatic ecosystems, Leroy *et al*. (2023) found that the global pool of non-native fishes were disproportionately sampled from the pool of naturally widespread species, resulting in a L-shaped pattern even more right-skewed than under a null assumption where introduced species would be randomly selected. At a regional scale, Vander Zanden, Hansen & Latzka (2017) also found that most non-native fishes and invertebrates in the U.S. occurred in only a few locations, while a small number became widely distributed invaders.

Despite broadly conforming to the expected L-shaped distribution, several studies document deviations when compared with native assemblages. Seabloom *et al*. (2006), for instance, showed that exotic plants in California had modal range sizes an order of magnitude smaller than those of comparable native species, likely reflecting incomplete range filling. Similarly, La Sorte & McKinney (2006) reported that non-native plants in urban green spaces of the eastern United States exhibited a more pronounced right skew than native species, characterized by many narrowly distributed species and a few with extremely broad ranges. Williamson *et al*. (2009) also emphasized the importance of residence time, noting that many naturalized neophytes initially occupy smaller ranges than native taxa, but tend to expand with time.

Taken together, these findings suggest that while non-native assemblages can introduce deviating signals reflecting introduction histories, dispersal limitations, and time since establishment, they often conform to or reinforce the classical right-skewed distribution of range sizes. Notably, this reinforcement of the skewness due to non-natives may be linked to the fact that naturally widespread species are much more likely to be introduced, as identified in multiple global-scale studies (Leroy *et al*. 2023, Pili *et al*. 2024). As non-native species continue to accumulate and expand their distributions, further restructuring may occur, however, to date their contributions appear largely consistent with theoretical expectations.

## III. CROSS-PATTERN SYNTHESIS AND FUTURE RESEARCH NEEDS

Our synthesis reveals that the distributional patterns of non-native species vary considerably in their alignment with classical biogeographical and macroecological rules, while also exerting a wide range of effects on broader biodiversity patterns (Fig. 5). Notably, multiple foundational patterns in contemporary biotas are already being strongly and consistently reshaped by biological invasions, particularly at broad spatial scales. These include the redrawing of global biogeographic regions across multiple taxa, the breakdown of the island isolation-diversity relationship, the emergence of Bergmann’s Rule-consistent body size gradients in areas and for taxa where there was none, and widespread shifts in community body size distributions.

**Figure 5.**
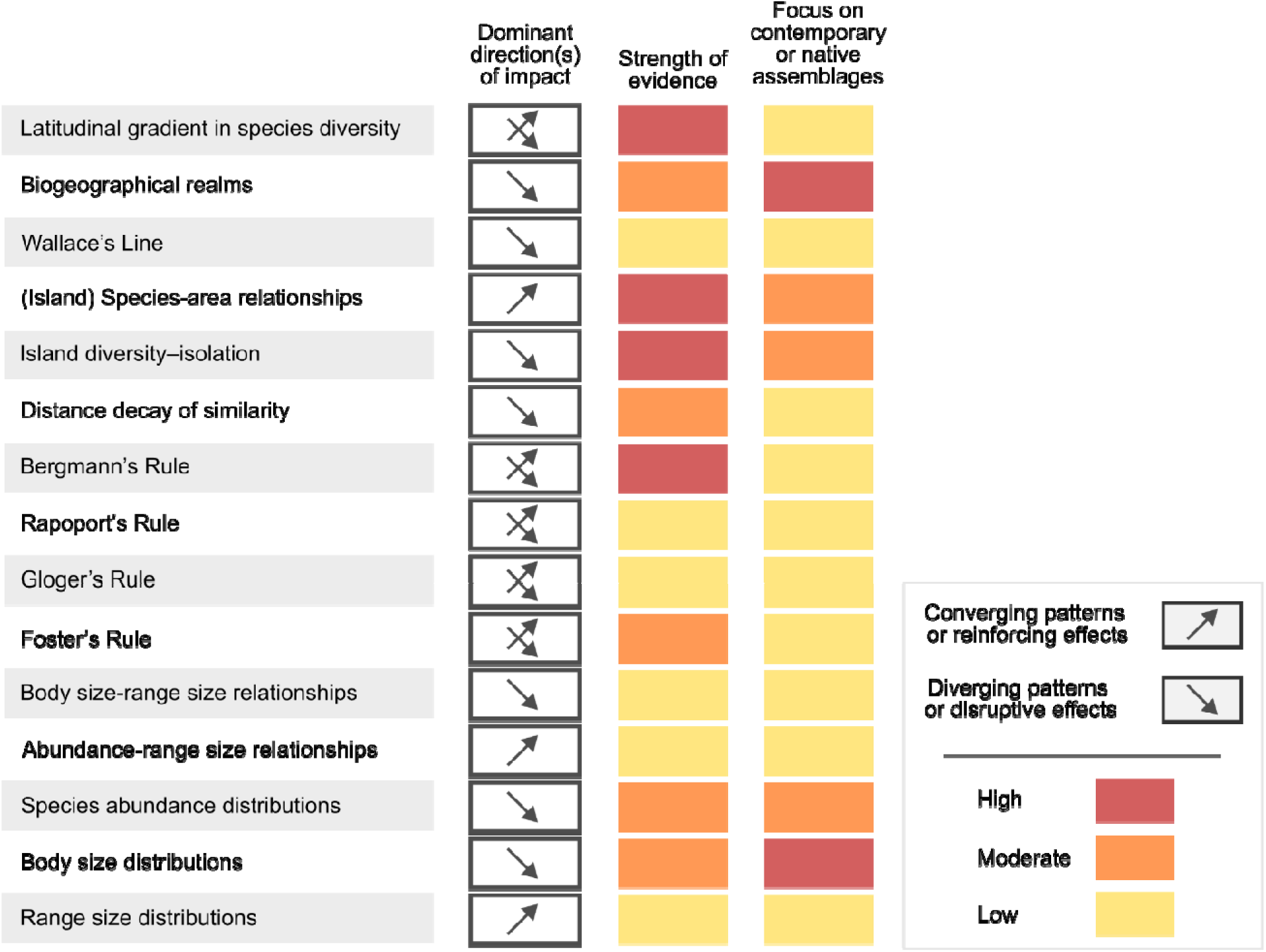
Qualitative synthesis of identified evidence on the impacts of non-native species across the assessed biogeographical and macroecological patterns. Rows represent individual patterns or rules. Columns show (1) the joint dominant alignment of patterns observed for non-native species and the nature of their effects on native or contemporary assemblages (2) the strength of evidence, based on number and comprehensiveness of studies available, the consistency of findings and the diversity of systems analysed; and (3) the share of available evidence specifically assessing effects on native or contemporary assemblages.

In contrast, certain macroecological patterns, such as the abundance-range size relationship and range size distributions, appear to remain largely consistent, or even more pronounced, across both native and non-native assemblages (Fig. 5). Despite some variability, species-area relationships, especially in their island-specific form, also tend to be reinforced under invasion, particularly in island systems where native species richness is low and colonization pressure is high. Such reinforcing effects may contribute to the persistence of structural regularities in Anthropocene biotas, even as other dimensions of biodiversity undergo significant disruption.

Altogether, these results ascertain that human activities have now become a major driving force affecting at the largest scale the natural patterns of biodiversity. However, understanding the factors that drive variation in the effects of biological invasions across different biogeographical patterns and rules remains an unresolved challenge. Explaining why certain patterns (e.g., the abundance–range size relationship) appear robust under invasion, while others (e.g., isolation–diversity relationships) break down, will likely require integrating human drivers with species’ functional traits, evolutionary histories, and environmental contexts. This is a complex challenge, as biogeographical and macroecological patterns and rules often result from multiple interacting drivers and processes (Gaston & Blackburn 2000, Lomolino *et al*., 2017), on top of which human drivers add further complexity. In addition, we can expect interactions among patterns and rules, whereby the impact of biological invasions on certain patterns will cascade onto others. For example, part of the hypotheses explaining the natural distribution of species range sizes were related to the natural distribution of body sizes and the relationship between body sizes and range size (Lomolino *et al*., 2017), which would imply cascading effects among the impacts of these rules. Likewise, the mechanisms underlying the reshaping of biogeographical regions were linked to the change in range size distributions, whereby the fact that widespread species (which, due to their overlapping distributions that cluster in specific areas, form the basis of biogeographical regions) are disproportionately introduced in turn explain the severity of the reshaping (Leroy *et al*., 2023).

For many of the patterns assessed, however, the available evidence remains limited (e.g., Wallace’s Line) (Figs. 2 5), often based on a small number of empirical studies, with narrow taxonomic or geographic focus. In the clearest cases (e.g., Gloger’s Rule), results tend to be context-dependent or inconsistent, limiting our ability to draw general conclusions about invasion-driven impacts. This uncertainty is further compounded by a disproportionate focus on non-native assemblages in isolation, rather than on their interactive or additive effects on native or contemporary communities. For example, this bias is particularly evident in studies of the latitudinal gradient in species diversity, where available evidence indicates consistent divergence in non-native species distributions at broad scales. However, their cumulative impact on overall biodiversity patterns remains poorly understood. While such studies are valuable for identifying tendencies to conform to or diverge from classical expectations, they currently fall short of capturing the broader biogeographical consequences of biological invasions for contemporary biodiversity.

In addition, important taxonomic, geographic, and ecosystem-level gaps persist. While some patterns or rules are inherently relevant only to specific taxonomic groups (e.g., Wallace’s Line; Bergmann’s Rule, Gloger’s Rule), the overall body of evidence remains heavily skewed toward plants and vertebrates (Fig. 6a). Similar biases are evident in ecosystem representation, with most studies conducted in terrestrial and, to a lesser extent, freshwater environments (Fig. 6b). Expanding research efforts toward marine environments, underrepresented taxa (e.g., invertebrate taxa, fungi, protists) and geographic regions (e.g., the tropics and Southern Hemisphere) remains critical to fully capture the extent and diversity of invasion-driven biogeographical change.

**Figure 6.**
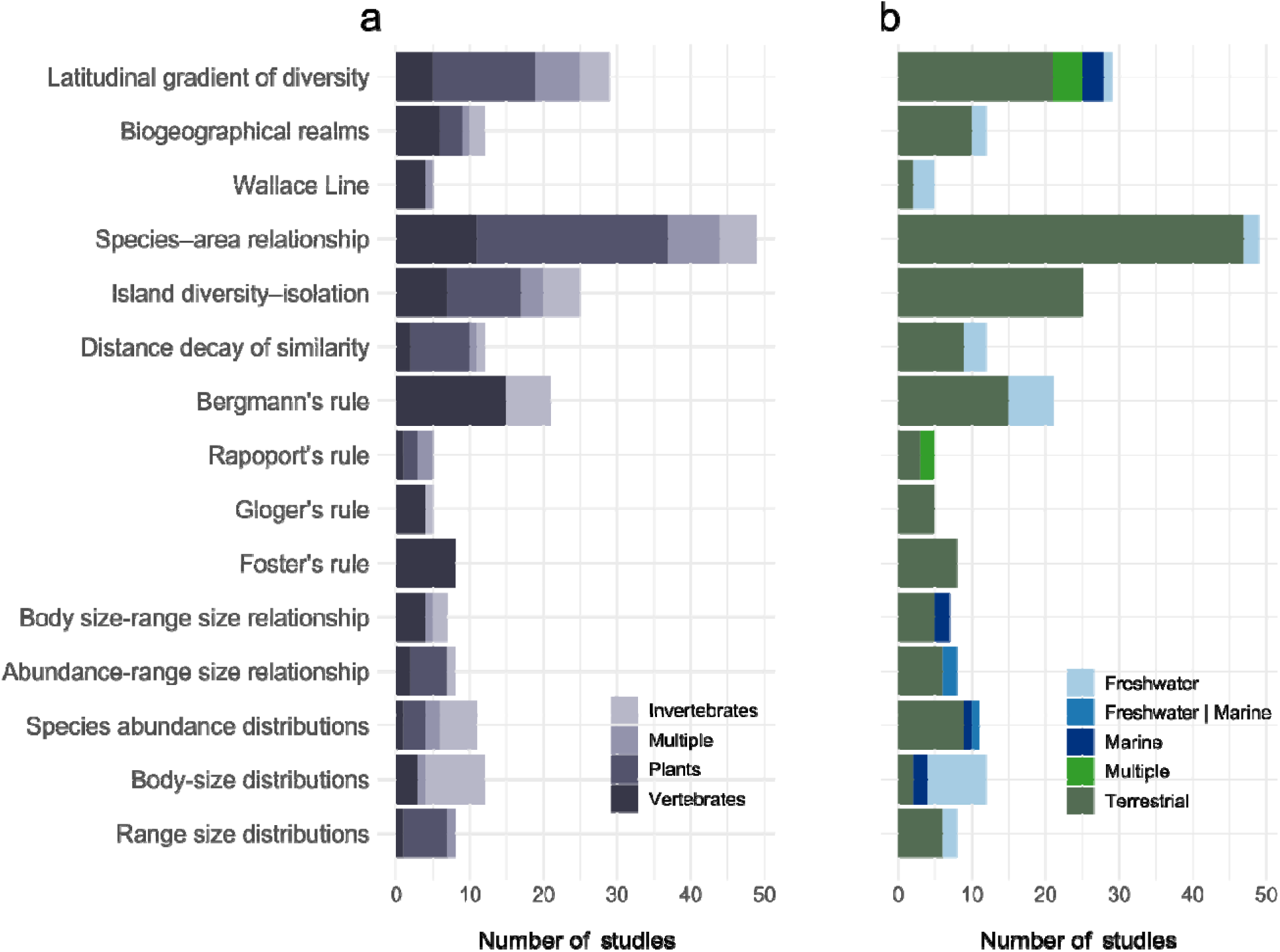
Distribution of reviewed studies across (a) major taxonomic groups and (b) environmental domains for each assessed biogeographical and macroecological pattern.

It would also be important to have longitudinal and high-resolution studies to disentangle the temporal dynamics of invasions and their cumulative effects on spatial patterns. Patterns of non-native species have been shown to be sensitive to the spatial resolution of sampling (e.g., Fridley *et al*., 2007), underscoring the importance of understanding how these patterns scale across different spatial grains and extents. In addition, many of currently observed patterns and effects are likely to be transitional since many introduced species have not yet reached equilibrium in their distributions, reflecting both lagged responses and ongoing processes. Assessed studies frequently identified human introduction patterns (including colonization pressure, introduction history, and asymmetrical species flows) as major drivers of spatial outcomes. For example, consistent deviations from the latitudinal diversity gradient in non-native assemblages result from disproportionately high introduction rates into temperate regions (Dawson *et al*., 2017), while patterns relating range size and trait-environment associations are frequently structured by colonization pressure and region of origin (Leroy *et al*. 2023, Briski *et al*., 2024; Pili *et al*. 2024). In this regard, as non-native species spread beyond places of introduction, the spatial imprint of human introductions may fade, potentially allowing other factors such as climatic and ecological constraints to reassert greater influence. This temporal effect has been demonstrated, for example, in assessments of distance decay of similarity (DDS), where recently introduced neophytes exhibit stronger DDS patterns, attributed to the spatial clustering of early introduction events, compared to archaeophytes, which have had more time to disperse (La Sorte *et al*., 2008; Jehlík *et al*., 2019). However, as time progresses, new introductions are likely to continue, and may even intensify, sustaining anthropogenic signatures in species distributions (Seebens *et al*., 2021). Establishing time series of non-native species impacts across spatial scales will be critical to detect delayed disruptions and understand how invasion-driven patterns evolve over time.

Finally, the major changes to biogeographical and macroecological rules and patterns highlighted in our review highlight two important consequences for fundamental and applied science in ecology and biodiversity conservation. First, it is now well established that even scientific assumptions should adapt to global change, questioning whether the reference unit should be natural or contemporary assemblages. This question should expand to whether the investigation of natural assemblages carries or not a hidden footprint of human activities, especially for lesser-known taxa for which introductions patterns are undocumented. Second, there is a pressing need to bridge biogeographical change with applied conservation. As invasions alter long-standing spatial templates of biodiversity, forecasting future scenarios under combined pressures of invasions, climate change, and land-use transformation, will require incorporating shifting baselines and hybrid assemblages in analytical and modelling frameworks. Hence, consideration of invasion impacts into biogeographical theory is not only a conceptual challenge but also a practical necessity for anticipating and managing biodiversity in the Anthropocene.

## V. CONCLUSIONS

1. Biological invasions are now a fundamental and accelerating force of biogeographical transformation. By facilitating the movement of species beyond historical biogeographical barriers, human activity is actively redrawing long-established spatial patterns, from the breakdown of classical diversity gradients and trait-based rules to the erosion of biogeographical regions and boundaries.
2. Species introductions lead to heterogeneous changes in biogeographical and macroecological patterns and rules. While some patterns are substantially disrupted, others remain surprisingly robust. These responses also vary across spatial and temporal scales, taxonomic groups, and introduction histories, reflecting a complex and contingent reorganisation of biodiversity.
3. We are still far from understanding the processes by which species introductions interact with biogeographical rules, due to the complex interactions between anthropogenic and natural drivers. Current insights are also constrained by taxonomic, geographic, and temporal biases, and often lack integration across spatial and temporal scales. Future research should prioritise the investigation of contemporary assemblages, cross-scale analyses, incorporate trait-based and functional dimensions, and adopt analytical frameworks that capture the multidimensional and dynamic nature of invasion-driven change over time.
4. The co-assembly of native and non-native species is giving rise to novel spatial configurations of biodiversity. Understanding and managing this reorganisation will require bridging macroecology, historical biogeography, and applied conservation to develop predictive models and responsive strategies for a rapidly changing biosphere.

## VI. METHODS

To assess how non-native species impact biogeographical and macroecological rules, we performed a systematic literature review following a predefined and reproducible search protocol. Our objective was to identify empirical studies evaluating these impacts using a consistent analytical framework across rules.

Between June and July 2024, we performed structured literature searches in Web of Science (WoS). For each rule, we used tailored search terms that combined the rule’s name with associated keywords (e.g., rule name + relevant terms), paired with widely used invasion-related terminology (e.g., “invasive,” “alien,” “non-native,” “exotic”). The complete list of keywords used, WoS queries, query links (with listed results), and the number of documents retrieved per rule are provided in Table S1 (Appendix 1).

To capture potentially relevant literature not indexed in WoS, including grey literature, we supplemented our search with additional queries in Google Scholar, performed in April 2025. Simplified keyword combinations emphasizing core terms were used to maximize the chances of retrieving relevant results across document types. Considering Google Scholar scoring of results by relevance (Nourbakhsh *et al*., 2012), we assessed the first 20 results for each query. Query terms, and search results are given in Supplementary Text 1 (Appendix 1).

During the screening and data collection process (see below), we also added studies cited within retrieved documents when deemed relevant to our aims. In total, the combined search efforts yielded 2,819 records: 2,484 from Web of Science, 300 from Google Scholar, and 35 manually identified.

An initial screening of each study was carried out through manual review of title, abstract, and full text (when necessary) to determine the alignment with our scope and potential for data supply. To meet our eligibility criteria, studies had to document patterns or effects of non-native species relevant to the rule of interest, under real-world conditions, and be based on empirical data. We included review or perspective type articles if they presented data analyses. Studies based solely on laboratory or controlled experiments were excluded, as our focus was on results observed under natural conditions. Studies passing this screening were then assessed for relevant data extraction.

For data extraction, we categorized results according to three main ecological units of analysis: 1) non-native assemblages or individual non-native taxa (species and subspecies); 2) native assemblages or individual native (sub)species, and 3) contemporary assemblages (i.e., current communities including both native and non-native species; Table 1). For non-native taxa, we recorded whether reported patterns: (i) aligned with the theoretical expectation of the rule, (ii) showed an inverse relationship, or (iii) exhibited other non-agreeing patterns. For native and contemporary assemblages, we classified non-native species’ effects into three categories: reshaping or eroding effects, indicating a weakening or deviation from expected patterns; reinforcing effects, indicating a strengthening of expected patterns; and no change, when no clear effect was observed. A fourth category, ‘mixed’ effects, was used when studies reported variable outcomes across distinct contexts (e.g., differing effects in different regions).

For each result, we also recorded: (i) the ecological unit analysed (e.g., assemblage or species level); (ii) the inferred mechanism, either additive effects (species additions) or ecological interactions by dominant invaders (e.g., predation, competition, habitat modification); (iii) the spatial scale (local/regional or continental/global); (iv) the taxonomic group (plants, vertebrates, invertebrates, fungi, or other); and (v) the environment type (terrestrial, freshwater, or marine).

After applying all eligibility and scope criteria, a total of 211 unique studies were retained. The total number of rule-specific observations analysed was slightly higher (n=217; Fig. 2), as some studies contributed to more than one rule. PRISMA flow diagrams (Page *et al*., 2021) for each rule are given in Figs S1 to S15 (Appendix S1).

Figures 1 to 4 and 6 were created in R v4.5.1 (R Core Team, 2025) using the following packages: ggplot2 v3.5.2 (Wickham, 2016), scales v1.4.0 (Wickham, Pedersen & Seidel, 2020), emojifont v0.5.5 (Yu & Ekstrøm, 2021), ggtext v0.1.2 (Wilke & Wiernik, 2022), and patchwork v1.3.0 (Pedersen, 2025). Figure 5 was created in Inkscape (https://inkscape.org/).

## Supporting Information

**Appendix S1.** Web of Science and Google Scholar search queries, Google Scholar search results and PRISMA 2020 flow diagrams.

## Acknowledgements

CC acknowledges from the support of the Portuguese Foundation for Science and Technology (FCT) through InvaSTOP project grant (https://doi.org/10.54499/2023.12533.PEX) and funds to CEG/IGOT Research Unit (UIDB/00295/2020 and UIDP/00295/2020). RP acknowledges funding from FCT (PRT/BD/153694/2021; https://doi.org/10.54499/PRT/BD/153694/2021) for funds to GHTM - UIDB/04413/2020 & UIDP/04413/2020) and would like to thank the AIR Centre for their support. BL was funded by his salary as a French public servant.

