## Appendix 1 - Supplementary Information for "Impacts of human-introduced species on the geography of life on Earth"

### Supporting Information for: Impacts of human-introduced species on the geography of life on Earth

#### Supplementary Tables

**Supplementary Table 1.** Web of Science search queries for each biogeographical pattern and rule, including the base keywords, full search strings used, and links to the corresponding search results.

| Pattern/rule | Main keywords (withouth derivates) | Search query | Query link |
|---|---|---|---|
| Latitudinal gradient in species diversity | "latitudinal gradient", "latitudinal diversity", "latitudinal variation", "biodiversity gradient", "latitudinal pattern" | ALL=(("richness" OR diversit*) AND ("latitud* gradient*" OR "latitud* variation*" OR latitud* pattern*)) AND ALL=("biological invasion*" OR "invasive species" OR "non$native species" OR "alien species" OR "introduced species" OR "exotic species" OR "allochthonous species" OR "non$indigenous species" OR "species invasion*" OR "species introduction*") | https://www.webofscience.com/wos/woscc/summary/11842cf1-5aed-402d-90e6-fb6476b0d661-f69a1b87/relevance/1 |

| | | | |
|---|---|---|---|
| Biogeographical regions | "biogeographical regions, biogeographic zones", "phylogenetic regions", "biogeographic provinces", "taxonomic regions", "zoogeographical regions", "floristic regions", "bioregions", "ecozones", "ecoregions", "biotic provinces", "biotic regions", "species turnover", "taxonomic turnover", "phylogenetic turnover", "beta-diversity", compostional (dis)similarity | ALL=(("biogeographi* region*" OR "biogeographi* zones" OR "phylogenetic region*" OR "biogeographi* provinces" OR "taxonomic* region*" OR "zoogeographic* region*" OR "floristic* region*" OR "bioregion*" OR "ecozone*" OR "ecoregion*" OR "biotic province*" OR "biotic region*") AND ("species turnover" OR "beta$diversity" OR "composition*" OR "similarity" OR "dissimilarity" OR "bio* homogeni?ation" OR "bio* differentiation" OR "change* in species composition" OR "alteration* in species composition" OR "shift* in species pool" OR "change* in species assemblage" OR "modification* in species composition" OR "species composition change*" OR "species assemblage change*" OR "transformation* in species composition")) AND ALL=("biological invasion*" OR "invasive species" OR "non$native species" OR "alien species" OR "introduced species" OR "exotic species" OR "allochthonous species" OR "non$indigenous species" OR "species invasion*" OR "species introduction*") | https://www.webofscience.com/wos/woscc/summary/bfb156d7-c6e5-47e9-b222-ed99789a8c52-f69a73b0/relevance/1 |
| Wallace's line | Wallace's line, "Wallace line","Wallacea" | ALL=("Wallace's line" OR "Wallace line" OR "Wallacea") AND ALL=("biological invasion*" OR "invasive species" OR "non$native species" OR "alien species" OR "introduced species" OR "exotic species" OR "allochthonous species" OR "non$indigenous species" OR "species invasion*" OR "species introduction*") | https://www.webofscience.com/wos/woscc/summary/0289d7c3-f64d-476e-ba2e-5c2bae743b34-f69ac8ce/relevance/1 |
| Species-area relationship | "species-area relationship", "SAR", "species-area curve", "species-area effect", "species-area theory", "area-species scaling", "species accumulation curve" | ALL=("species-area" OR "area-diversity" OR "area-species" OR "species-accumulation" OR "SAR" OR "diversity-area" OR "area-species" OR "area-richness" OR "richness-area") AND ALL=("biological invasion*" OR "invasive species" OR "non$native species" OR "alien species" OR "introduced species" OR "exotic species" OR "allochthonous species" OR "non$indigenous species" OR "species invasion*" OR "species introduction*") | https://www.webofscience.com/wos/woscc/summary/5e1af806-c52c-40c4-8a99-7a69e02330d5-f69a8ee4/relevance/1 |

| | | | |
|---|---|---|---|
| Island diversity-isolation | "island biogeography", "equilibrium theory", "MacArthur-Wilson" | ALL=("island biogeograph*" OR "equilibrium theory" OR "MacArthur-Wilson") AND ALL=("biological invasion*" OR "invasive species" OR "non$native species" OR "alien species" OR "introduced species" OR "exotic species" OR "allochthonous species" OR "non$indigenous species" OR "species invasion*" OR "species introduction*") | https://www.webofscience.com/wos/woscc/summary/616e5853-77fd-41c2-a316-a8a430817323-f69adae1/relevance/1 |
| Distance decay of similarity | "distance decay", "distance effect" | ALL=("distance decay" OR "distance effect*") AND ALL=("biological invasion*" OR "invasive species" OR "non$native species" OR "alien species" OR "introduced species" OR "exotic species" OR "allochthonous species" OR "non$indigenous species" OR "species invasion*" OR "species introduction*") | https://www.webofscience.com/wos/woscc/summary/7f0d899e-046b-418b-8da2-0015fe1ba49b-f69ab153/relevance/1 |
| Bergmann's rule | "Bergmann's rule", "body size and climate", "latitudinal clines in body size", "Bergmann's cline", "temperature-size rule", "climate and body mass", "size-latitude relationship", "body mass and latitude, "size-temperature relationship" | ALL=("Bergmann*" OR "body size and climate" OR "latitudinal clines in body size" OR "temperature-size rule" OR "climate and body mass" OR "size-latitude relationship" OR "body mass and latitude" OR "size-temperature relationship") AND ALL=("biological invasion*" OR "invasive species" OR "non$native species" OR "alien species" OR "introduced species" OR "exotic species" OR "allochthonous species" OR "non$indigenous species" OR "species invasion*" OR "species introduction*") | https://www.webofscience.com/wos/woscc/summary/a8b532cd-bc5c-4efd-bddb-2089f46cf616-f69c01aa/relevance/1 |
| Rapoport's rule | "Rapoport", "geographic range size gradient", "latitudinal range size", "altitudinal range size", "depth range size" | ALL=("rapoport*" OR (("range size*" OR "distribution range*" OR "geographic range*" OR "distribution area*" OR "area* of occupancy" OR "extent of occurrence" OR "species range*") AND ("altitud* gradient*" OR "altitud* variation*" OR "altitud* pattern*" OR "latitud* gradient*" OR "latitud* variation*" OR latitud* pattern* OR "depth* gradient*" OR "depth* variation*" OR "depth* pattern*"))) AND ALL=("biological invasion*" OR "invasive species" OR "non$native species" OR "alien species" OR "introduced species" OR "exotic species" OR "allochthonous species" OR "non$indigenous species" OR "species invasion*" OR "species introduction*") | https://www.webofscience.com/wos/woscc/summary/ecfe660f-eeca-433a-9a94-869427459d88-f69d4296/relevance/1 |

| | | | |
|---|---|---|---|
| Gloger's rule: | | ALL=("Gloger* rule" OR "Gloger* principle" OR "pigmentation" OR "coloration" OR "color variation*" OR "variation in color" OR "climate-driven color") AND ALL=("biological invasion*" OR "invasive species" OR "non$native species" OR "alien species" OR "introduced species" OR "exotic species" OR "allochthonous species" OR "non$indigenous species" OR "species invasion*" OR "species introduction*") | https://www.webofscience.com/wos/woscc/summary/6b3046fb-3308-44ca-a556-eb168a01620d-f69c3871/relevance/1 |
| Foster's rule (Island Rule) | "Foster's Rule", "Island Rule", "insular dwarfism", "insular gigantism", "size evolution on islands", "insular size change", "size scaling on islands", "insular body size" | ALL=("Foster* Rule" OR "Island Rule" OR "dwarfism" OR "gigantism" OR "size evolution" OR "insular size" OR "size scaling") AND ALL=("biological invasion*" OR "invasive species" OR "non$native species" OR "alien species" OR "introduced species" OR "exotic species" OR "allochthonous species" OR "non$indigenous species" OR "species invasion*" OR "species introduction*") | https://www.webofscience.com/wos/woscc/summary/da51f976-e13f-4daa-8614-396c7435e1ca-f69c4965/relevance/1 |
| Body size - range size relationships | "body size-range size relationship", "size-range relationship", "body size and geographic range", "size and range", "range size and body mass", "distribution and body size", "species range and body size", "body mass and range size", "distribution size and body mass", "size-area relationship" | ALL=("size-range" OR "size* and range*" OR "range size and body mass" OR "distribution and body size" OR "range* and body size*" OR "body mass and range size*" OR "size-area" OR "range-size" OR "range* and size*" OR "body mass and range size" OR "body size and distribution" OR "body size* and range*" OR "range size* and body mass" OR "area-size") AND ALL=("biological invasion*" OR "invasive species" OR "non$native species" OR "alien species" OR "introduced species" OR "exotic species" OR "allochthonous species" OR "non$indigenous species" OR "species invasion*" OR "species introduction*") | https://www.webofscience.com/wos/woscc/summary/0fbff551-2873-4314-8445-1c894c193add-f69d0b9a/relevance/1 |

| | | | |
|---|---|---|---|
| Abundance-range size relationships | "abundance-range size relationships", "abundance-range relationship", "distribution-abundance relationship", "occupancy-abundance relationship", "range-abundance scaling", "abundance-area relationship" | ALL=("abundance-range*" OR "distribution-abundance*" OR "occupancy-abundance" OR "range-abundance" OR "abundance-diversity" OR "abundance-area" OR "densit* range*" OR "density-abundance*" OR "occupancy-density" OR "range-density" OR "density-diversity" OR "density-area" OR "abundance-richness" OR "distribution-richness" OR "occupancy-richness" OR "range-richness" OR "richness-area" OR "densit* richness*" OR "density-richness") AND ALL=("biological invasion*" OR "invasive species" OR "non$native species" OR "alien species" OR "introduced species" OR "exotic species" OR "allochthonous species" OR "non$indigenous species" OR "species invasion*" OR "species introduction*") | https://www.webofscience.com/wos/woscc/summary/836fd016-476d-438c-9ed9-e34be515bc8b-f69b4786/relevance/1 |
| Species abundance distributions | "species abundance distributions", "SAD" , "abundance distributions", "species abundance patterns", "relative species abundance", "log-normal distribution", "abundance-diversity relationship" | ALL=("abundance distributions*" OR "abundance pattern*" OR ("log-normal distribution" AND "macroecolog*") OR "abundance-diversity" OR "densit* distributions*" OR "densit* pattern*" OR "density-diversity"OR "SAD") AND ALL=("biological invasion*" OR "invasive species" OR "non$native species" OR "alien species" OR "introduced species" OR "exotic species" OR "allochthonous species" OR "non$indigenous species" OR "species invasion*" OR "species introduction*") | https://www.webofscience.com/wos/woscc/summary/c2626427-eb72-44c2-845f-48c84714f30e-f69b2bb6/relevance/1 |
| Body size distributions | "body size distributions", "size distribution patterns", "body size frequency", "size structure", "body size variation", "L-shaped", "size-frequency distribution", "body size patterns" | ALL=("size distribution*" OR "size frequenc*" OR "size structure" OR "size variation" OR "L-shaped" OR "size pattern*") AND ALL=("biological invasion*" OR "invasive species" OR "non$native species" OR "alien species" OR "introduced species" OR "exotic species" OR "allochthonous species" OR "non$indigenous species" OR "species invasion*" OR "species introduction*") | https://www.webofscience.com/wos/woscc/summary/78839fe3-5b04-417e-aa04-30d18aa53b85-f69d27ba/relevance/1 |

| | | | |
|---|---|---|---|
| Species range size distributions | "range size frequency distribution", "range size distributions" | ALL=(("range size*" OR "distribution range*" OR "geographic range*" OR "distribution area*" OR "area* of occupancy" OR "extent of occurrence" OR "species range*") AND ("frequenc* distribution*" OR "macroecolog*" OR "frequenc* pattern*")) AND ALL=("biological invasion*" OR "invasive species" OR "non$native species" OR "alien species" OR "introduced species" OR "exotic species" OR "allochthonous species" OR "non$indigenous species" OR "species invasion*" OR "species introduction*") | https://www.webofscience.com/wos/woscc/summary/a5781d04-7612-4118-9828-7761d628ca04-f69d8d0f/relevance/1 |

**Supplementary Text**

**Supplementary Text 1.** Results of Google Scholar searches

Latitudinal Gradient of Species Diversity - Date: 03/04/2025.

Search query: latitudinal gradient invasi*

Plant species **invasions** along the **latitudinal gradient** in the United States

TJ Stohlgren, D Barnett, C Flather, J Kartesz… - Ecology, 2005 - Wiley Online Library

… **latitudinal gradient** for the density of native plant species and a significant, slightly positive **latitudinal gradient** … We found stronger evidence of a significant, positive productivity **gradient** (…

A **latitudinal gradient** of deep-sea **invasions** for marine fishes

ST Friedman, MM Muñoz - Nature Communications, 2023 - nature.com

… we find a strong **latitudinal gradient** in community differences … the depth axis, reflecting repeated **invasions** of new depth zones. … Here, we identify a **latitudinal gradient** for niche lability in …

Plant species **invasions** along the **latitudinal gradient** in the United States: comment

JD Fridley, H Qian, PS White, MW Palmer - Ecology, 2006 - JSTOR

… documented as the **latitudinal gradient** in species richness. … as to whether clear **latitudinal** patterns are manifest at … attempt to address the **latitudinal gradient** of plant richness for the full …

**Latitudinal** patterns of alien plant **invasions**

Q Guo, BS Cade, W Dawson, F Essl… - Journal of …, 2021 - Wiley Online Library

… as well as the subset of all invasive plants in each regional flora (such as countries, states or provinces) along a full **latitudinal gradient** from tropical to polar regions across continents. …

**Latitudinal gradients** and geographic ranges of exotic species: implications for biogeography

DF Sax - Journal of Biogeography, 2001 - Wiley Online Library

… biogeography, a synthetic explanation for the **latitudinal gradient** in diversity, and an examination of species richness on oceanic islands following the **invasion** of exotic species. …

Biogeography of a plant **invasion**: drivers of **latitudinal** variation in enemy release

WJ Allen, LA Meyerson, D Cummings… - Global ecology and …, 2017 - Wiley Online Library

… Our aim was to compare **latitudinal gradients** in herbivory between native and
… **latitudinal**
**gradients** in herbivory between native and invasive plants and to investigate
whether **gradients** …

Incumbency, diversity, and **latitudinal gradients**

JW Valentine, D Jablonski, AZ Krug, K Roy - Paleobiology, 2008 - cambridge.org

… The result is a **gradient** wherein the ma jority of taxa in each **latitudinal** bin is shared with the … Over the past few centuries, successful species **invasions** have been less frequent in …

**[PDF]** Is there a **latitudinal gradient** in the importance of biotic interactions?

DW Schemske, GG Mittelbach, HV Cornell… - Annual Review of …, 2009 - academia.edu

… keywords (eg, **latitudinal gradient**, biotic interactions, … **latitudinal gradient**. More
common
are studies that compare single localities from high and low latitudes, or that
examine **latitudinal** …

Biogeography of a plant **invasion**: genetic variation and plasticity in **latitudinal** clines for traits related to herbivory

GP Bhattarai, LA Meyerson, J Anderson… - Ecological …, 2017 - Wiley Online Library

… **invasions** with **latitudinal gradients** in herbivore pressure is an important yet mostly
unexplored
issue in **invasion** … a **latitudinal gradient**, which can contribute to spatial heterogeneity in …

Biogeography of a plant **invasion**: plant–herbivore interactions

JT Cronin, GP Bhattarai, WJ Allen, LA Meyerson - Ecology, 2015 - Wiley Online Library

… should exhibit **latitudinal gradients** in the strength of their interactions with herbivores. We hypothesize that if an invasive plant species exhibits a different **latitudinal gradient** in response …

Save Cite Cited by 149 Related articles All 17 versions

**Invasion** by alligator weed, *Alternanthera philoxeroides*, is associated with decreased species diversity across the **latitudinal gradient** in China

H Wu, J Carrillo, J Ding - Journal of Plant Ecology, 2016 - academic.oup.com

… of alien **invasion** and its interaction with **latitudinal gradients** of biodiversity on … **invasion** by A. philoxeroides is associated with decreased species diversity across the **latitudinal gradient** …

Save Cite Cited by 45 Related articles All 5 versions

**[HTML]** springer.com

**[HTML]** **Latitudinal** shifts of introduced species: possible causes and implications

Q Guo, DF Sax, H Qian, R Early - Biological **Invasions**, 2012 - Springer

… global lowest **latitudinal** limit were … **gradients**. Both regions are located in the northern hemisphere and have tropical, subtropical, temperate, and boreal areas along a **latitudinal gradient**…

**[HTML]** **Latitudinal** trends in growth, reproduction and defense of an invasive plant

L Xiao, MR Hervé, J Carrillo, J Ding, W Huang - Biological **invasions**, 2019 - Springer

… plants allocate resources along **latitudinal gradients** from a whole plant perspective. … A recent study suggests that the successful **invasion** of P. americana may result from a lack of …

**Latitudinal** directionality in ectotherm **invasion** success

P Amarasekare, MW Simon - Proceedings of the …, 2020 - royalsocietypublishing.org

… are rare, as are re-**invasions** of the tropics by taxa adapted to … **Latitudinal** directionality in **invasion** success is also observed … the evolutionary play of **latitudinal** diversity **gradients** unfolds. …

[**Latitudinal** variation in the diversity and composition of various organisms associated with an exotic plant: the role of climate and plant **invasion**]

L Gao, C Wei, H Xu, X Liu, E Siemann, X Lu - New Phytologist, 2021 - Wiley Online Library

… (the distance decay of similarity) with **latitudinal** distance (Nekola & White, 1999; Bahram et al., 2013). These **latitudinal gradients** serve as natural experiments for uncovering the …

**[HTML]** Life history and parasites of the invasive mosquitofish (*Gambusia holbrooki*) along a **latitudinal gradient**

L Benejam, C Alcaraz, P Sasal, G Simon-Levert… - … **Invasions**, 2009 - Springer

… mosquitofish along a **latitudinal gradient** from southern … **latitudinal** studies of invasive species to understand the interactive mechanisms of climate change and biological **invasions**…

**Latitudinal**-diversity **gradients** can be shaped by biotic processes: new insights from an eco-evolutionary model

R Henriques-Silva, A Kubisch, PR Peres-Neto - Ecography, 2019 - Wiley Online Library

… involved in shaping **latitudinal**-diversity **gradients** (LDGs) … in diversity decline across **latitudinal** **gradients** among groups. In … Our simulation aims at representing the **invasion** of a vacant …

Impact of an invasive alien plant on litter decomposition along a **latitudinal gradient**

K Helsen, SW Smith, J Brunet, SAO Cousins… - …, 2018 - Wiley Online Library

… Our results show that both **invaded** and non-**invaded** plant communities had a higher … the **latitudinal gradient**, partly explaining the variation in decomposition rates along the **gradient**. At …

A **latitudinal gradient** of beta diversity for exotic vascular plant species in North America

H Qian - Diversity and Distributions, 2008 - Wiley Online Library

… for a low value for PC2 in Zone D (Table S2 in Supplementary Material), indicating that different **latitudinal** zones had similar climate variation among floras within each **latitudinal** zone. …

Invasive species: genetics, characteristics and trait variation along a **latitudinal gradient**

KP Acharya - 2014 - ntnuopen.ntnu.no

… The aim of this study is to study the fundamental issues in **invasion** biology: genetics, characteristics of invasive species, and trait variation along a **latitudinal gradient**. A better

**Biogeographical regions (Date: 03/04/2025).**

**Search Terms: Biogeographic* regions invasi***

**[HTML]** Economic drivers of biological invasions: A worldwide, bio-geographic analysis

S Dalmazzone, S Giaccaria - Ecological Economics, 2014 - Elsevier

… incidence of **invasive** species to resource extraction, pollution and to import volumes disaggregated by country and **region** of origin. The model, estimated with data on **invasive** species …

Comparative **biogeography** of marine invaders across their native and introduced ranges

PE Gribben, JE Byers - Oceanography and Marine Biology, 2020 - library.oapen.org

… **biogeography** of marine invasions. Additionally, we catalogue the geographic **regions** of the **invasive** species used in **biogeographic** … is **biogeographical** comparisons of an **invasive** …

A taxonomic, **biogeographical** and ecological overview of **invasive** woody plants

P Binggeli - Journal of vegetation Science, 1996 - Wiley Online Library

… Included in these **regions** are areas which have fewer highly **invasive** species, eg islands on continental shelves, such as the British Isles. Although most invasions occur in disturbed …

A **biogeographical** approach to plant invasions: the importance of studying exotics in their introduced *and* native range

JL Hierro, JL Maron, RM Callaway - Journal of ecology, 2005 - Wiley Online Library

… some issues in **invasion** ecology that do not require an explicit **biogeographical** perspective, … among plants from different **biogeographical regions** have profound implications for plant …

Marine **biogeography** and ecology: invasions and introductions

JC Briggs - Journal of **Biogeography**, 2007 - Wiley Online Library

… communities as being open to **invasion** from areas with greater … **Regional** pools were assembled from published lists and by consulting taxonomic experts. **Biogeographical regions** …

The **biogeography** of naturalization in alien plants

P Pyšek, DM Richardson - Journal of **Biogeography**, 2006 - Wiley Online Library

… the **invasive** status of a species in a given **region** should be based exclusively on measures of population growth and spread; definitions based on such features capture **biogeographical**…

Naturalization of introduced plants: ecological drivers of **biogeographical** patterns

DM Richardson, P Pyšek - New Phytologist, 2012 - Wiley Online Library

… of drivers of **invasion** and to predict **invasion** dynamics, because … in different **regions** and global **biogeographical** patterns of plant … species are therefore **invasive** in some **regions** while in …

**Biogeographical** patterns and determinants of **invasion** by forest pathogens in Europe

A Santini, L Ghelardini, C De Pace… - New …, 2013 - Wiley Online Library

… Spread of endemic species within Europe was the main cause of **invasion** before the 1940s. The **region** of origin of alien pathogens was mainly North America from the early 1940s to …

**[BOOK]** **Biogeography** of Mediterranean invasions

RH Groves, F Di Castri - 1991 - books.google.com

… the challenge to all biological scientists to better understand the **invasion** process as it is worked out by many plants and animals in **regions** of mediterranean climate. If this book helps …

Naturalized alien flora of the world: species diversity, taxonomic and phylogenetic patterns, geographic distribution and global hotspots of plant **invasion**.

P Pyšek, J Pergl, F Essl, B Lenzner, W Dawson, H Kreft… - 2017 - digital.csic.es

… of **invasion** ecology as a distinct research field, understanding the macroecological and **biogeographic** … of naturalized, **invasive** and native species in particular **regions** of the world (…

Species invasions and the changing **biogeography** of Australian freshwater fishes

JD Olden, MJ Kennard, BJ Pusey - … Ecology and **Biogeography**, 2008 - Wiley Online Library

… found that **invasive** species have significantly changed the present-day **biogeography** of fish … in beta-diversity among **regions** with very different **invasion** histories and contrasting …

**Biogeography** of a plant **invasion**: plant–herbivore interactions

JT Cronin, GP Bhattarai, WJ Allen, LA Meyerson - Ecology, 2015 - Wiley Online Library

… **invasion** at lower than at higher latitudes. Our study points to the need for **invasion** biology to
include a **biogeographic** … 1 in which an **invasive** plant species comes from a **region** where it …

Latitudinal gradients and geographic ranges of exotic species: implications for **biogeography**

DF Sax - Journal of **Biogeography**, 2001 - Wiley Online Library

… **region** than another, the explanation cannot be due to differential rates of speciation between **regions**, … on 'The paradox of **invasion**' in Global Ecology and **Biogeography**. He is currently …

The **biogeography** of **invasive** alien plants in California: an application of GIS and spatial regression analysis

SJ Dark - Diversity and Distributions, 2004 - Wiley Online Library

… The SAR model for **invasive** alien plants included three significant effects; elevation, road … as **invasive** alien plants. Both **invasive** and noninvasive alien plants are found in **regions** with …

The **biogeography** of prediction error: why does the introduced range of the fire ant over-predict its native range?

MC Fitzpatrick, JF Weltzin, NJ Sanders… - … and **biogeography**, 2007 - Wiley Online Library

… occurrences under-predicted the **invasive** potential of fire ants, … understanding of the **biogeography** of **invasive** and native … techniques, more **invasive** species and different **regions** of …

**[HTML]** **Invasive** alien plants of Russia: insights from **regional** inventories

…, M Hejda, M van Kleunen, **Regional** Contributors… - Biological …, 2018 - Springer

… , **biogeographic** and ecological characteristics of its **invasive** flora. We also elucidate (3) the … between the **invasive** species composition in different **biogeographic regions** of Russia, and …

**Biogeographic** effects of red fire ant **invasion**

NJ Gotelli, AE Arnett - Ecology Letters, 2000 - Wiley Online Library

… in species richness measured at **regional** scales, whereas we … the range limit of a competitively aggressive **invasive** species. … of the **invasive** fire ant are evident at a large **biogeographic** …

*Prosopis*: a global assessment of the **biogeography**, benefits, impacts and management of one of the world's worst woody **invasive** plant taxa

RT Shackleton, DC Le Maitre, NM Pasiecznik… - AoB plants, 2014 - academic.oup.com

… , and local and **regional** economies in their native and even more so in their **invasive** ranges; … a lack of research for less prominent **invasive** alien species in poorer **regions** of the world. …

**[HTML]** Alien flora of India: taxonomic composition, **invasion** status and **biogeographic** affiliations

AA Khuroo, ZA Reshi, AH Malik, E Weber, I Rashid… - Biological …, 2012 - Springer

… One of these indicators is the number of **invasive** species in a **region**, which can only be … to their stage of **invasion**, and (c) to explore **biogeographic** affiliations of the alien flora of India. …

Ecological patterns and biological invasions: using **regional** species inventories in macroecology

MW Cadotte, BR Murray, J Lovett-Doust - Biological Invasions, 2006 - Springer

… sets to test hypotheses in **invasion** biology. Analysis of **regional** species inventories can … and so likely do not match geographical or **biogeographical regions**. Again, from the Ontario …

**Species area relationships - Date: 03/04/2025.**

**Search Terms: species area relationships invasi***

**[HTML]** Comparing **species–area relationships** of native and exotic **species**

B Baiser, D Li - Biological Invasions, 2018 - Springer

… across spatial scales and exploring the influence of **invasive species** on native biodiversity. Here, we assess published studies to determine if SARs differ between native and exotic …

Disentangling the abundance–impact **relationship** for **invasive species**

BA Bradley, BB Laginhas, R Whitlock, JM Allen… - Proceedings of the …, 2019 - pnas.org

… **species** evenness and diversity than on **species** richness. Our results show that native responses
to **invasion** depend critically on **invasive species**' … –impact **relationships** reveal how IAS …

**Species** interactions–**area relationships**: biological invasions and network structure in **relation** to island **area**

S Sugiura - Proceedings of the Royal Society B …, 2010 - royalsocietypublishing.org

… **relationship** between island **area** and (i) interactions with exotic **species** and (ii) network structure of **species** … 2009 **Invasive** plant integration into native plant–pollinator networks across …

A synthesis of plant **invasion** effects on biodiversity across spatial scales

KI Powell, JM Chase, TM Knight - American journal of botany, 2011 - Wiley Online Library

… **invasive species** across spatial scales by explicitly recognizing how **invasive species** alter **species**-… • Key results: We found a negative **relationship** between the spatial extent of the study …

Scale-dependent effects of habitat **area** on **species** interaction networks: **invasive species** alter **relationships**

S Sugiura, H Taki - BMC ecology, 2012 - Springer

… by **species–area relationships**. However, when associations with exotic **species** were excluded,
the total number of **species** … positively correlated with forest **area** (the highest correlation …

The **invasion** paradox: reconciling pattern and process in **species** invasions

JD Fridley, JJ Stachowicz, S Naeem, DF Sax… - Ecology, 2007 - Wiley Online Library

… We conclude that natively rich ecosystems are likely to be hotspots for exotic **species**, but that reduction of local **species** richness can further accelerate the **invasion** of these and other …

The evolutionary impact of **invasive species**

HA Mooney, EE Cleland - Proceedings of the National Academy of …, 2001 - pnas.org

… of **species** crossing borders there is also a buildup in the **invasive** potential of those nonnative **species** … "Introduced **species**" may stay at a fairly low population size for years and then …

The statistics and biology of the **species-area relationship**

EF Connor, ED McCoy - The American Naturalist, 1979 - journals.uchicago.edu

… In an effort to clarify the **relation**ship between the equilibrium hypothesis … **species-area relationship**, we pose three questions regarding the basis, use, and interpretation of **species-area** …

**[HTML]** The link between international trade and the global distribution of **invasive** alien **species**

MI Westphal, M Browne, K MacKinnon, I Noble - Biological invasions, 2008 - Springer

… **species** across regions have been limited geographically or taxonomically or have not considered economics. We used a global **invasive species** … positive **relationship** between **species** …

Global indicators of biological **invasion**: **species** numbers, biodiversity impact and policy responses

MA McGeoch, SHM Butchart, D Spear… - Diversity and …, 2010 - Wiley Online Library

… control threats from **invasive** alien **species** and the two … **invasive species** and to (2) have management plans in place for major alien **species** that threaten ecosystems, habitats or **species** …

The population biology of **invasive species**

AK Sakai, FW Allendorf, JS Holt… - Annual review of …, 2001 - annualreviews.org

… these **species** is evident; costs of **invasive species** areestimated … Inaddition to economic impacts, **invasive species** have … the impacts ofinvasive **species** on native **species** and community …

**[PDF]** Plant **invasion** across space and time: factors affecting nonindigenous **species** success during four stages of **invasion**.

KA Theoharides, JS Dukes - New phytologist, 2007 - blogs.law.harvard.edu

… **invasion** of nonindigenous plant **species** (NIPS) is an important component of global environmental change. **Invasive** … , growth of **invasive species** can be suppressed by **species** that …

Five potential consequences of climate change for **invasive species**

JJ Hellmann, JE Byers, BG Bierwagen… - Conservation …, 2008 - Wiley Online Library

… , moving into **areas** where they were previously absent. These are all reasons to specify carefully what is meant by an **invasive species**. We define **invasive species** as those taxa that …

**Invasion** of exotic plant **species** in tallgrass prairie fragments

AC Cully, JF Cully Jr, RD Hiebert - Conservation Biology, 2003 - Wiley Online Library

… by exotic plant **species** at 25 tallgrass prairie sites in central … **species** richness and relative cover as measures of **invasion**. Exotic **species** richness and cover were not related to **area** for …

**[BOOK]** **Invasion** biology

MA Davis - 2009 - books.google.com

… we call invasive **species** are really **invasive** populations, since very few **species** are **invasive** … be clear, when I refer to non-native **species**, I am referring to all recently introduced **species**, …

Global threats from **invasive** alien **species** in the twenty-first century and national response capacities

R Early, BA Bradley, JS Dukes, JJ Lawler… - Nature …, 2016 - nature.com

… **species** invasions. We find that one-sixth of the global land surface is highly vulnerable to **invasion**, including substantial **areas** … need for proactive **invasion** strategies in **areas** with high …

Trade, transport and trouble: managing **invasive species** pathways in an era of globalization

PE Hulme - Journal of applied ecology, 2009 - Wiley Online Library

… have established novel pathways for the spread of alien **species**. Increasingly, the science advances underpinning **invasive species** management must move at the speed of commerce. …

Socioeconomic legacy yields an **invasion** debt

F Essl, S Dullinger, W Rabitsch, PE Hulme… - Proceedings of the …, 2011 - pnas.org

… range of alien plant and animal **species**. However, human population … **invasion** debt hypothesis: If lag times between introduction and establishment are short for a majority of **species**…

**Species** diversity, **invasion** success, and ecosystem functioning: disentangling the influence of resource competition, facilitation, and extrinsic factors

JJ Stachowicz, JE Byrnes - Marine ecology progress series, 2006 - int-res.com

… loss of resident **species** increases the likelihood of the establishment of new **species**. In the … to a study of the consequences of **species** loss for **invasion** in order to assess the generality …

Interactive effects of habitat modification and **species invasion** on native **species** decline

RK Didham, JM Tylianakis, NJ Gemmell… - Trends in ecology & …, 2007 - cell.com

… effect of habitat modification on the interaction between **invasive** and native **species**. However,
**species invasion** might also alter the impact of habitat modification on native **species** (eg [ …

**Distance Decay of similarity - Date: 03/04/2025.**

**Search Terms: distance decay invasi***

**Distance Decay** in Vascular Plant Families of Eastern North America

F Levy - Journal of the Torrey Botanical Society, 2024 - meridian.allenpress.com

… **Distance decay**, the … **distance decay** among eight large vascular plant families and examined
the role of exotics and species geographic range size as contributors to **distance decay**. …

Species range size shapes **distance-decay** in community similarity

R Martín-Devasa, S Martínez-Santalla… - Diversity and …, 2022 - Wiley Online Library

… **distance-decay** curves from simulated communities to assess how the species range sizes shape the functional form of **distance-decay** … the **distance-decay** form in an empirical dataset. …

**Distance decay** of similarity among European urban floras: the impact of anthropogenic activities on β diversity

FA La Sorte, ML McKinney, P Pyšek… - Global Ecology and …, 2008 - Wiley Online Library

… of city centres to evaluate **distance decay** of similarity for … **distance decay** patterns, whereas
neophytes were associated with lower compositional similarity and stronger **distance decay** …

The intermediate distance hypothesis of biological invasions

H Seebens, F Essl, B Blasius - Ecology letters, 2017 - Wiley Online Library

… to the inverse of the **distance decay** of the similarity of ecological … However, in contrast to the **distance decay** of ecological … shows that on average **invasion** dynamics follow simple rules…

**Invasion** in patchy landscapes is affected by dispersal mortality and mate-finding failure

JA Walter, AL Firebaugh, PC Tobin, KJ Haynes - Ecology, 2016 - Wiley Online Library

… We evaluated how the **distance-decay** of recapture probabilities differed between forest and matrix and time (days since release) by fitting dispersal kernel equations to release data …

Geographic and temporal **distance–decay** relationships across taxa

TA Dallas, LA Holian, CT Caten - Oikos, 2024 - Wiley Online Library

… relationships to weaken in the face of dispersal barrier reduction or **invasion** of widespread species. On the other hand, local species extirpations driven by land use change or other …

**Distance decay** of similarity among parasite communities of three marine invertebrate hosts

DW Thieltges, MNAD Ferguson, CS Jones, M Krakau… - Oecologia, 2009 - Springer

… We use published field survey data to investigate **distance decay** of similarity in trematode communities from three prominent coastal molluscs of the Eastern North-Atlantic: the …

Least-cost transportation networks predict spatial interaction of **invasion** vectors

DAR Drake, NE Mandrak - Ecological Applications, 2010 - Wiley Online Library

… **Distance–decay** models have gained wide popularity … **distance-decay** models to movement
patterns of **invasion** vectors, and, in addition, suggest that broad concepts of **distance decay** …

Extra-regional residence time as a correlate of plant invasiveness: European archaeophytes in North America

FA La Sorte, P Pyšek - Ecology, 2009 - Wiley Online Library

… We examine **distance-decay** patterns for each of the three … estimate the probability of **distance-decay** patterns for European … study suggest that **invasion** history and **invasive** potential …

[Estimating dispersal and predicting spread of nonindigenous species]

[JR Muirhead], AM Bobeldyk… - … of **invasive** species …, 2009 - books.google.com

… apply to these stages of the **invasion** sequence. We address … management strategies for **invasive** population spread where … is a distance coefficient, or **distance-decay** parameter, which …

**[HTML]** [Environment and dispersal paths override life strategies and residence time in determining regional patterns of **invasion** by alien plants]

[JR Vicente], [HM Pereira], [CF Randin]… - Perspectives in Plant …, 2014 - Elsevier

… which all alien **invasive** species were recorded … **invasion** and therefore the pool of species reaching each site. As geographic distances (eg ecotones and roads) tend to explain **invasion** …

[Exotic plant **invasion** in agricultural landscapes: A matter of dispersal mode and disturbance intensity]

[F Boscutti], M Sigura, S De Simone… - Applied Vegetation …, 2018 - Wiley Online Library

… For each habitat and plant trait combination, **distance-decay** of similarity was assessed by … less dispersal-limited than native species showing a weaker **distance-decay** of similarity. …

**[PDF]** [Different levels of disturbance influence the distributional patterns of native but not]

F Mologni - Frontiers, 2022 - researchgate.net

… Third, I used generalized linear models to investigate **distance-decay** relationships inside … Coutts, SR, Helmstedt, KJ & Bennett, JR (2018) **Invasion** lags: the stories we tell ourselves and …

[Drivers of **distance-decay** in bryophyte assemblages at multiple spatial scales: Dispersal limitations or environmental control?]

C Cacciatori, [E Tordoni], [F Petruzzellis]… - Journal of …, 2020 - Wiley Online Library

… in **distance decay** of compositional similarity in ecology, the scale dependence of geographical
versus environmental control on **distance decay** … limitations on **distance decay** patterns of …

Different levels of disturbance influence the distributional patterns of native but not exotic plant species on New Zealand small islands

F Mologni - Frontiers of Biogeography, 2022 - escholarship.org

… Third, I used generalized linear models to investigate **distance-decay** relationships inside … Coutts, SR, Helmstedt, KJ & Bennett, JR (2018) **Invasion** lags: the stories we tell ourselves and …

Seagrass-associated fungal communities show **distance decay** of similarity that has implications for seagrass management and restoration

BJ Wainwright, GL Zahn, J Zushi, NLY Lee… - Ecology and …, 2019 - Wiley Online Library

… We show a significant pattern of **distance decay**, with samples collected close to each other having more similar fungal communities in comparison with those that are more distant, …

Drivers of species turnover vary with species commonness for native and alien plants with different residence times

G Latombe, DM Richardson, P Pyšek, T Kučera… - Ecology, 2018 - Wiley Online Library

… **Invasive** alien species pose a major threat to biodiversity. A large proportion of research on … Although both EF and IBD can contribute to the ubiquitous **distance decay** of compositional …

Drivers and implications of **distance decay** differ for ectomycorrhizal and foliar endophytic fungi across an anciently fragmented landscape

EA Bowman, AE Arnold - The ISME Journal, 2021 - academic.oup.com

… **distance decay** as a function of dispersal limitation, but that foliar endophytes would demonstrate **distance decay** as a … We first measured **distance decay** in these fungal communities …

**Distance-decay** relationships partially determine diversity patterns of phyllosphere bacteria on Tamrix trees across the Sonoran Desert

OM Finkel, AY Burch, T Elad, SM Huse… - Applied and …, 2012 - journals.asm.org

… This measure, the “**distance-decay** relationship,” is also the topic of the present study, in … While many Tamarix species are highly **invasive**, T. aphylla has limited sexual reproduction in …

**Distance-decay** effect in stone tool transport by wild chimpanzees

LV Luncz, T Proffitt, L Kulik… - Proceedings of the …, 2016 - royalsocietypublishing.org

… Panda nut-cracking sites, we tested for a **distance-decay** effect, in which the weight of material
… Where hominin landscapes have discrete material sources, a **distance-decay** effect, and …

**Wallace Line - Date: 03/04/2025.**

**Search Terms: Wallace Line invasi***

Crossing the (**Wallace**) **line**: local abundance and distribution of mammals across biogeographic barriers

JF Brodie, O Helmy, M Pangau-Adam, G Ugiek… - Biotropica, 2018 - Wiley Online Library

… We surveyed eight study areas to the west of the **Wallace line** and six areas to the east (Fig. …
**Wallace Line** is a much stronger barrier for animals than for plants. Forests across the **line** …

**[PDF]** Monitors, mammals and **Wallace's Line**

SS Sweet, ER Pianka - Mertensiella, 2007 - zo.utexas.edu

… Because juveniles of large monitor species can survive in communities with placental carnivores to the west of **Wallace**’s **Line**, we speculate that predation rather than resource …

Human influences on vertebrate zoogeography: animal translocation and biological invasions across and to the east of **Wallace's Line**

TE Heinsohn - Faunal and floral migrations and evolution in SE …, 2001 - books.google.com

… of the relative natural zoogeographic integrity of **Wallace's Line**, the transition zone of Wallacea,
… The initial human **invasion** of a pristine Wallacea and Meganesia (the combined New …

Integrating phylogenetic and taxonomic evidence illuminates complex biogeographic patterns along Huxley's modification of **Wallace's Line**

JA Esselstyn, CH Oliveros, RG Moyle… - Journal of …, 2010 - Wiley Online Library

… Huxley originally noted transitional elements in Palawan's fauna; we therefore suggest that his modification of **Wallace**'s **Line** should be recognized as a filter zone, reflecting both his …

… systematics of the *Rana signata* complex of Philippine and Bornean stream frogs: reconsideration of Huxley's modification of **Wallace's Line** at the Oriental …

RM Brown, SI Guttman - Biological Journal of the Linnean …, 2002 - academic.oup.com

… of **Wallace's Line** (as modified by Huxley) include exceptions to an otherwise discrete faunal separation. These results suggest the need for revision of this biogeographical barrier, …

Phylogenetic relationships of the globally distributed freshwater prawn genus *Macrobrachium* (Crustacea: Decapoda: Palaemonidae): biogeography, taxonomy and …

NP Murphy, CM Austin - Zoologica Scripta, 2005 - Wiley Online Library

… There is evidence for a possible effect of **Wallace's Line** at the … **Wallace's Line**. It may be that they predate the existence of … traits associated with the **invasion** of freshwater habitats are …

Notes on the profile of Indonesian **invasive** alien plant species

SS Tjitrosoedirjo - Biotropia, 2007 - journal.biotrop.org

… found at the east part of the **Wallace line**. The species that is found at the eastern part might become alien at the west of the **Wallace line** and vise versa including the **invasive** species. …

Dipterocarp biology as a window to the understanding of tropical forest structure

PS Ashton - Annual Review of Ecology and Systematics, 1988 - JSTOR

… Like dipterocarps, few Fagaceae appear east of **Wallace's line**. Fagaceae are absent from peninsular … There, dipterocarps establish through later **invasion**, following a mast year and the …

**A simple, non-invasive marker of gastric damage: sucrose permeability**

LR Sutherland, M Verhoef, JL Wallace… - The Lancet, 1994 - Elsevier

Disaccharides do not cross intact gastrointestinal mucosa to any appreciable extent unless there is damage to the epithelium. Furthermore, since sucrose is rapidly broken down in the …

Centrosome abnormalities and chromosome instability occur together in pre-**invasive** carcinomas

GA Pihan, J **Wallace**, Y Zhou, SJ Doxsey - Cancer research, 2003 - aacrjournals.org

… of cell cycle control in **invasive** carcinoma. Others have suggested … To address this issue, we examined pre-**invasive** human … Because most pre-**invasive** lesions are not uniformly mutant …

Taxonomic revision of the humpback dolphins (*Sousa* spp.), and description of a new species from Australia

TA Jefferson, HC Rosenbaum - Marine Mammal Science, 2014 - Wiley Online Library

… -Pacific humpback dolphin by a wide distributional gap that coincides with **Wallace's Line**. …
It often has an oval **invasion** of light color from below, at the level of the dorsal fin. Australian …

Orchid historical biogeography, diversification, Antarctica and the paradox of orchid dispersal

TJ Givnish, D Spalink, M Ames, SP Lyon… - Journal of …, 2016 - Wiley Online Library

… Asia and Australia at **Wallace's Line**, or intermittently separate … of epiphytism; and (3) **invasion**
of the northern Andes by the … epiphytism and (3) **invasion** of an extensive tropical cordillera…

Biogeography of the Indo-Australian archipelago

DJ Lohman, M de Bruyn, T Page… - Annual Review of …, 2011 - annualreviews.org

… **Wallace's Line** demarcates the most abrupt faunal transition in the world. To a seasoned … Nevertheless, **Wallace's Line** has remained a heuristic concept and is still widely cited because …

**[HTML]** A demographic model for Palaeolithic technological evolution: the case of East Asia and the Movius **Line**

SJ Lycett, CJ Norton - Quaternary International, 2010 - Elsevier

… Thus, the Movius **Line** – as is the case with its namesake '**Wallace's line**' – must be examined
in terms of its biogeographical context, if the divergent evolutionary trajectories of entities …

Evaluating trans-Tethys migration: an example using acrodont lizard phylogenetics

JR Macey, JA Schulte, A Larson… - Systematic …, 2000 - academic.oup.com

… Asia and Australia-New Guinea, known as **Wallace's line**: (1) primary vicariance caused by … Drifting microcontinents may introduce **invasive** faunal elements to megacontinents in the …

Neo-adjuvant (pre-emptive) cisplatin therapy in **invasive** transitional cell carcinoma of the bladder

DMA **Wallace**, D Raghavan, KA Kelly… - British journal of …, 1991 - Wiley Online Library

… Developments in surgical and radiotherapy techniques for the treatment of localised muscle **invasive** bladder cancer have produced little improvement in overall survival. The presence …

Non-**invasive** and quantitative near-infrared haemoglobin spectrometry inthe piglet brain during hypoxic stress, using a frequency-domainmultidistance instrument

…, JR Ballesteros, S Fantini, D **Wallace**… - Physics in medicine …, 2001 - iopscience.iop.org

… defined by ln(Racr2) versus r, and S is the slope of the **line** … Idc, Iac and I define the intercepts
of the **lines** and are functions … In this study, a compromise solution was used (**Wallace** et al …

**[BOOK]** Encyclopedia of islands

RG Gillespie, D Clague - 2009 - books.google.com

" An exceptionally concise and well-organized compilation of lucid accounts of the historical background and current research into all aspects of island science. Anyone with a serious …

Terahertz Pulsed Imaging of Human Breast Tumors[1]

AJ Fitzgerald, VP **Wallace**, M Jimenez-Linan, L Bobrow… - Radiology, 2006 - pubs.rsna.org

… Findings of this study demonstrate the potential of terahertz pulsed imaging to depict both **invasive** breast carcinoma and ductal carcinoma in situ under controlled conditions and …

Expression of microRNAs and protein-coding genes associated with perineural **invasion** in prostate cancer

RL Prueitt, M Yi, RS Hudson, TA **Wallace**… - The …, 2008 - Wiley Online Library

… Perineural **invasion** (PNI) is the dominant pathway for local **invasion** in prostate cancer. To date, only few studies have investigated the molecular differences between prostate tumors …

**Island Biogeography Theory - Date: 04/04/2025.**

**Search Terms: Island Biogeography Invasi***

On the **island biogeography** of aliens: a global analysis of the richness of plant and bird species on oceanic **islands**

TM Blackburn, S Delean, P Pyšek… - … and **Biogeography**, 2016 - Wiley Online Library

… We might expect quantitatively similar relationships in the **island biogeography** of native and
alien species if assemblages of both sets of species are responding to common structuring …

A roadmap to plant functional **island biogeography**

J Schrader, IJ Wright, H Kreft, M Westoby - Biological Reviews, 2021 - Wiley Online Library

… In Section E, we discuss how global change in form of rising sea levels, habitat destruction and **invasive** species threatens native **island** plants and how functional ecology can be used …

New directions in **island biogeography**

AMC Santos, R Field, RE Ricklefs - … Ecology and **Biogeography**, 2016 - Wiley Online Library

… To these, we add the important question of how **invasive** species affect ecosystem functioning: do they replace native species' functions, add functions not previously performed or …

A roadmap for **island** biology: 50 fundamental questions after 50 years of *The Theory of **Island Biogeography***

J Patiño, RJ Whittaker, PAV Borges… - … of **Biogeography**, 2017 - Wiley Online Library

… partially overlapping **island** topics, including: (Macro)Ecology and **Biogeography**, (… ); **island**
ontogeny and past climate change (4); **island** rules and syndromes (3); **island biogeography** …

Genetic and phylogenetic consequences of **island biogeography**

KP Johnson, FR Adler, JL Cherry - Evolution, 2000 - Wiley Online Library

… of **island biogeography**. We derive a population genetic model of **island biogeography** that incorporates **island** … the mainland, and extinction of **island** populations. The model provides a …

[The **island biogeography** of exotic bird species]

[TM Blackburn], [P Cassey]… - … and **Biogeography**, 2008 - Wiley Online Library

… to studies on both exotic **island biogeography** and the invasibility of native communities. … Julie Lockwood has interests in **invasion** ecology, the role of non-indigenous species in …

[Oceanic **islands** and **biogeographical** theory: a review]

JD Sauer - Geographical Review, 1969 - JSTOR

… **ISLANDS** AS UNITS In spite of some durable controversies, modern theory of **island biogeography** … terial, they have been extremely vulnerable to **invasion** by exotics introduced by …

[Equilibrium theory of **island biogeography** and ecology]

[DS Simberloff] - Annual review of Ecology and Systematics, 1974 - JSTOR

… , is made more sophisticated by assuming different **invasion** and colonization capabilities for different species. First, it is clear from even the simplest equilibrium model that distant …

[Marine **biogeography** and ecology: invasions and introductions]

JC Briggs - Journal of **Biogeography**, 2007 - Wiley Online Library

… as being open to **invasion** from areas with greater biodiversity. … or less dependent on the **invasion** of species from elsewhere. … does not indicate that **invasive** species have caused native …

[The **biogeography** of globally threatened seabirds and **island** conservation opportunities]

[DR Spatz], [KM Newton], R Heinz, [B Tershy]… - Conservation …, 2014 - Wiley Online Library

… **islands** with the 359 **islands** where **invasive** species status was known; this increased the number of **islands** considered to have **invasive** … -occur with **invasive** species on an **island** was …

**[BOOK]** The theory of **island biogeography**

RH MacArthur, EO Wilson - 2001 - books.google.com

… It was my privilege to collaborate with Robert (he preferred not to be called Bob) on **island biogeography**, arguably the piece of work for which he is best and most favorably …

Testing theories of **island biogeography**

TJ Case, ML Cody - American Scientist, 1987 - JSTOR

… **island** communities might be altered by coevolutionary adjustments such that eventually the **island** becomes resistant to **invasion**… steadily with time as **island** assemblages become more …

The biology of insularity: an introduction

DR Drake, CPH Mulder, DR Towns… - … of **biogeography**, 2002 - Wiley Online Library

… by uncritically applying **island biogeography** theory to the … processes, **invasive** alien species
and habitat fragmentation. … -reported phenomenon: **invasive** alien animals that increase …

**Island biogeography** of remote archipelagoes

RG Gillespie, BG Baldwin - The theory of **island biogeography** …, 2010 - degruyter.com

… results have intriguing parallels to the ETIB, and lay a foundation for developing hypotheses to test the predictability of species accumulation, extinction, and **invasion** on remote **islands**. …

Remoteness promotes biological invasions on **islands** worldwide

D Moser, B Lenzner, P Weigelt, W Dawson… - Proceedings of the …, 2018 - pnas.org

… in **island biogeography** is the species–isolation relationship (SIR), a decrease in the number of native species with increasing **island** … on an **island** from the Global Avian **Invasion** Atlas (…

**Invasive** alien species on **islands**: impacts, distribution, interactions and management

JC Russell, JY Meyer, ND Holmes… - Environmental …, 2017 - cambridge.org

… Patterns of **island invasion** are not consistent across SIDS geographic …
in **island biogeography**
and human development. We identify 15 of the most globally prevalent IASs on **islands**. …

Using **island biogeographic** distributions to determine if colonization is stochastic

D Simberloff - The American Naturalist, 1978 - journals.uchicago.edu

… Although the **islands** are so close that **invasion** must frequently occur, their probability of success must be miniscule since no successful **invasion** has been recorded in historic times. If …

**[HTML]** A global comparison of plant invasions on oceanic **islands**

C Kueffer, CC Daehler, CW Torres-Santana… - Perspectives in Plant …, 2010 - Elsevier

… within or between **biogeographic** regions. We present in this … of **invasive** plant species in natural areas of oceanic **islands**. We … **island** groups, we also recorded present but not **invasive** …

The **biogeography** of naturalization in alien plants

P Pyšek, DM Richardson - Journal of **Biogeography**, 2006 - Wiley Online Library

… the **invasive** status of a species in a given region should be based exclusively on measures of population growth and spread; definitions based on such features capture **biogeographical**…

A global picture of biological **invasion** threat on **islands**

C Bellard, JF Rysman, B Leroy, C Claud… - Nature Ecology & …, 2017 - nature.com

… **biogeographical** approach to represent connections between **islands**, IAS-threatened species and **invasive** … Indeed, native and **invasive** species do not respect political boundaries, but …

**Species Abundance Distributions – Date: 04/04/2025.**

**Search Terms: "abundance distribution*" invasi***

The temporal **abundance-distribution relationship** in a global invader sheds light on species distribution mechanisms

C Ewers, M Normant-Saremba… - Aquatic …, 2023 - repozytorium.bg.ug.edu.pl

… abundance fluctuations of **invasive** species offer unprecedented … However, the abundance of **invasive** species is rarely … of the world's 100 worst **invasive** species. Thus for the first time, …

Defining optimal sampling effort for large-scale monitoring of **invasive** alien plants: a Bayesian method for estimating abundance and distribution

C Hui, LC Foxcroft, DM Richardson… - Journal of Applied …, 2011 - Wiley Online Library

… Species-**abundance distribution of** the 167 **invasive** plant species in the Kruger National Park, as estimated from the Bayesian model. The grey bar shows an overwhelming number of …

The relationship between invader abundance and impact

HR Sofaer, CS Jarnevich, IS Pearse - Ecosphere, 2018 - Wiley Online Library

… **invasive** species control strategies. With increasing extent and resolution of datasets of **invasive** … This work illustrated how both the shape of the **abundance distribution and** different …

**Abundance, distribution and** spread of the **invasive** Asian toad *Duttaphrynus melanostictus* in eastern Madagascar

F Licata, GF Ficetola, K Freeman, RH Mahasoa… - Biological …, 2019 - Springer

… the expansion of the **invasive** range of this species, calculates the **invasion** spread rate, and … of the rate of spread, showing a shift of the **invasion** towards the North-West, most probably …

**[HTML]** Using species **abundance distribution models** and diversity indices for biogeographical analyses

S Fattorini, F Rigal, P Cardoso, PAV Borges - Acta oecologica, 2016 - Elsevier

We examine whether Species **Abundance Distribution models** (SADs) and diversity indices can describe how species colonization status influences species community assembly on …

**[PDF]** Abundance and distribution of **invasive** alien plant species in Illu Ababora Zone of Oromia National Regional State, Ethiopia

J Tola, T Tessema - Journal of Agricultural Science and Food …, 2015 - researchgate.net

… The study was conducted to determine the **invasive** alien plant species (IAPS) composition, … on prevention and management measures to combat against **invasive** in the study area. …

Spatiotemporal distribution of an **invasive** insect in an urban landscape: introduction, establishment and impact

SM Thomas, GS Simmons, MP Daugherty - Landscape Ecology, 2017 - Springer

… is to predict potential distribution by generating pseudo-absence such that occurrence (ie presence/pseudo-absence) distribution is comparable to the **abundance distribution pattern**. …

Commonly rare and rarely common: comparing population abundance of **invasive** and native aquatic species

GJA Hansen, MJ Vander Zanden, MJ Blum… - PLoS …, 2013 - journals.plos.org

… We characterized the **abundance distribution of** each … of **invasive** and native species differed
in terms of these statistical moments. We conclude by discussing the implications of **invasive** …

Models of spatiotemporal variation in rabbit abundance reveal management hot spots for an **invasive** species

SC Brown, K Wells, E Roy-Dufresne… - Ecological …, 2020 - Wiley Online Library

… and environmental pest species in its **invasive** range. To better … Long-term monitoring data for **invasive** species can be used … spatial abundance patterns of **invasive** species at the near-…

Spatial heterogeneity in **invasive** species impacts at the landscape scale

AW Latzka, GJA Hansen, M Kornis… - Ecosphere, 2016 - Wiley Online Library

… To generate distributions of impact, we randomly draw an abundance value (α) from an **abundance distribution and** plug it into an abundance–impact curve to calculate impact for that …

**[HTML]** Species-**abundance distribution patterns** of plant communities in the Gurbantünggüt desert, China

Z Zang, Y Zeng, D Wang, F Shi, Y Dong, N Liu, Y Liang - Sustainability, 2022 - mdpi.com

Abstract: It is important to study the species-**abundance distribution pattern** in a community …
model) were used to fit the species-**abundance distribution pattern** of six scales (10 m× 10 m, …

On the species **abundance distribution in** applied ecology and biodiversity management

TJ Matthews, RJ Whittaker - Journal of Applied Ecology, 2015 - Wiley Online Library

… (2005) used a deconstruction approach to look at **invasive** and native species and found that for US birds, on average, **invasive** species obtain higher maximum abundances than native …

Early phase of the **invasion** of *Balanus glandula* along the coast of Eastern Hokkaido: changes in **abundance, distribution, and** recruitment

AKM Rashidul Alam, T Hagino, K Fukaya, T Okuda… - Biological …, 2014 - Springer

… during the early phase of **invasion** along the Pacific coast of … km east of the eastern front of the **invasion** of this species in 2000. … **invasion** dynamics during this early phase of the **invasion**, …

Effect of habitat persistence on the relationship between geographic distribution and local abundance

V Novotný - Oikos, 1991 - JSTOR

… by **invasive** generalists with wide geographic ranges,
the **abundance**/**distribution corr**elation …
the nature of the species **abundance**/ **distribution relationships** is supposed to be essentially …

The **abundance, distribution and** edge associations of six non-indigenous, harmful plants across North Carolina

RW Merriam - Journal of the Torrey Botanical Society, 2003 - JSTOR

Six species of non-indigenous, harmful plants were surveyed throughout North Carolina: Lonicera japonica, Rosa multiflora, Pueraria lobata, Ligustrum sinense, Ailanthus altissima, …

**[HTML]** Distribution and abundance of **invasive** plants in Pacific Northwest forests

A Gray - UNITED STATES DEPARTMENT OF AGRICULTURE …, 2007 - books.google.com

… and **invasion** of … **invasive** species found were Bromus tectorum L., Hypericum perforatum L., and Rubus laciniatus Willd., but several non-native species that are not considered **invasive** …

**[PDF]** Distribution and Abundance

SPIN EXOTIC - marquet.cl

… abundances and show a more even **abundance-distribution rela**tionship, probably as a … of these patterns, we also want to understand the **invasion** process itself, as it offers an unpar…

Disentangling the relationships among abundance, invasiveness and invasibility in trait space

C Hui, P Pyšek, DM Richardson - npj Biodiversity, 2023 - nature.com

… are abundant; the species-**abundance distribution at** fine spatial scales is almost universally … Consequently, invasibility examines all possible **invasion** outcomes for any given **invasive** …

The **abundance, distribution and** diversity of **invasive** and indigenous freshwater snails in a section of the Ogunpa River, southwest Nigeria

MK Oladejo, OO Oloyede, TA Adesakin… - Molluscan …, 2021 - Taylor & Francis

This study investigated the abundance, distribution and diversity of freshwater snails at four sites in the Ogunpa River, Nigeria from May 2018 to December 2018. A total of 2067 …

**Abundance, distribution and** ecological impacts of **invasive** plant species in Maputo Special Reserve, Mozambique

B Sylívér, N Ribeiro, E Cavane… - International Journal of …, 2020 - academicjournals.org

… × 80 m and each **invasive** plants species number were counted … location and coordinates where **invasive** plants species occur … Settlement stratum recorded the highest level of **invasive** …

**Abundance-range size relationship - Date: 04/04/2025.**

**Search Terms: Abundance-range size invasi***

Dimensions of invasiveness: Links between local abundance, geographic range **size**, and habitat breadth in Europe's alien and native floras

TS Fristoe, M Chytrý, W Dawson, F Essl… - Proceedings of the …, 2021 - pnas.org

… **Invasive** alien species pose major threats to biodiversity and ecosystems. However, identifying
drivers of **invasion** … atypical invasions, providing increased clarity into **invasion** processes. …

GIRAE: a generalised approach for linking the total impact of **invasion** to species' range, abundance and per-unit effects

G Latombe, JA Catford, F Essl, B Lenzner… - Biological …, 2022 - Springer

… SAPIA records are, with the exception of the earliest records, geo-referenced to point localities,
but notes were taken as to the abundance of **invasive** plants at the landscape **scale** (and …

Interspecific **abundance-range size** relationships: an appraisal of mechanisms

KJ Gaston, TM Blackburn, JH Lawton - Journal of animal Ecology, 1997 - JSTOR

… This observation would result if ecological flexibility improves the likelihood of successful **invasion**. It might, however, equally be expected from several other hypotheses for the …

Range **size**-abundance relationships in Australian passerines

MRE Symonds, CN Johnson - Global Ecology and …, 2006 - Wiley Online Library

… We instead propose an alternative explanation: that evolutionary age may account for the lack of an **abundance-range size** relationship in Australian passerines. The Passeriformes …

The relationships between **abundance**, **range size** and niche breadth in Central European tree species

B Köckemann, H Buschmann… - Journal of …, 2009 - Wiley Online Library

… Results The relationship between abundance in the distribution centre and range **size** was weak for the Central European tree species. However, significant **abundance–range size** …

**Abundance-range size** relationships in stream vegetation in Denmark

T Riis, K Sand-Jensen - Plant Ecology, 2002 - Springer

… between abundance and range **size** through mechanismsof metapopulation dynamics and … relationship betweenlocal abundance and geographical range **size** of the vascular flora. We …

Testing **abundance-range size** relationships in European carabid beetles (Coleoptera, Carabidae)

DJ Kotze, J Niemelä, RB O'Hara, H Turin - Ecography, 2003 - Wiley Online Library

… In this paper we test four of the eight mechanisms proposed for explaining the **abundance-range size** relationship: the sampling artifact, phylogenetic-non independence, range position …

Commonly rare and rarely common: comparing population abundance of **invasive** and native aquatic species

GJA Hansen, MJ Vander Zanden, MJ Blum… - PLoS …, 2013 - journals.plos.org

… **size** increases. Indeed, the relationship between range **size** and abundance does not differ for **invasive** and native British bird species, but similar to our findings, **invasive** species reach …

Shifting hotspots: Climate change projected to drive contractions and expansions of **invasive** plant abundance habitats

AE Evans, CS Jarnevich, EM Beaury… - Diversity and …, 2024 - Wiley Online Library

… Arrow indicates the average direction of **abundance range** shift from the centroid of current abundance habitat (yellow circle) to the centroid of future abundance habitat (yellow star). …

Adaptive multi-**scale** sampling to determine an **invasive** crab's habitat usage and range in New Zealand

N Gust, GJ Inglis - Biological Invasions, 2006 - Springer

… **invasive** species' impacts on native ecosystems. Here we adaptively increased the spatial **scale** … optimized detection rates prior to larger **scale** geographic surveys that defined the range …

Modeling habitat suitability across different levels of **invasive** plant abundance

EM Beaury, CS Jarnevich, I Pearse, AE Evans… - Biological …, 2023 - Springer

… providing robust sample **sizes** for predicting ranges and how **invasion** population dynamics … of **invasion** risk varies across different thresholds for defining an **abundance range**, and (2) …

On the relationship between species diversity and range **size**

Q Guo, H Qian, J Zhang - Journal of Biogeography, 2022 - Wiley Online Library

… **invasion** biology and conservation. … the **abundance**/**range size** of component species reach a certain level, competition could become stronger and thus constrain species range **size** …

**[HTML]** A broad framework to organize and compare ecological **invasion** impacts

MS Thomsen, JD Olden, T Wernberg, JN Griffin… - Environmental …, 2011 - Elsevier

… ► Few frameworks exist to organize and structure **invasion** impact studies around. ► We propose a framework that separate **invasion** impact into different "impact attributes". ► Impact …

Categorizing **invasive** weeds: the challenge of rating the weeds already in California

JM Randall, N Benton, LE Morse - Weed risk assessment …, 2001 - books.google.com

… Lists of **invasive** wildland weeds can help draw attention to the problem, determine priorities … listing and categorizing **invasive** wildland weeds at a national or regional **scale**. Both cases …

Eight-year record of Hemigrapsus sanguineus (Asian shore crab) **invasion** in western Long Island Sound estuary

GP Kraemer, M Sellberg, A Gordon, J Main - Northeastern Naturalist, 2007 - BioOne

… Full range indicates capture of at least one individual; highest **abundance range** shows distribution where population densities ranged between 30–100% of the maximum value. …

Herbivore regulation of plant abundance in aquatic ecosystems

KA Wood, MT O'Hare, C McDonald… - Biological …, 2017 - Wiley Online Library

… reductions in plant **abundance range** from 0 to 100% … **invasive** plants experience relatively
small losses in biomass due to native herbivores, which may aid the establishment of **invasive** …

**[PDF]** **Invasive** alien species in Northern Bangladesh: identification, inventory and impacts

A Akter, MI Zuberi - International journal of biodiversity and …, 2009 - researchgate.net

… The **Invasive** Species Specialist Group (ISSG) is part of the Species Survival Commission (… on **invasive** species from 41 countries that provides advice on threats from **invasive** and …

Can the impacts of **invasive** species be predicted

M Williamson - Weed risk assessment, 2001 - books.google.com

… of predicting new **invasive** species and their impacts. My interests are in **invasive** species as a … **Abundance-range size** relationships in the herbaceous flora of central England. Journal of …

Using delimiting surveys to characterize the spatiotemporal dynamics facilitates the management of an **invasive** non-native insect

PC Tobin, LM Blackburn, RH Gray, CT Lettau… - Population ecology, 2013 - Springer

… Specifically, within the lowest **abundance range** (0–1 males/trap), there were 376 delimiting … their initial abundance, were more likely to be detected closer to the **invasion** front (Fig. 3c). …

InvasiBES: Understanding and managing the impacts of **Invasive** alien species on Biodiversity and Ecosystem Services

B Gallardo, S Bacher, B Bradley, FA Comín, L Gallien… - NeoBiota, 2019 - hal.science

… continental **scale** can … **scale** as detrimental impacts (eg by quantifying how much native species or human activities benefit from the presence of an **invasive** species), plus a 3-point **scale** …

**Bergmann's rule - Date: 04/04/2025.**

**Search Terms: Bergmann's rule invasi***

Non-native species disrupt the worldwide patterns of freshwater fish body size: implications for **Bergmann's rule**

S Blanchet, G Grenouillet, O Beauchard… - Ecology …, 2010 - Wiley Online Library

… pattern that fits **Bergmann**'s **rule**, with assemblages tending to … , before introduction, **Bergmann**'s **rule** holds for the northern … On the contrary, after introductions, **Bergmann**'s **rule** holds …

**Bergmann's rule** in alien birds

TM Blackburn, DW Redding, EE Dyer - Ecography, 2019 - Wiley Online Library

… If so, we might expect to see **Bergmann's rule** recapitulated in the distributions of alien bird species, for example if small-bodied bird species cannot establish or spread at higher …

**Bergmann's rule** is maintained during a rapid range expansion in a damselfly

C Hassall, S Keat, DJ Thompson… - Global Change …, 2014 - Wiley Online Library

… among samples along the **invasion** route that manifests as a strong, positive correlation between latitude and body size consistent with **Bergmann's rule**. This positive correlation cannot …

Where are we now? **Bergmann's rule** sensu lato in insects

M Shelomi - The American Naturalist, 2012 - journals.uchicago.edu

… **Bergmann**'s **rule** for any group and range of insects is highly idiosyncratic and partially depends on the study design. I conclude that studies of **Bergmann**'s **rule** … its 1970s **invasion** of the …

**Bergmann's** and Allen's **rules** in native European and Mediterranean Phasmatodea

M Shelomi, D Zeuss - Frontiers in Ecology and Evolution, 2017 - frontiersin.org

… For simplicity, we will use the terms "**Bergmann's rule**" to cover body size clines and "Allen's
**rule**" to cover extremity size clines over latitude or altitude regardless of taxon-level, thermal …

[**Bergmann's rule** in ectotherms: a test using freshwater fishes]

MC Belk, [DD Houston] - The American Naturalist, 2002 - journals.uchicago.edu

… The most well known pattern of variation in body size is **Bergmann**'s **rule** (**Bergmann** 1847; Mayr 1956). The intraspecific version of **Bergmann**'s **rule** holds that within endothermic …

[Breaking **Bergmann's rule**: truncation of Northwest Atlantic marine fish body sizes]

JAD Fisher, [KT Frank], [WC Leggett] - Ecology, 2010 - Wiley Online Library

… **Bergmann's rule**. These changes have persisted despite reduced potential for intraspecific competition and favorable bottom water temperatures, both of which should lead to increased …

[Predicting biotic responses to future climate warming with classic ecogeographic **rules**]

[L Tian], [MJ Benton] - Current Biology, 2020 - cell.com

… **Bergmann's rule** is well known, namely that warm-blooded animals are generally … **rules** — Allen's **rule**, Gloger's **rule**, Hesse's **rule**, Jordan's **rule**, Rapoport's **rule** and Thorson's **rule** — …

[Colony size as a buffer against seasonality: **Bergmann's rule** in social insects]

[M Kaspari], EL Vargo - The American Naturalist, 1995 - journals.uchicago.edu

… Colony Size and **Bergmann's Rule** We test a simplified version of **Bergmann's rule** by asking whether tropical species have smaller colony sizes than temperate species. We do this …

[Testing mechanisms of **Bergmann's rule**: phenotypic decline but no genetic change in body size in three passerine bird populations]

[A Husby], SM Hille, [ME Visser] - The American Naturalist, 2011 - journals.uchicago.edu

… for **Bergmann**'s **rule**. Our goals in this study were to provide insight into the mechanisms behind **Bergmann**'s **rule** … According to **Bergmann**'s **rule**, we predicted that great tits would have …

Beyond **Bergmann's rule**: size–latitude relationships in marine Bivalvia world-wide

SK Berke, D Jablonski, AZ Krug, K Roy… - Global Ecology and …, 2013 - Wiley Online Library

Aim Variations in body size are well established for many taxa of endotherms and ectotherms, but remain poorly documented for marine invertebrates. Here we explore how body size …

Tests of ecogeographical relationships in a non-native species: what **rules** avian morphology?

APA Cardilini, KL Buchanan, CDH Sherman, P Cassey… - Oecologia, 2016 - Springer

… clinal patterns in morphology that are consistent with **Bergmann**'s **rule**, Allen's **rule** and Hesse's **rule**. In relation to **Bergmann**'s, Allen's and Hesse's **rules**, respectively, we predict that …

The cold-water connection: **Bergmann's rule** in North American freshwater fishes

AL Rypel - The American Naturalist, 2014 - journals.uchicago.edu

… **Bergmann**'s **rule** and converse **Bergmann**'s **rule** using elevation as the independent variable were rare and observed in only four cases (table 1). Neither of the introduced species (…

**Bergmann's** clines in ectotherms: illustrating a life-history perspective with sceloporine lizards

MJ Angilletta, Jr, PH Niewiarowski… - The American …, 2004 - journals.uchicago.edu

… is the recognition that **Bergmann**'s **rule** has garnered attention … In our opinion, the focus on **Bergmann**'s **rule** as an … Documenting the generality of **Bergmann**'s **rule** is a reasonable …

Natural and sexual selection drive multivariate phenotypic divergence along climatic gradients in an **invasive** fish

XU Ouyang, J Gao, M Xie, B Liu, L Zhou, B Chen… - Scientific Reports, 2018 - nature.com

… **Bergmann**'s **rule**. Finally, Stockwell and Vinyard 58 studied life-history variation of four newly
established (**invasive**) … In this study, we collected **invasive** mosquitofish along 17 degrees of …

The application of ecological **rules** to the racial anthropology of the aboriginal New World

MT Newman - American Anthropologist, 1953 - JSTOR

… Basic to **Bergmann's rule** is the principle that bodies, the larger one has the smaller skin surf … down the Brazilian coast, and the historic **invasion** of the Pampas by Araucanians. The …

Ecogeographical **rules** and the macroecology of food webs

B Baiser, D Gravel, AR Cirtwill… - Global Ecology and …, 2019 - Wiley Online Library

… **Bergmann's rule** applies (interspecific versus intraspecific). The strongest evidence for **Bergmann's rule** … as **Bergmann's rule** (see **Bergmann's rule** section above). Under the island …

Phenotypic variation of an alien species in a new environment: the body size and diet of American mink over time and at local and continental scales

A Zalewski, M Bartoszewicz - Biological Journal of the Linnean …, 2012 - academic.oup.com

… In this way, **invasive** species offer a window into the general … study), a wave of feral mink **invasion** occurred in the early 1990s. … size variation of birds: strong evidence
for **Bergmann's rule** …

Urbanization disrupts latitude-size **rule** in 17-year cicadas

DAE Beasley, CA Penick, NS Boateng… - Ecology and …, 2018 - Wiley Online Library

… in climate, a pattern termed converse **Bergmann's rule**. Urban conditions—particularly warmer … Body size of rural cicadas followed converse **Bergmann's rule**, but this pattern was …

Directionality theory and the evolution of body size

L Demetrius - Proceedings of the Royal Society of …, 2000 - royalsocietypublishing.org

… indicates that the conditions for **invasion** will be determined by … , we can express the **invasion**
conditions uniquely in terms of … of Cope's **rule**, the island **rule** and the **Bergmann** size cline …

**Gloger rule - Date: 04/04/2025.**

**Search Terms: Gloger's rule invasi***

A review of **Gloger's rule**, an ecogeographical **rule** of colour: Definitions, interpretations and evidence

K Delhey - Biological Reviews, 2019 - Wiley Online Library

… **Gloger's rule** and its original definition; (ii) survey the literature to establish how **Gloger's rule**
… the degree of published support for **Gloger's rule** in its different interpretations, summarising …

**[HTML] Gloger's rule**

K Delhey - Current Biology, 2017 - cell.com

What is **Gloger**'s **rule**? **Gloger**'s **rule** describes how variation in animal coloration relates to broad-scale climatic gradients. It was named by Bernhard Rensch in 1929 to honour …

**[PDF]** Do Invaders Conform to Biogeographic "**Rules**"? Testing Allen's, Bergmann's, and **Gloger's Rules** in Monk Parakeets

F Montaño-Centellas, C van Rees, K Block… - Authorea …, 2023 - researchgate.net

… Where collected specimens are available across the native and **invasive** range
… **rules** (Bergmann's,
Allen's, and **Gloger**'s **rules**, see below) with data collected on a widespread **invasive** …

Predicting biotic responses to future climate warming with classic ecogeographic **rules**

L Tian, MJ Benton - Current Biology, 2020 - cell.com

… How **Gloger**'s **rule** operates through the prevalence of pigments requires a complex balance between temperature, humidity and latitude. The simple version of **Gloger**'s **rule** is that all …

Phenotypic divergence despite low genetic differentiation in house sparrow populations

S Ben Cohen, R Dor - Scientific reports, 2018 - nature.com

… validity of this **rule** for birds (42% vs. 72% of bird species 8,9 ). **Gloger**'s **rule** predicts that animals … in **invasion** biology 34,37 . One of the classic textbook study cases of rapid evolution in …

Pig pigmentation: testing **Gloger's rule**

C Newell, H Walker, T Caro - Journal of Mammalogy, 2021 - academic.oup.com

… In this study, we examined the simple and complex versions of **Gloger**'s **rule** in North American wild pigs (also known as wild hogs, wild boar, or feral swine) S. scrofa, descendants of …

Ant assemblages have darker and larger members in cold environments

TR Bishop, MP Robertson, H Gibb… - Global Ecology and …, 2016 - Wiley Online Library

… The two **rules** differ in their target animal groups and in their … **Gloger's rule** is typically applied to endotherms and states … the UV-B protection mechanism of **Gloger's rule** may also be …

Real-time evaluation of glioblastoma growth in patient-specific zebrafish xenografts

E Almstedt, E Rosén, M **Gloger**, R Stockgard… - Neuro …, 2022 - academic.oup.com

… **invasion** of the PDCs and identify cell cultures suited for studies of GBM **invasion** in zebrafish…
Together, our results provide a fast way of assessing GBM tumor initiation capacity, **invasion**…

Rapid evolution of a divergent ecogeographic cline in introduced lady beetles

EM O'neill, EJ Hearn, JM Cogbill, Y Kajita - Evolutionary Ecology, 2017 - Springer

… Specifically, we tested for patterns consistent with **Gloger**'s **rule** and Bogert's **rule** in both native and introduced populations. Spot size, like most quantitative traits can be affected by the …

A population-based longitudinal study on the implications of demographics on future blood supply

…, K Weitmann, A Lebsa, U Alpen, D **Gloger**… - …, 2016 - Wiley Online Library

… In addition, the medical progress is further contributing to less **invasive** treatments with a reduced demand of blood transfusions, for example, by minimal **invasive** heart valve …

Specialtion in the genus Amytornis Stejneger (Passeres: Muscicapidae, Malurinae) in Australia.

A Keast - Australian Journal of Zoology, 1958 - CSIRO Publishing

… increasing aridity, but as a result of the **invasion** of former spinifex or porcupine grass areas by … The application of **Gloger's** and Bergmann's ecogeographical **rules** to members of the …

The island **rule** and a research agenda for studying ecogeographical patterns

MV Lomolino, DF Sax, BR Riddle… - Journal of …, 2006 - Wiley Online Library

… random pattern of range collapse on body-size variation in native or **invasive** species. Similarly,
studies of **invasive** species may provide key insights into the processes that lead to other …

RAF expression in human astrocytic tumors

C Hagemann, J **Gloger**, J Anacker… - International …, 2009 - spandidos-publications.com

… They are highly **invasive** and very difficult to treat, despite of surgery, $\gamma$-irradiation and chemotherapy. Although a role of the mitogenic Ras-RAF-MEK-ERK signalling cascade in brain …

**[PDF]** Towards non-**invasive** biomarkers: intraspecific colour variations in the red wasp Vespula rufa

Q Nguyen - 2023 - erepo.uef.fi

… The colour transformation seems to conform to the ecogeographical **rule** of **Gloger**. In accordance with **Gloger**, species and populations with heavily pigmented forms tend to be found in …

Systematics and evolution of the North American Merlins

SA Temple - The Auk, 1972 - JSTOR

… by Allen's **rule**. The brightness of Merlin plumage color varies as predicted by **Gloger's rule**. … One way to account for this is to suppose a double **invasion** of Merlins from the Palearctic, …

**[CITATION]** The role of competition in extinction: Written discussion of presidential address

JPA Angseesing - Proceedings of the Geologists' Association, 1978 - Elsevier

Biogeographical variation of plumage coloration in the sexually dichromatic Hawai'i 'Amakihi (*Chlorodrepanis virens*)

JM Gaudioso-Levita, PJ Hart, DA LaPointe… - Journal of …, 2017 - Springer

… Our results do not show support for **Gloger**'s **rule**: Hawai'i 'Amakihi had darker plumages in … ), a Monarch Flycatcher, supports **Gloger**'s **rule**. The pale Cs bryani subspecies is found on …

Pigmentation in *Drosophila melanogaster* reaches its maximum in Ethiopia and correlates most strongly with ultra-violet radiation in sub-Saharan Africa

H Bastide, A Yassin, EJ Johanning, JE Pool - BMC Evolutionary Biology, 2014 - Springer

… of the peppered moth [1],[4], and **Gloger's rule** in endotherms and Bogert's **rule** in ectotherms
stating that pigmentation should decrease and increase with latitude, respectively [5]. …

Rapid morphological changes as agents of adaptation in introduced populations of the common myna (*Acridotheres tristis*)

T Magory Cohen, RE Major, RS Kumar, M Nair… - Evolutionary …, 2021 - Springer

… **Invasive** species present an opportunity to test the … made regarding characteristics of **invasive**
populations within a limited range … **rules** of morphological variation (ie, Bergmann **rule** and …

Transcriptomic analysis of skin pigmentation variation in the Virginia opossum (*Didelphis virginiana*)

SF Nigenda-Morales, Y Hu, JC Beasley… - Molecular …, 2018 - Wiley Online Library

… across its distribution range following **Gloger's rule** (lighter pigmentation in temperate … alternative hypotheses that may explain **Gloger's rule** pattern of skin pigmentation variation …

**Foster's rule - Date: 04/04/2025.**

**Search Terms: Foster's rule invasi***

Testing the assumption of environmental equilibrium in an **invasive** plant species over a 130 year history

SL **Foster**, HM Kharouba, TW Smith - Ecography, 2022 - Wiley Online Library

… Using the **invasive** vine Vincetoxicum rossicum, we tested the hypotheses that: 1) **invasive** … equilibrium separately when modelling the distribution of **invasive** species. In light of our …

The varying success of invaders

M Williamson, A Fitter - Ecology, 1996 - JSTOR

… The tens **rule**, showing ratios of the three **invasion** transition stages. The suggested range for the tens **rule** is shown by the black diamond at the top. For British angiosperms (from …

The role of evolution in the **invasion** process

SJ Novak - Proceedings of the National Academy of Sciences, 2007 - pnas.org

… indicating that multiple introductions of **invasive** species may be the **rule** rather than the exception (10). … evidence informing about strategies to reduce xenophobic sentiment and **foster** ... …

Invasibility and compositional stability in a grassland community: relationships to diversity and extrinsic factors

BL **Foster**, VH Smith, TL Dickson, T Hildebrand - Oikos, 2002 - Wiley Online Library

… Our results show that community susceptibility to **invasion** is greatest in high diversity sites … and compositional stability, we indeed cannot **rule** out the possibility that this link is more a …

**[HTML]** Plasmacytoid bladder cancer: variant histology with aggressive behavior and a new mode of **invasion** along fascial planes

…, TA Masterson, KC Cary, JA Pedrosa, RS **Foster**… - Urology, 2014 - Elsevier

… **Invasion** and progression of disease within and along Waldeyer's sheath may be explained … with PCV, with understaging as the **rule** and raised clinical suspicion for advanced disease, …

On the evolution of altruistic ethical **rules** for siblings

TC Bergstrom - The American Economic Review, 1995 - JSTOR

… The relation between the set of equilibrium strategies under Hamilton's **rule** and the sets of strategies that resist **invasion** by dominant and recessive mutants depends critically on …

Mind your own business! Longitudinal relations between perceived privacy **invasion** and adolescent-parent conflict.

ST Hawk, L Keijsers, WW Hale III… - Journal of Family …, 2009 - psycnet.apa.org

… also **foster invasion** perceptions… **rules** have been broken or need to be renegotiated (ie, conflict situations), we predicted that associations from adolescent-father conflict to later **invasion** …

Community disassembly by an **invasive** species

NJ Sanders, NJ Gotelli, NE Heller… - Proceedings of the …, 2003 - pnas.org

… **rules** act to organize these ant communities. The most influential and fiercely debated assembly
**rule** … evidence informing about strategies to reduce xenophobic sentiment and **foster** ... …

Human agency in biological invasions: secondary releases **foster** naturalisation and population expansion of alien plant species

I Kowarik - Biological invasions, 2003 - Springer

… strongly **foster** invasions throughout the whole **invasion** process. … be explained nor predicted
by ecological **rules**. This may be an … In the last section of the paper, the final **invasion** stage …

Is there a global sports law?

K **Foster** - Lex Sportiva: What is Sports Law?, 2012 - Springer

… Hyde, the Australian High Court considered whether a governing body could be held to be negligent when drafting the **rules** of the game or for failing to alter the **rules**. Footnote 12 The …

The community ecology of **invasive** species: where are we and what's next?

L Gallien, M Carboni - Ecography, 2017 - Wiley Online Library

… the search **rule**: (Darwin's naturalization OR assembly **rule***  … OR relatedness) AND (**invasive**
OR alien OR exotic OR native … that **foster** exotic species across their **invasion** stages (…

The effect of public information and competition on trading volume and price volatility

FD **Foster**, S Viswanathan - The Review of Financial Studies, 1993 - academic.oup.com

… decision **rules** does not depend on the specific distribution used. Second, if the model is altered so that the decision to become informed is made endogenous, then the decision **rules** of …

Emergence of spatial structure in cell groups and the evolution of cooperation

CD Nadell, KR **Foster**, JB Xavier - PLoS computational biology, 2010 - journals.plos.org

… (increasing r in Hamilton's **Rule**) can be outweighed by the cost … groups, we performed an **invasion** analysis to determine whether … The results of both our local competition and **invasion** …

Geographic and temporal correlations of mammalian size reconsidered: a resource **rule**

BK McNab - Oecologia, 2010 – Springer

… These patterns are the basis of some geographical and ecological "**rules**." Bergmann (1847) …
The island **rule** (**Foster** 1964; Van Valen 1973) states that with the **invasion** of islands, large …

Rapid spread of **invasive** genes into a threatened native species

BM Fitzpatrick, JR Johnson, DK Kump, JJ Smith… - Proceedings of the …, 2010 - pnas.org

… and fixation of these exceptional **invasive** alleles. The legal status … protection, because different **rules** could result in dramatically … strategies to reduce xenophobic sentiment and **foster** ... …

**[PDF]** Constitutional limitations on corporate activity-protection of personal rights from **invasion** through economic power

AA Berle Jr - U. Pa. L. Rev., 1951 - HeinOnline

… The other area, far less defined, includes a body of **rules** … favors to a group it wished to **foster**
at the expense of the rest of … And this legal restraint would apply not only to **rules** embodied …

**[HTML]** Conservation conundrums and the challenges of managing unexplained declines of multiple species

DB Lindenmayer, J Wood, C MacGregor, C **Foster**… - Biological …, 2018 - Elsevier

… , data from other studies completed both inside Booderee National Park and outside (where intensive fox baiting does not occur yet depleted fauna species remain), allowed us to **rule** …

**[BOOK]** The Oxford illustrated history of Ireland

RF **Foster** - 2000 - books.google.com

… redefining preoccupations and casting a cold eye on **ruling** pieties. All are specialists in their … The same might be said of Katharine Simms's view of the Norman **invasion** of Ireland and …

**[BOOK]** Promoting the **rule** of law abroad: in search of knowledge

T Carothers - 2010 - books.google.com

… Although the United States and other Western countries can and should **foster** the **rule** of law, even large amounts of aid will not bring rapid or decisive results. Thus, it is good that …

**[PDF]** **Invasive** alien species and biodiversity in India

KP Singh - Current Science, 2005 - researchgate.net

… identification and mapping of alien species in their **foster** habitats. This could be attributed to … Algorithm for **Rule**-set Prediction) based modelling in predicting the **invasion** of Prosopis …

**Body size - range size relationships - Date: 04/04/2025.**

**Search Terms: body size range size invasi***

**Body size** and **invasion** success in marine bivalves

K Roy, D Jablonski, JW Valentine - Ecology Letters, 2002 - Wiley Online Library

… more direct approach to test for the role of **body size** in invasions. We focus on the role of **body size** in post-establishment **range** expansion within a given region, thereby circumventing …

Ant **body size** predicts dispersal **distance** of ant-adapted seeds: implications of small-ant invasions

JH Ness, JL Bronstein, AN Andersen, JN Holland - Ecology, 2004 - Wiley Online Library

… be disrupted in habitats dominated by **invasive** ants. We propose that this disruption is related to changes in mean ant **body size**, given that **invasive** ants are smaller than most native …

Biological invasions drive **size** increases in marine and estuarine invertebrates

ED Grosholz, GM Ruiz - Ecology Letters, 2003 - Wiley Online Library

… In this study an unusual pattern of **size** change associated with the **invasion** of 19 species of … any decrease in **size** following **invasion**. This **invasion**-driven increase in **body size** sharply …

Do **invasive** species perform better in their new **ranges**?

JD Parker, ME Torchin, RA Hufbauer, NP Lemoine… - Ecology, 2013 - Wiley Online Library

… **range**, roughly half of the **invasive** species we investigated performed similarly between the home and away **ranges**. … introduced **range** may only partly explain success in a new **range**. …

**Body** mass patterns predict invasions and extinctions in transforming landscapes

CR Allen, EA Forys, CS Holling - Ecosystems, 1999 - Springer

… **body**-mass aggregation patterns. We tested five competing hypotheses by determining whether **invasive** … community **body**-mass patterns. In addition to testing those hypotheses, we …

**Invasive** species grows faster, competes better, and shows greater evolution toward increased seed **size** and growth than exotic non-**invasive** congeners

RC Graebner, RM Callaway, D Montesinos - Plant Ecology, 2012 - Springer

… In sum, our results show greater growth rates and competitive responses for an **invasive** species than its non-**invasive** congeners, and suggest that the **invasive** congener has …

'On being the right **size**'–Do aliens follow the rules?

AAE van der Geer, MV Lomolino… - Journal of …, 2018 - Wiley Online Library

… For each island, we calculated the average **body size measurement** of the focal, introduced … dominate and transform most insular habitats, thus favouring the **invasive** commensals. …

The evolutionary impact of **invasive** species

HA Mooney, EE Cleland - Proceedings of the National Academy of …, 2001 - pnas.org

… evolution of apparently adaptive clines in **body size** and feather color in English sparrows … geographical **range**. Further, Cody and Overton (21) described the reduction in **distance** of …

Progress in **invasion** biology: predicting invaders

CS Kolar, DM Lodge - Trends in ecology & evolution, 2001 - cell.com

… **invasion** process in the future. For example, a species that fails to become entrained in the ballast water of one ship might become entrained in that of another. Also, **invasive** … of **invasion** …

*Drosophila suzukii* (Diptera: Drosophilidae): **Invasive** Pest of Ripening Soft Fruit Expanding its Geographic **Range** and Damage Potential

DB Walsh, MP Bolda, RE Goodhue… - Journal of Integrated …, 2011 - academic.oup.com

… Cooperative projects involving Western states and Canada are underway to address knowledge gaps and prepare recommendations for dealing with this **invasive** new pest. …

**Body size distributions-Date: 04/04/2025.**

**Search Terms: body size distribution* invasi***

Non-native introductions influence fish body size distributions within a dryland river

KJ Fritschie, JD Olden - Ecosphere, 2016 - Wiley Online Library

… composition can also respond to the **body size distribution of** a focal species or community (… However, a more parsimonious interpretation of extinction and **invasion** risk will likely arise …

Climate change and body size shift in Mediterranean bivalve assemblages: unexpected role of biological invasions

R Nawrot, PG Albano… - Proceedings of the …, 2017 - royalsocietypublishing.org

… We show that the **invasion** leads to increase in median body size of the Mediterranean … Mediterranean bivalves, thus shifting the **body size distribution of** the recipient biota toward larger …

Non-native species disrupt the worldwide patterns of freshwater fish body size: implications for Bergmann's rule

S Blanchet, G Grenouillet, O Beauchard… - Ecology …, 2010 - Wiley Online Library

… In this study, we tested if ENNS are a major driver of functional changes (ie changes in **body size distribution)** in freshwater fish assemblages on a worldwide scale. Using information on …

**[HTML]** Evolution of body size, range size, and food composition in a predator–prey metapopulation

C Hui, MA McGeoch - Ecological Complexity, 2006 - Elsevier

… However, species richness and **body size distribution in** different tropic levels have been … and the coarseness or lumpiness of **body-size distribution is** even amplified in higher trophic. …

Evolution, ecology, and multimodal distributions of body size

GS Cumming, TD Havlicek - Ecosystems, 2002 - Springer

… With the single exception of a static, unimodal distribution, the likelihood of detecting consistent
multiple modes in a **body size distribution over** time correlates with the difference in total …

The role of body size variation in community assembly

S Pawar - Advances in ecological research, 2015 - Elsevier

… A necessary (but not sufficient) condition for the **invasion** to … where neither **invasion** (Eq. 10) nor stability (Eq. 11) matter. … the regional species pool's **body size distribution and** size-based …

Variation of morphometric traits in populations of an **invasive** carabid predator (*Merizodus soledadinus*) within a sub-Antarctic island

M Laparie, M Lebouvier, L Lalouette, D Renault - Biological Invasions, 2010 - Springer

… **Invasive** predators may change their own trophic conditions by progressively displacing or reducing diversity and abundance of native prey. As food quality and quantity are two main …

Body-size distributions and size-spectra: universal indicators of ecological status?

OL Petchey, A Belgrano - 2010 - royalsocietypublishing.org

The sizes of individual organisms, rather than their taxonomy, are used to inform management and conservation in some aquatic ecosystems. The European Science Foundation …

**[PDF]** Morphology of **invasion**: body size patterns associated with establishment of Coccinella septempunctata (Coleoptera: Coccinellidae) in western North America

EW Evans - European Journal of Entomology, 2000 - eje.cz

… I also compared **body size distribution of** the invading species with that of native species. The invader was distinctive in having particularly large variation in body size among individuals …

**[HTML]** Non-**invasive** quantification of throat-size distribution and corresponding capillary pressure

AP Garcia, Z Heidari - Journal of Petroleum Science and Engineering, 2021 - Elsevier

… To calculate the pore-**body-size distribution from** Equation 1, we need to relate the surface-to-volume ratio to a characteristic pore scale (eg, radius). In this paper, we consider that the …

**Body size distribution in** flea communities harboured by Siberian small mammals as affected by host species, host sex and scale: scale matters the most

EN Surkova, NP Korallo-Vinarskaya, MV Vinarski… - Evolutionary …, 2018 - Springer

… Studying **body size distribution in** parasites allows us to … We
studied **body size distribution**
**of** fleas parasitic on small … No effect of host sex on the pattern
of **body size distribution was** …

Patch size distribution affects species **invasion** dynamics in dendritic networks

K Holenstein, E Harvey, F Altermatt - Oikos, 2022 - Wiley Online Library

… Competitive hierarchy, **body size distribution and** community dynamics of this set of species
is very well established (Warren 1996, McGrady-Steed et al. 1997, Holyoak and Lawler 2005…

Intraspecific body size frequency distributions of insects

EJ Gouws, KJ Gaston, SL Chown - PLoS One, 2011 - journals.plos.org

Although interspecific body size frequency distributions are well documented for many taxa, including the insects, intraspecific body size frequency distributions (IaBSFDs) are more …

**Body Size Distribution and** Frequency of Anthropogenic Injuries of Bluntnose Sixgill Sharks, *Hexanchus griseus*, at Flora Islets, British Columbia

R Dunbrack, R Zielinski - The Canadian Field-Naturalist, 2005 - canadianfieldnaturalist.ca

… Here we have demonstrated that meaningful population data for such species can be obtained using non**invasive** remote imaging techniques applied at strategically placed observing …

Spawning and brood defense of smallmouth bass under the process of **invasion** into a novel habitat

K Iguchi, T Yodo, N Matsubara - Environmental Biology of Fishes, 2004 - Springer

… of the world's most disastrous **invasive** species, was introduced … to the right in this **invasive** population but rather a skewed normal … decelerate the expansion of **invasive** smallmouth bass. …

Effects of invading vendace (*Coregonus albula* L.) on species composition and body size in two zooplankton communities of the Pasvik River System, northern …

T Bøhn, PA Amundsen - Journal of Plankton Research, 1998 - academic.oup.com

… Abstract Species composition and **body-size distribution were** … A recent **invasion** and successive downstream expansion of … We assumed that the **invasion** and establishment of a …

The population biology of **invasive** species

AK Sakai, FW Allendorf, JS Holt… - Annual review of …, 2001 - annualreviews.org

… In contrast, when theyexamined the combined pattern of vertebrates and invertebrates, they found apositive correlation between mean **body size** and probability ofestablishment. …

**[PDF]** Plant **invasion** across space and time: factors affecting nonindigenous species success during four stages of **invasion**.

KA Theoharides, JS Dukes - New phytologist, 2007 - blogs.law.harvard.edu

… the risk of **invasion** across … **body** of research that suggests that the success of **invasive** NIPS is controlled by a series of key processes or filters. These filters are common to all **invasion** …

**[HTML]** Pore structure characterization of North American shale gas reservoirs using USANS/SANS, gas adsorption, and mercury intrusion

CR Clarkson, N Solano, RM Bustin, AMM Bustin… - Fuel, 2013 - Elsevier

… data allowed pore **size distributions** to be … **invasion** and radiation methods have historically been used for the characterization of shale samples [1]. Knowledge of pore **size distribution** is …

Genetic variation increases during biological **invasion** by a Cuban lizard

JJ Kolbe, RE Glor, L Rodríguez Schettino, AC Lara… - Nature, 2004 - nature.com

… and become **invasive**. To address this issue, we studied the brown anole, a worldwide **invasive** lizard. … Here we show that one key to **invasion** success may be the occurrence of multiple …

**Rapoport Rule - Date: 04/04/2025.**

**Search Terms: rapoport* rule invasi***

Evidence for **Rapoport's rule** and latitudinal patterns in the global distribution and diversity of alien bird species

EE Dyer, DW Redding, P Cassey… - Journal of …, 2020 - Wiley Online Library

Aim To quantify global latitudinal patterns in the distributions of alien bird species to assess whether these species conform to **Rapoport's rule** (ie show a positive latitudinal gradient in …

Latitudinal range variation of trees in the United States: a reanalysis of the applicability of **Rapoport's rule**

CS Lane - The Professional Geographer, 2007 - Taylor & Francis

… Furthermore, in an effort to further verify the applicability of **Rapoport's rule** to US tree taxa, I
… **Rapoport's rule** to North American plant taxa. First, I address the criticism that **Rapoport's rule** …

Altitudinal gradients of species richness and range size of vascular plants in Taiwan: a test of **Rapoport's rule**

W Zhang, Q Lu, J Liang, Z Shen - Biodiversity Science, 2010 - biodiversity-science.net

… **invasive** plants supported **Rapoport**'s **rule**, while that of endemic species and overall species did not. The distribution of pteridophytes supported **Rapoport**'s **rule**, … test of **Rapoport**'s **rule** …

Patterns of turtle species' geographic range size and a test of **Rapoport's rule**

SJ Hecnar - Ecography, 1999 - Wiley Online Library

… Much recent debate lias centered on whctlier **Rapoport's rule**, the tendency lor range size to increase wilh increasing huitude. is a general **rule** or a local etTecl. I calculated the sizes oi' …

The elevational gradient in altitudinal range: an extension of **Rapoport's** latitudinal **rule** to altitude

GC Stevens - The American Naturalist, 1992 - journals.uchicago.edu

… latitudinal range of species and latitude (**Rapoport's** latitudinal **rule**). Both of these **Rapoport** phenomena, the latitudinal and the new elevational **rule** discussed here, can be explained …

Latitudinal gradients in species diversity and **Rapoport's rule** revisited: a review of recent work and what can parasites teach us about the causes of the gradients?

K Rohde - Ecography, 1999 - Wiley Online Library

… conditions are the exception rather than the **rule** among animals. Dispersal abilities of … than at high latitudes (Thorson's **rule**), suggesting that **Rapoport's rule** does not apply to or …

Post-Pleistocene dispersal explains the **Rapoport** effect in North American salamanders

T Radomski, SR Kuchta, KH Kozak - Journal of Biogeography, 2022 - Wiley Online Library

… to the **invasive** range (N = 29). To evaluate our models' abilities to predict populations in the **invasive** … for niche position explaining **Rapoport** effects, but we could not **rule** out the climatic …

Latitudinal gradients and geographic ranges of exotic species: implications for biogeography

DF Sax - Journal of Biogeography, 2001 - Wiley Online Library

… size (**Rapoport**'s **rule**), species geographical range boundaries and patterns of **invasion** in … size, the results of this study predict that **Rapoport**'s **rule** should also be detectable in the …

**[BOOK]** Areography: geographical strategies of species

EH **Rapoport** - 2013 - books.google.com

… presence of mind enabled her to resist unflinchingly the morbid **invasion** of areography. … Esteban and EH **Rapoport** (unpublished results) involving the families Trochilidae, Psittacidae, …

The island **rule** and a research agenda for studying ecogeographical patterns

MV Lomolino, DF Sax, BR Riddle… - Journal of …, 2006 - Wiley Online Library

… **Rapoport's rule** (latitudinal, elevational and bathymetric clines in geographical range size: **Rapoport**, … Similarly, studies of **invasive** species may provide key insights into the processes …

Non-**invasive** detection of respiratory effort-related arousals (RERAs) by a nasal cannula/pressure transducer system

…, AC Krieger, A Rosen, RL O'Malley, DM **Rapoport** - Sleep, 2000 - academic.oup.com

… While this document proposed that non-**invasive** techniques be used for the detection of … desirable to replace it with a more acceptable non-**invasive** technique for detecting RERAs. …

Decreasing species richness with increase in elevation and positive **Rapoport** effects of Crambidae (Lepidoptera) on Mount Taibai

A Chen, Z Li, Y Zheng, J Zhan, B Yang, Z Yang - Insects, 2022 - mdpi.com

… the universality of **Rapoport**'s **rule** in Lepidoptera … **Rapoport**'s **rule** yielded positive results,
while Rohde's results show a unimodal distribution model and do not
support **Rapoport**'s **rule**…

**[CITATION]** Applications of game-theoretic concepts in biology

A **Rapoport** - Bulletin of mathematical biology, 1985 - Springer

Save Cite Cited by 22 Related articles All 5 versions Web of Science: 8

Predicting biotic responses to future climate warming with classic ecogeographic **rules**

L Tian, MJ Benton - Current Biology, 2020 - cell.com

… **rules** — Allen's **rule**, Gloger's **rule**, Hesse's **rule**, Jordan's **rule**, **Rapoport**'s **rule** and Thorson's
**rule** … These **rules** have been discussed in the recent ecological and physiological literature, …

Advances in study of the distribution area of species

Z Wen-Ju, C Jia-Kuan - Biodiversity Science, 2003 - biodiversity-science.net

… ,niche breadth,and ecological **invasion**, but is also connected with … geographic range size(**Rapoport**!s **rule**)sometimes is violated,… ;geographic range relationship,**Rapoport**!s **rule**,and …

Freshwater plant macroecology needs to step forward from the shadows of the terrestrial domain

J Alahuhta, J García-Girón… - Nordia Geographical …, 2025 - nordia.journal.fi

… richness gradient, **Rapoport**'s **rule** and species turnover vs. … Although findings on **Rapoport**'s **rule** are less clear, research … There are six **invasive** freshwater plant species occurring in …

Altitudinal gradients of species richness and range size of vascular plants in Taiwan: a test of **Rapoport's rule**.

ZWJ Zhang WanJun, LQ Lu Qian, LJ Liang Jun… - 2010 - cabidigitallibrary.org

… of **invasive** plants supported **Rapoport's rule**, while that of endemic species and overall species did not. The distribution of pteridophytes supported **Rapoport's rule**… test of **Rapoport's rule** …

On the relationship between species diversity and range size

Q Guo, H Qian, J Zhang - Journal of Biogeography, 2022 - Wiley Online Library

… relationship thus has significant implications for species **invasion** biology and conservation. … examined **Rapoport's rule**, across either latitude or altitude. In such cases, **Rapoport's rule** …

Preliminary Estimation of the Influence of *Cydalima perspectalis* **Invasion** on the Species Composition and Structure of Earthworm Population (Oligochaeta: Lumbricidae …

IB **Rapoport**, AY Puzachenko, C Csuzdi… - Russian Journal of …, 2022 - Springer

The earthworm fauna and population structure in Colchic ecosystems of the southern slope in the Western Caucasus were studied. First, in May 2013 we have sampled earthworms of …

Plant invasions: merging the concepts of species invasiveness and community invasibility

DM Richardson, P Pyšek - Progress in physical geography, 2006 - journals.sagepub.com

… key issues in plant **invasion** ecology, where … in **invasion** ecology. Some organizing and unifying themes in the field are organism-focused and relate to species invasiveness (the tens **rule**…

**Range size distributions-Date: 04/04/2025.**

**Search Terms: Range size distribution* invasi***

Using scale–area curves to quantify the **distribution**, abundance and **range** expansion potential of an **invasive** species

R Veldtman, SL Chown… - … and **Distributions**, 2010 - Wiley Online Library

… of biological invasions and **invasive** plant **distributions**. Here, we detected potential areas of **invasive** concern, plus differences in abundance and **distribution** patterns, and associated …

Space to invade? Comparative **range** infilling and potential **range** of **invasive** and native plants

BA Bradley, R Early, CJB Sorte - Global Ecology and …, 2015 - Wiley Online Library

… smaller potential **ranges** for non-natives than natives. We compare the **distributions** of native,
endemic, alien and **invasive** plants to determine how the different **range** attributes of these …

Climate controls the **distribution** of a widespread **invasive** species: implications for future **range** expansion

WG McDowell, AJ Benson, JE Byers - Freshwater biology, 2014 - Wiley Online Library

… that can control **distributions** is critical, especially for **invasive** species that … **distribution** of the widespread **invasive** freshwater clam Corbicula fluminea, and we model its future **distribution** …

Scale-area curves: a tool for understanding the ecology and **distribution** of **invasive** tree species

JE Donaldson, DM Richardson, JRU Wilson - Biological invasions, 2014 - Springer

… for **invasive** species with flat scale-area curves at the current edge of their **invasive range**, … species that occupies a small proportion of its suitable **range**, and compare our findings to …

Implementing and interpreting local-scale **invasive** species **distribution** models

TJ Brummer, BD Maxwell, MD Higgs… - … and **Distributions**, 2013 - Wiley Online Library

Aim Use of local-scale non-native plant species ( NNS ) **distribution** models has the potential to decrease survey effort and improve population prioritization for management. We …

A biogeographical approach to plant invasions: the importance of studying exotics in their introduced *and* native **range**

JL Hierro, JL Maron, RM Callaway - Journal of ecology, 2005 - Wiley Online Library

… We also touch upon some issues in **invasion** ecology that do not require an explicit biogeographical perspective, but where new approaches might be beneficial, and highlight how an …

The geographic **range**: **size**, shape, boundaries, and internal structure

JH Brown, GC Stevens… - Annual review of ecology …, 1996 - annualreviews.org

… This work was oftenmotivated by practical concerns about what limited the **distribution** ofcommercially valuable plants (eg 56),**invasive** weeds and insect pests (eg 1,91), or potential …

**[PDF]** Plant **invasion** across space and time: factors affecting nonindigenous species success during four stages of **invasion**.

KA Theoharides, JS Dukes - New phytologist, 2007 - blogs.law.harvard.edu

… of **invasive** NIPS is controlled by a series of key processes or filters. These filters are common
to all **invasion** … that are introduced across a wide swath of the nonnative **range** may be …

Insights from modeling studies on how climate change affects **invasive** alien species geography

C Bellard, JM Jeschke, B Leroy… - Ecology and …, 2018 - Wiley Online Library

… in the geographic **ranges** of **invasive** alien species related to … as a driver of **invasive** alien species **distribution** in the future. … a **range** of terms related to IAS (taxonomic and **invasion** terms…

Progress in **invasion** biology: predicting invaders

CS Kolar, DM Lodge - Trends in ecology & evolution, 2001 - cell.com

… influence the probability of a bird species becoming established outside its native **range**, independently from the probability that a bird species will become **invasive**, is unknown. …

Five potential consequences of climate change for **invasive** species

JJ Hellmann, JE Byers, BG Bierwagen… - Conservation …, 2008 - Wiley Online Library

… climate change could alter **invasive** species impacts via changes in **range**, abundance, and … on **range sizes** in sections above on climatic constraints and **distribution** of existing **invasive** …

Release of **invasive** plants from fungal and viral pathogens

CE Mitchell, AG Power - Nature, 2003 - nature.com

… We tested whether this decrease could be explained by changes in plant geographic **range size** 11 , estimated by geographic prevalence. Pathogen species richness increased with …

**[BOOK]** **Invasion** biology

MA Davis - 2009 - books.google.com

… Thus, for purposes of this book, I have restricted the use of the word **invasion** to those **range** … **invasion**, I decided to confine the use of the term **invasive** in this book to nonnative species. …

Use of niche models in **invasive** species risk assessments

A Jiménez-Valverde, AT Peterson, J Soberón… - Biological …, 2011 - Springer

… **invasion** and thereby not in distributional equilibrium (Peterson 2005a), many **invasive** species seem to have reached some level of distributional equilibrium in their new **ranges**, …

Human papillomavirus type **distribution** in 30,848 **invasive** cervical cancers worldwide: Variation by geographical region, histological type and year of publication

N Li, S Franceschi, R Howell-Jones… - … journal of cancer, 2011 - Wiley Online Library

… (HPV) type **distribution** in **invasive** cervical cancer (ICC) can … of the contribution of a broad
**range** of HPV types to ICC in … cause of **invasive** cervical cancer (ICC),1 data on HPV type …

Something in the way you move: dispersal pathways affect **invasion** success

JRU Wilson, EE Dormontt, PJ Prentis, AJ Lowe… - Trends in ecology & …, 2009 - cell.com

… The first dispersal pathway we discuss is diffusion (Figure 1a) where, as described by the leading-edge model of **range** shifts, species **distributions** expand from the edge of their **range** …

*Drosophila suzukii* (Diptera: Drosophilidae): **Invasive** Pest of Ripening Soft Fruit Expanding its Geographic **Range** and Damage Potential

DB Walsh, MP Bolda, RE Goodhue… - Journal of Integrated …, 2011 - academic.oup.com

… Cooperative projects involving Western states and Canada are underway to address knowledge gaps and prepare recommendations for dealing with this **invasive** new pest. …

… of populations introduced from a genetically structured native **range** by approximate Bayesian computation: case study of the **invasive** ladybird Harmonia axyridis

E Lombaert, T Guillemaud, CE Thomas… - Molecular …, 2011 - Wiley Online Library

… sets to retrace the origin of biocontrol and **invasive** populations of H. axyridis, … **invasive** population in eastern North America, which has served as the bridgehead for worldwide **invasion** …

A synthesis of plant **invasion** effects on biodiversity across spatial scales

KI Powell, JM Chase, TM Knight - American journal of botany, 2011 - Wiley Online Library

… on the effects of **invasive** plants on the species richness of invaded communities across a **range** of spatial extents. We then discuss studies that consider the role of **invasive** plants on …

A meta-analysis of trait differences between **invasive** and non-**invasive** plant species

M Van Kleunen, E Weber, M Fischer - Ecology letters, 2010 - Wiley Online Library

… pair-wise trait differences of a total of 125 **invasive** and 196 non-**invasive** plant species in the **invasive range** of the **invasive** species. We tested whether invasiveness is associated with …

### Supplementary Figures

PRISMA 2020 flow diagrams for identification and reviewing of documents.

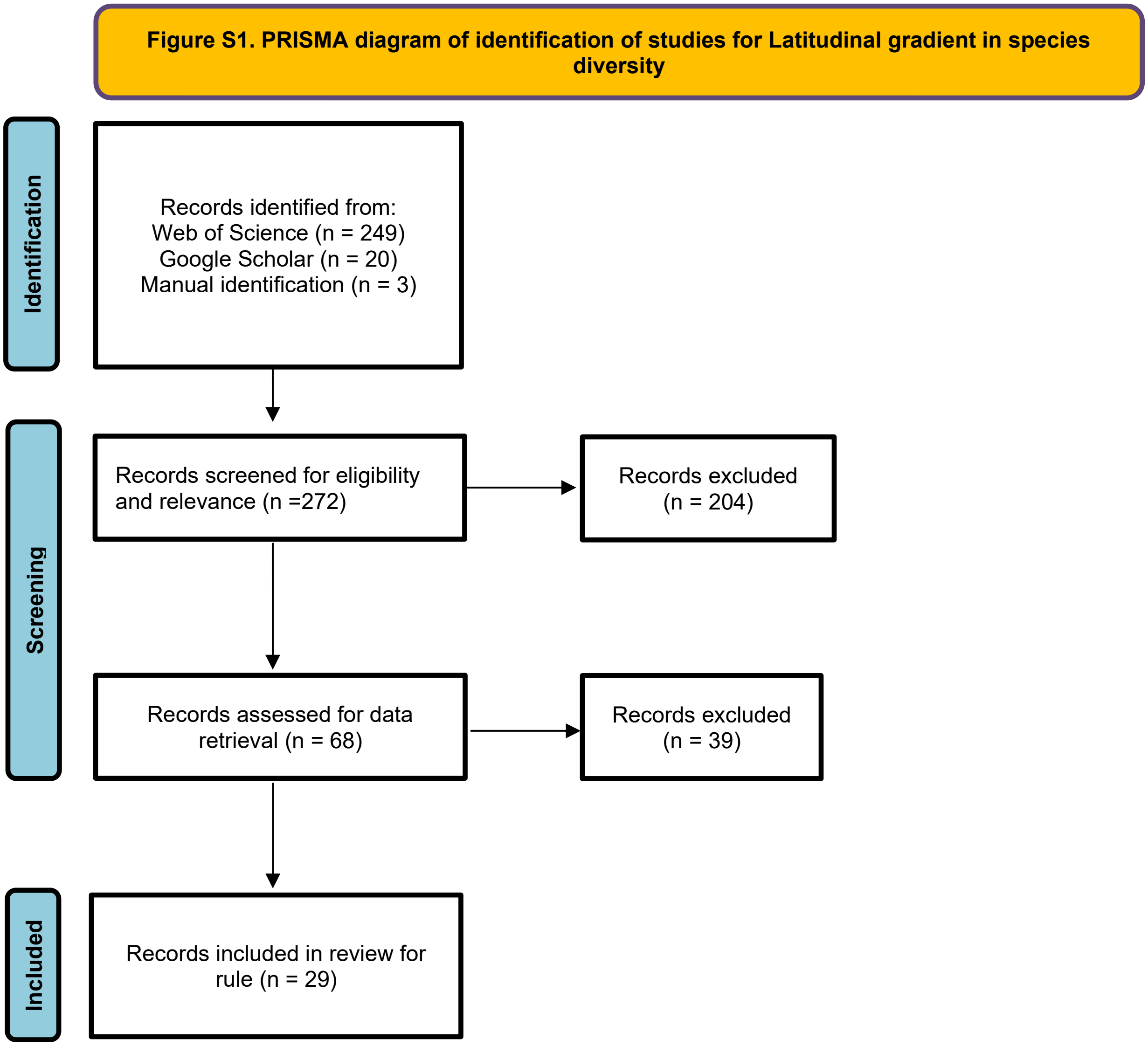

**Figure S2. PRISMA diagram of identification of studies for Biogeographical regions of the world**

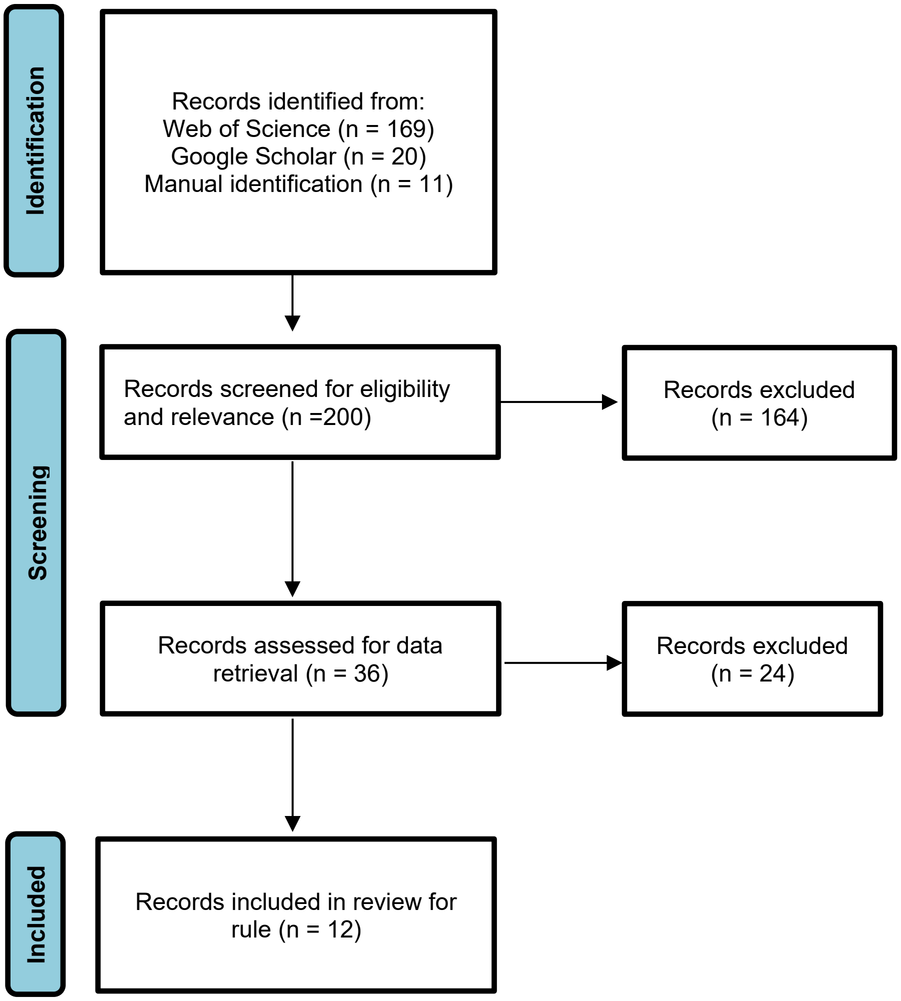

**Figure S3. PRISMA diagram of identification of studies for Wallace's Line**

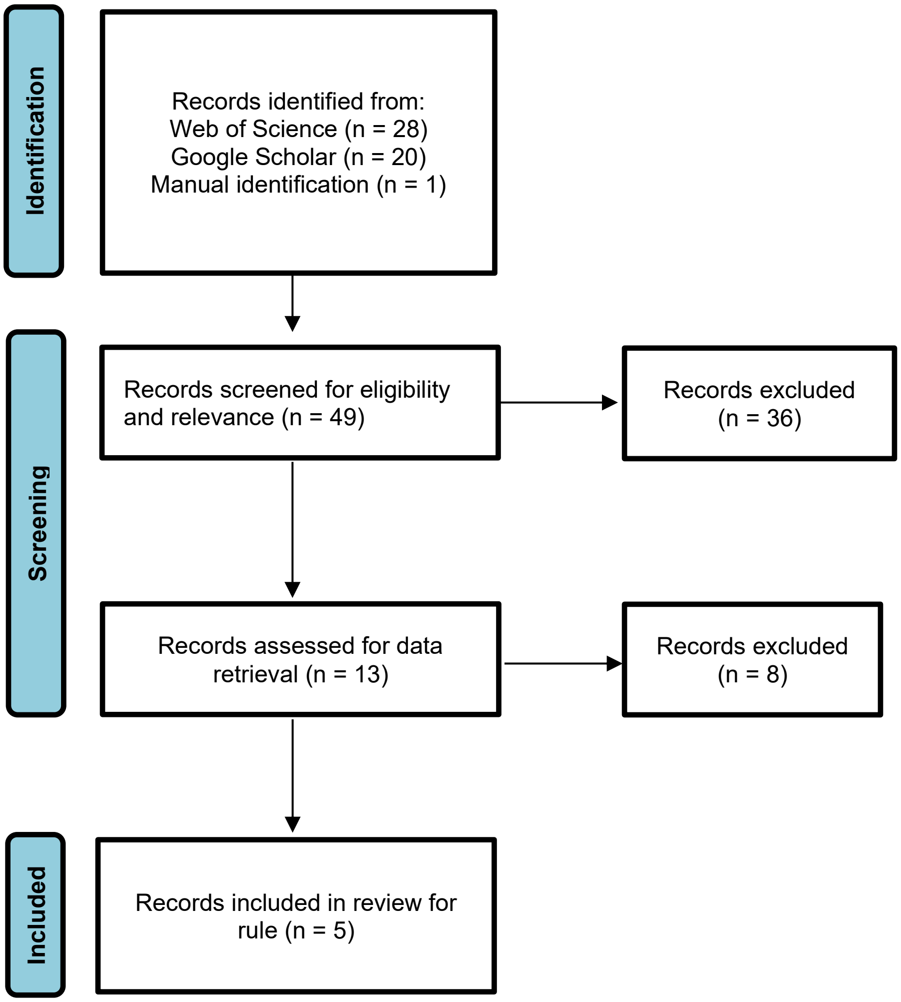

**Figure S4. PRISMA diagram of identification of studies for Species-area relationships and island species-area relationships**

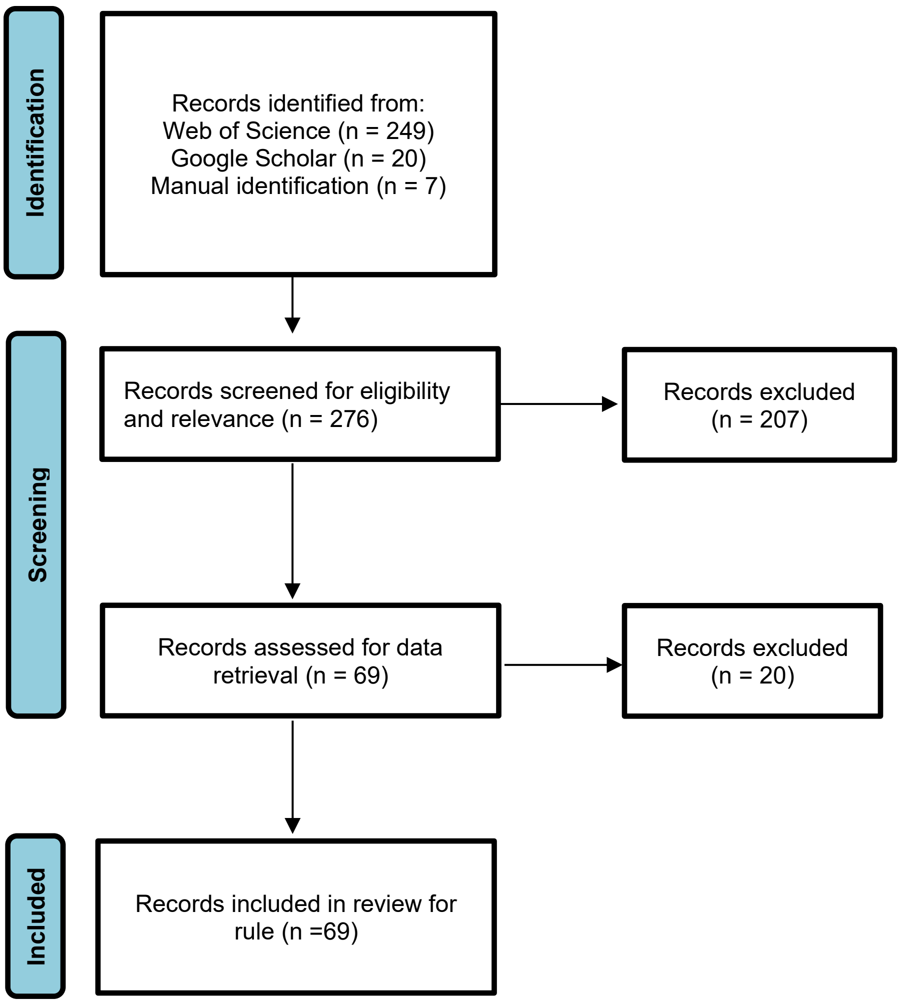

**Figure S5. PRISMA diagram of identification of studies for Island diversity-isolation**

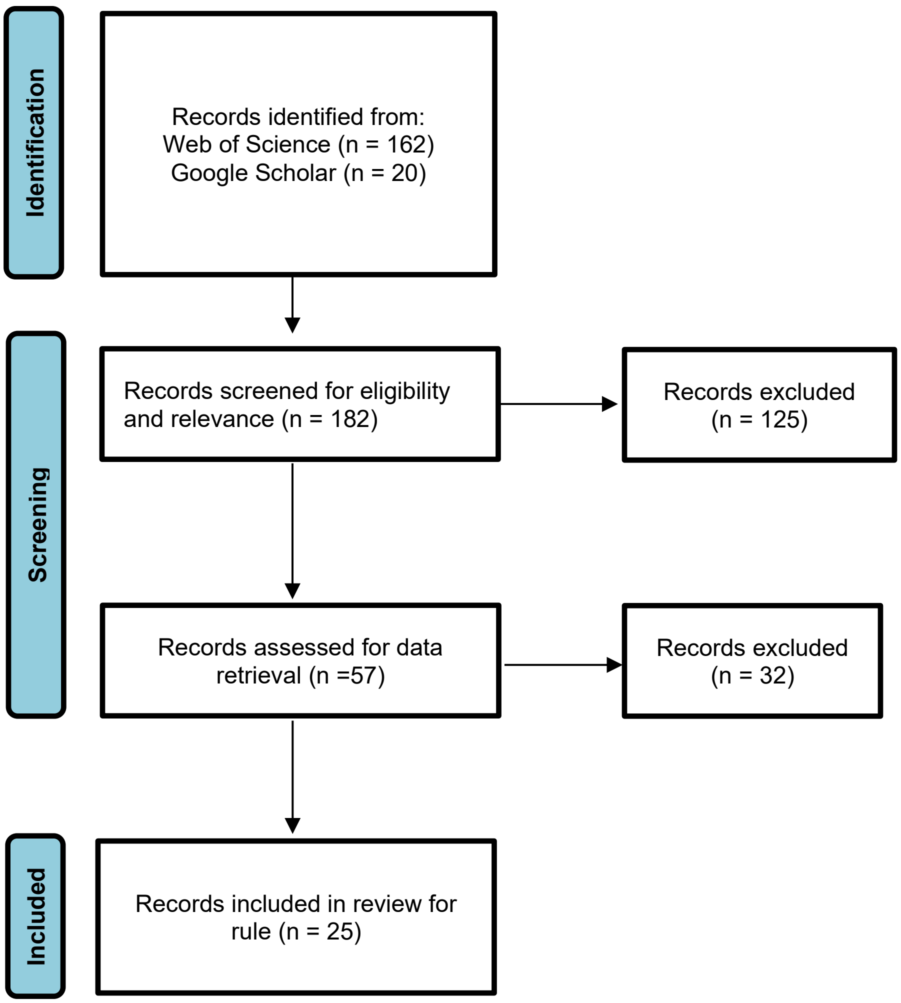

**Figure S6. PRISMA diagram of identification of studies for Distance decay of similarity**

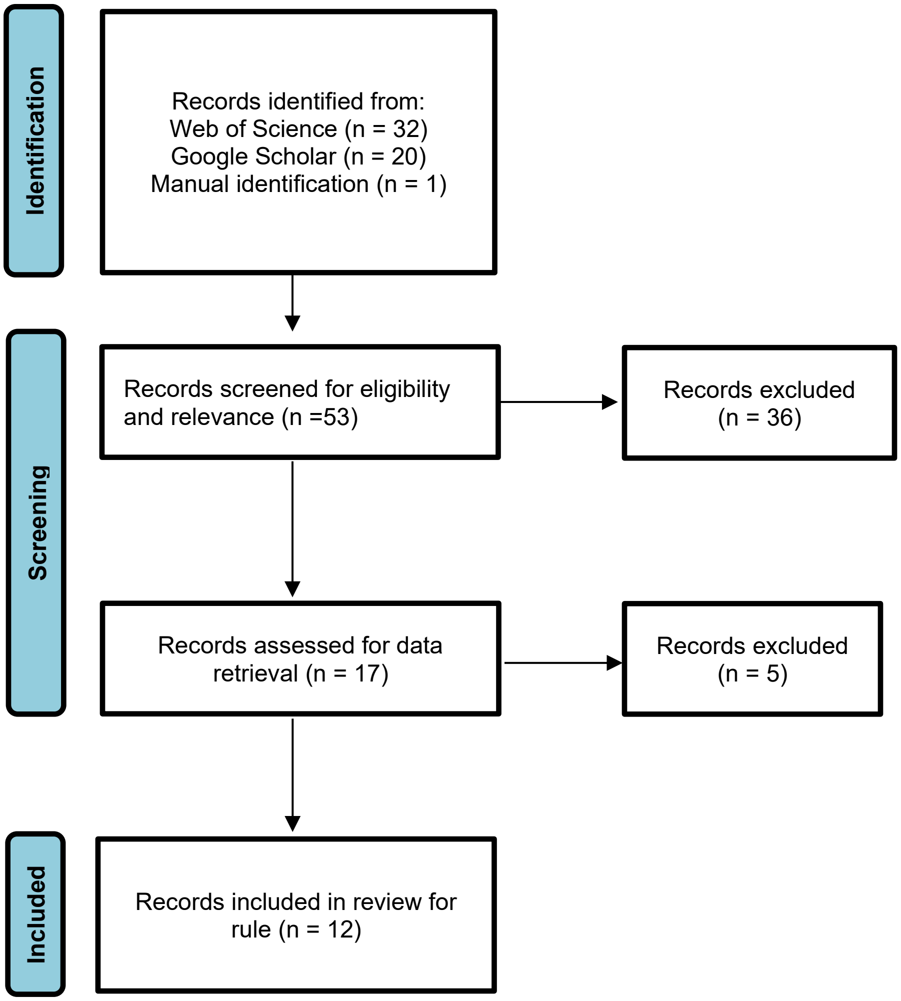

**Figure S7. PRISMA diagram of identification of studies for Bergmann's Rule**

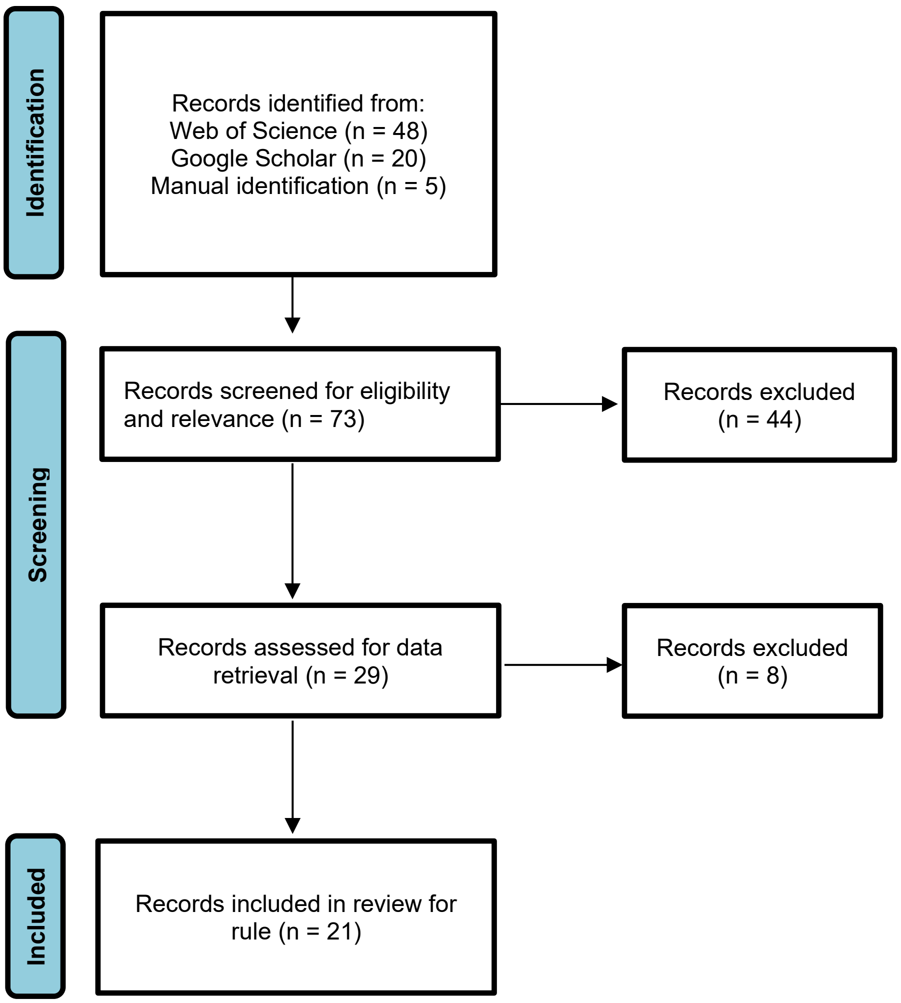

**Figure S8. PRISMA diagram of identification of studies for Rapoport's Rule**

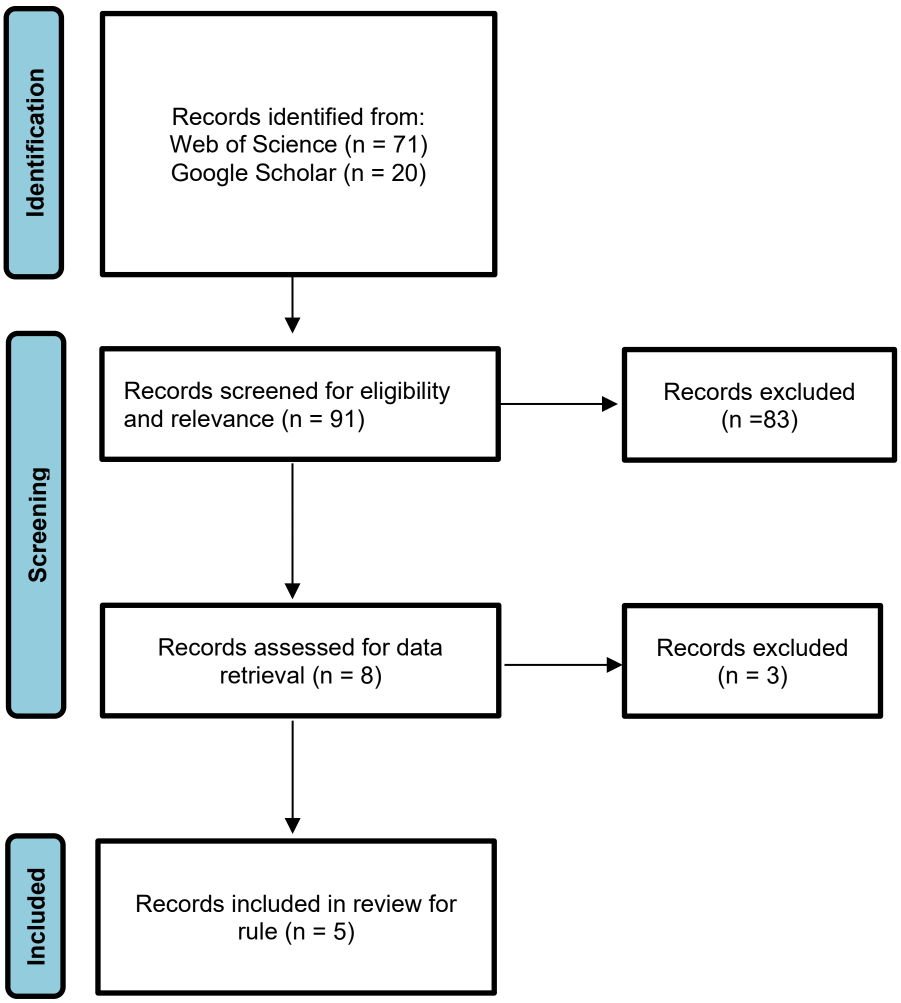

**Figure 9. PRISMA diagram of identification of studies for Gloger's Rule**

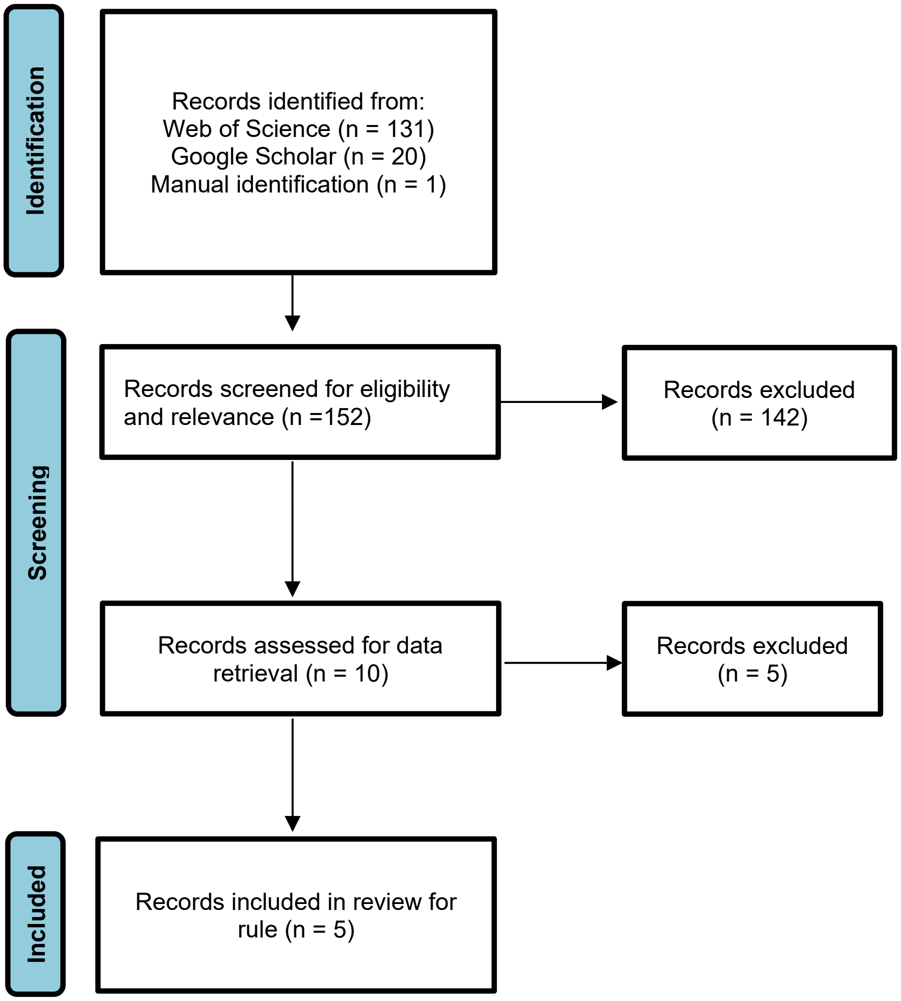

**Figure S10. PRISMA diagram of identification of studies for Foster's Rule (Island Rule)**

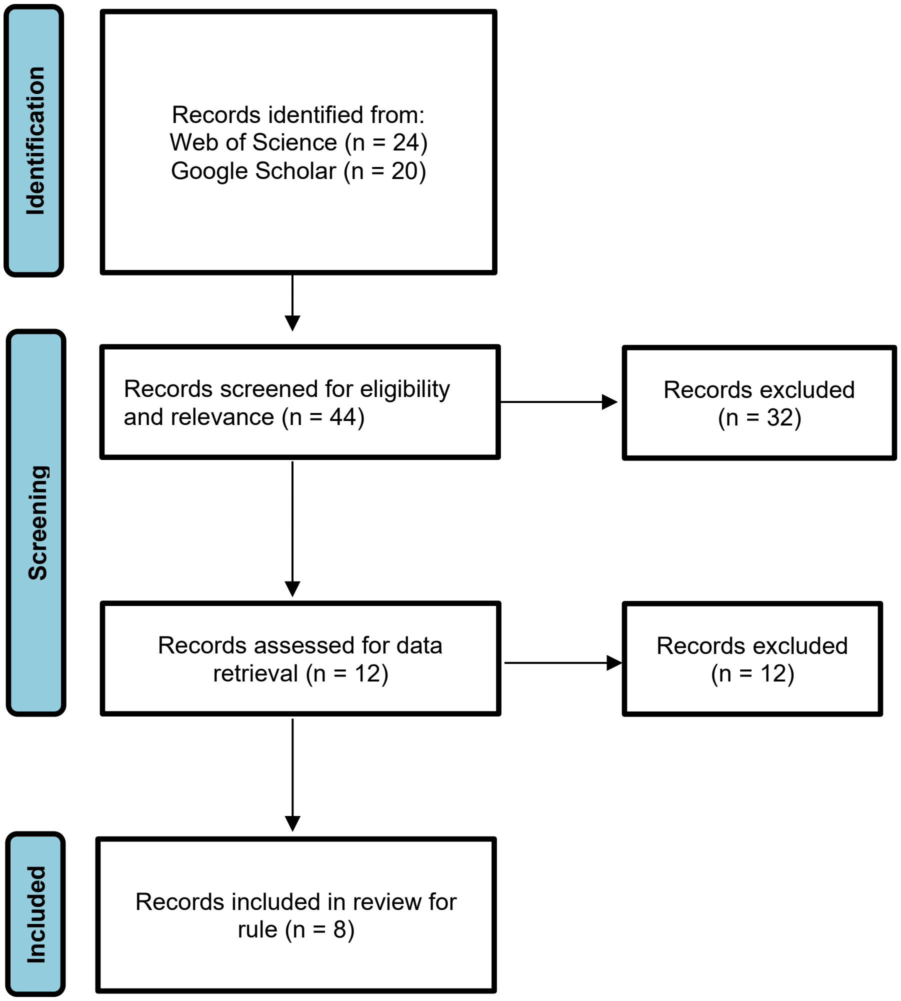

**Figure S11. PRISMA diagram of identification of studies for Body size-range size**

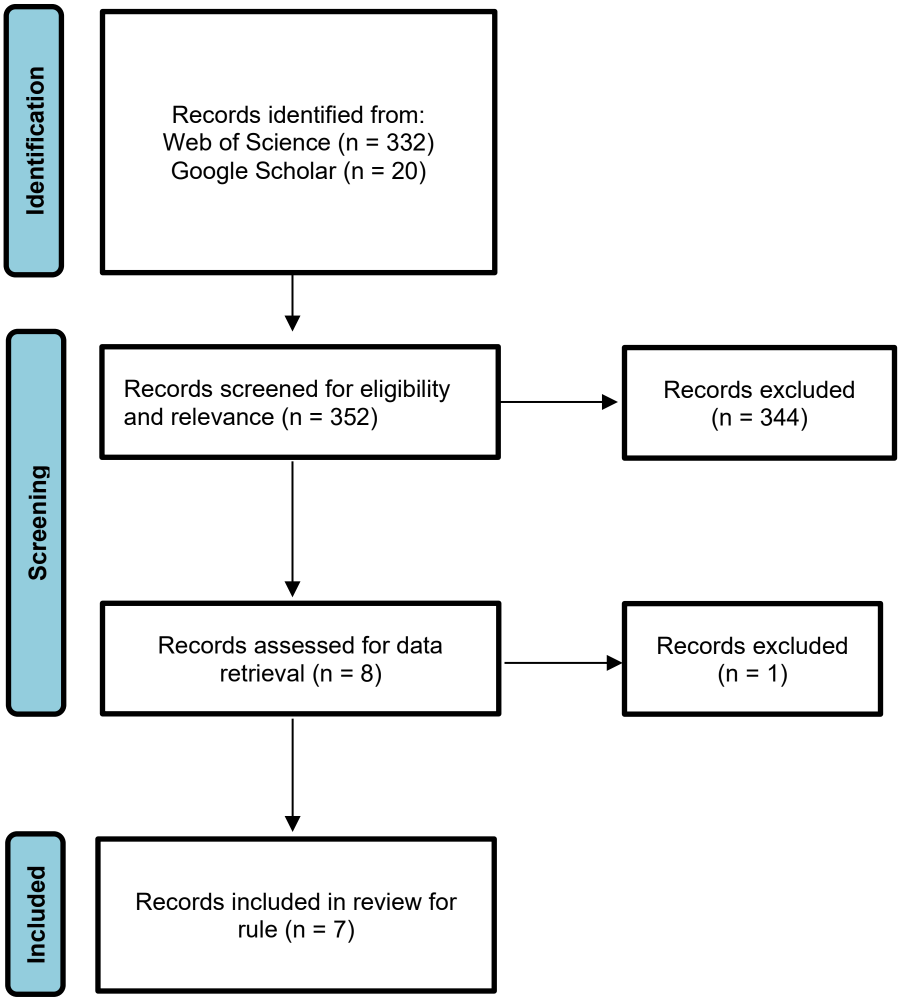

**Figure S12. PRISMA diagram of identification of studies for Abundance-range size**

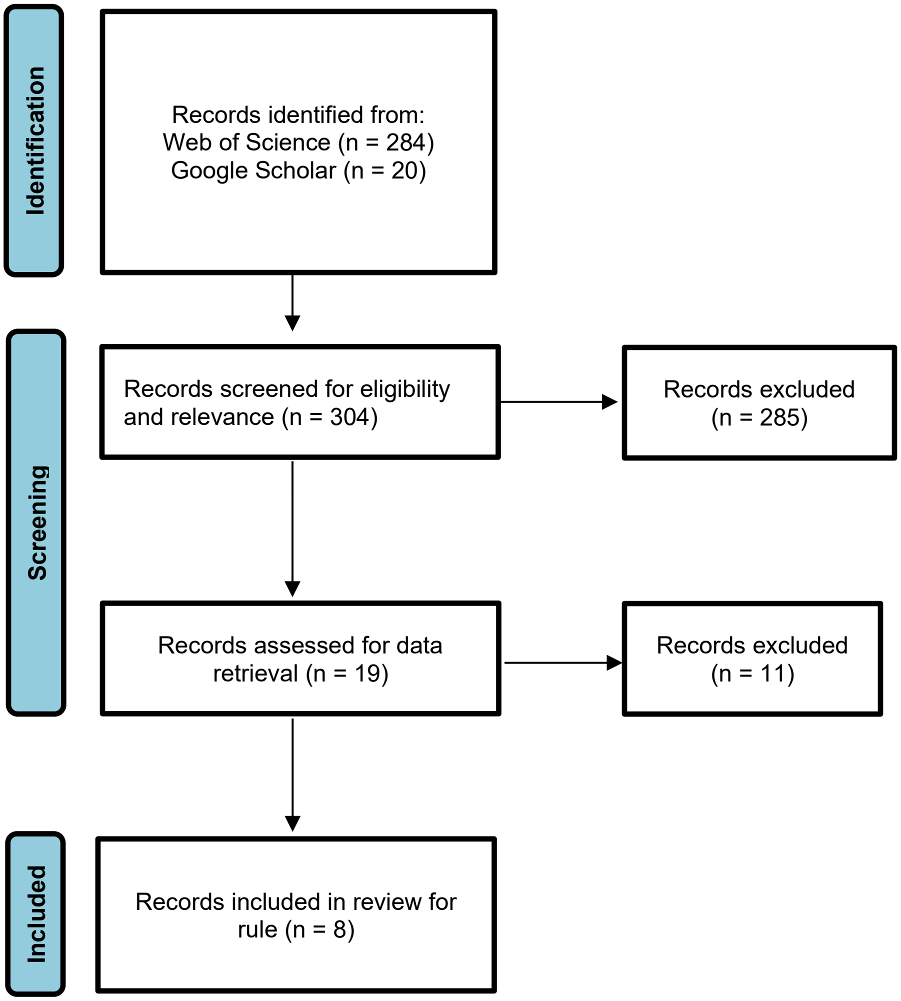

**Figure S13. PRISMA diagram of identification of studies for Species abundance distributions**

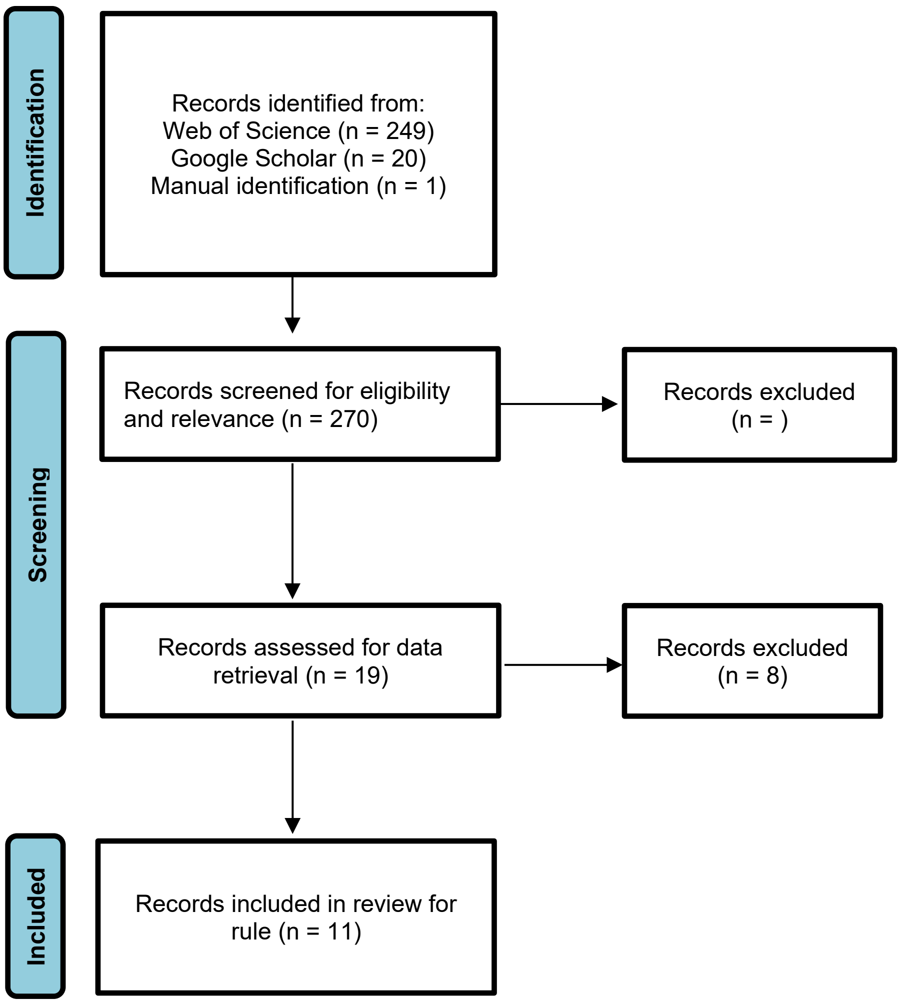

**Figure S14. PRISMA diagram of identification of studies for Body size distributions**

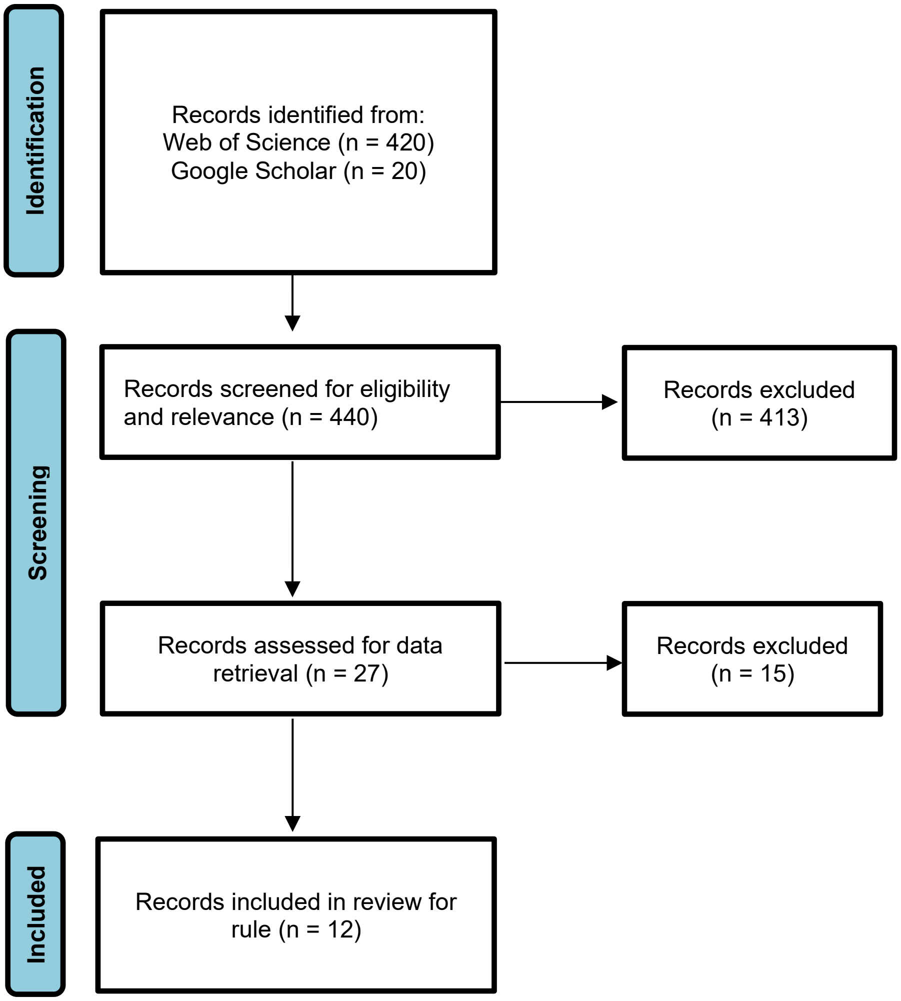

**Figure S15. PRISMA diagram of identification of studies for Range size distributions**

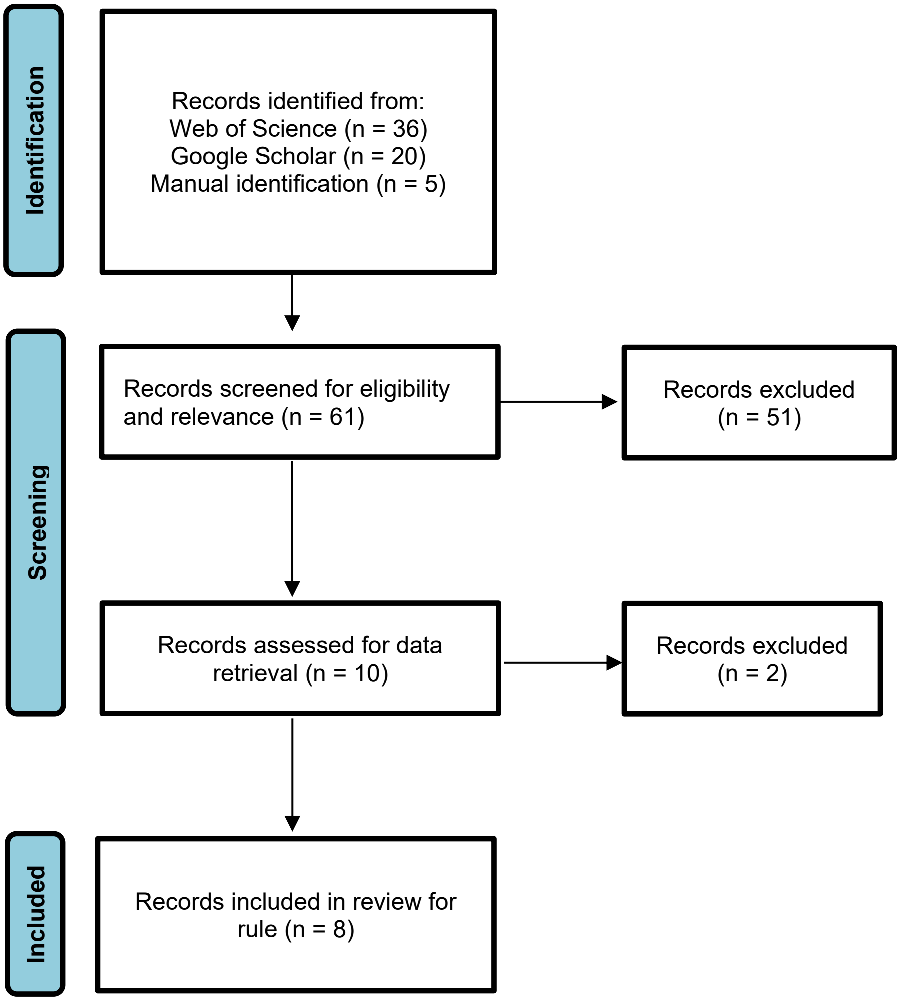
